# Optimal inference of molecular interactions in live FRET imaging

**DOI:** 10.1101/2022.03.29.486267

**Authors:** Keita Kamino, Nirag Kadakia, Kazuhiro Aoki, Thomas S. Shimizu, Thierry Emonet

## Abstract

Intensity-based live-cell fluorescence resonance energy transfer (FRET) imaging converts otherwise unobservable molecular interactions inside cells into fluorescence time-series signals. However, inferring the degree of molecular interactions from these observables is challenging, due to experimental complications such as spectral crosstalk, photobleaching, and measurement noise. Conventional methods solve this inverse problem through algebraic manipulations of the observables, but such manipulations inevitably accumulate measurement noise, limiting the scope of FRET analysis. Here, we introduce a Bayesian inference framework, B-FRET, which estimates molecular interactions from FRET data in a statistically optimal manner. B-FRET requires no additional measurements beyond those routinely conducted in standard 3-cube FRET imaging methods, and yet, by using the information contained in the data more efficiently, dramatically improves the signal-to-noise ratio (SNR). We validate B-FRET using simulated data, and then apply it to FRET data measured from single bacterial cells, a system with notoriously low SNR, to reveal signaling dynamics that are otherwise hidden in noise.

## Introduction

FRET is a short range (< 10 nm) effect whereby the energy of an excited fluorescence donor is transferred to an acceptor. By labeling proteins with the donor and acceptor, FRET transforms molecular interactions (i.e., protein-protein interactions for bimolecular FRET or protein conformational changes for unimolecular FRET) in live cells into fluorescence signals in real time^1–5^. The resulting fluorescence signals reflect the molecular states as time series of noisy observations, which must in turn be inverted to uncover the underlying molecular interactions within the cells. Thus, FRET analysis consists of two steps. First, information about molecular interactions is encoded into fluorescence signals (i.e., a fluorescence measurement). Second, data analysis is used to recover the information about molecular interactions from the measured fluorescence signals. The importance of optimizing the efficiency of encoding is well-recognized – various studies have emphasized fine-tuning FRET pairs^1,2,4–7^, as well as aspects of the microscope setup^8,9^, such as fluorescence filters and photodetectors, to maximize the photon budget. However, less attention has been paid to the efficiency of the data analysis used for decoding.

Conventional time-lapse FRET methods are based on sensitized emission from the acceptor, elicited by donor excitation, which encodes the information of molecular interaction into a few fluorescence time series. Decoding molecular information from these signals is complicated by spectral crosstalk^4,10,11^, photobleaching^12^ and measurement noise^13^. A range of decoding methods have been proposed^12,14–16^: some are qualitative (e.g., simple ratiometry^17^), often neglecting spectral crosstalk and photobleaching, while others are quantitative (e.g., E-FRET^12^), correcting both spectral crosstalk and photobleaching in a principled manner. Although different in assumptions, these methods decode the information of molecular interaction by algebraically processing the time-series data, and computing a “FRET index” as a measure of the degree of molecular interactions. However, algebraic manipulation of noise-corrupted data inevitably accumulates noise – for example, if a fluorescence intensity *I*_1_ is subtracted from another *I*_2_, the resulting intensity is smaller than *I*_2_ (i.e., *I*_2_ – < *I*_1_ < *I*_2_), but its noise measured by variance is larger than that of *I*_2_ (i.e., Var(*I*_2_ – *I*_1_) = Var(*I*_2_) + Var(*I*_1_) > Var(*I*_2_)). This lowers signal-to-noise ratio (SNR) of the computed FRET indices, and thus makes it more difficult to discern the dynamics of underlying molecular interaction. Such effects have limited the application of FRET methods to cases where photon budgets are large and SNR is inherently high.

Here, we develop a computational framework, B-FRET, to infer, from standard 3-cube FRET data^11,12^, the degree of molecular interactions defined by a FRET index *in a statistically optimal manner*. By applying the well-developed frameworks of Bayesian inference^18,19^ and filtering theory^20,21^, B-FRET systematically deals with the many confounding factors associated with the sensitized-emission-based FRET imaging methods – including the measurement noise – without algebraic manipulations. This enables B-FRET to maximally exploit the information from measured data, drastically improving the SNR of extracted FRET time series. Furthermore, B-FRET produces not just optimally estimated *values* of FRET signals, but their full probability *distributions*. Thus, B-FRET quantifies the statistical uncertainty of the estimation of molecular interactions at each time point, an aspect that is absent in previous algebra-based methods^16^. We use B-FRET to analyze noisy FRET data from live single bacterial cells, and show that it estimates FRET signals and hence cellular dynamics at an unprecedented level of precision.

## Results

### B-FRET framework and learning algorithm

A FRET sample under investigation contains fluorescent proteins whose states (fluorescent or photobleached, and free or complexed) change in time. A FRET measurement is a (noisy) map from a configuration of fluorescent proteins in various states to observable fluorescence signals (Fig. 1a). The goal of quantitative FRET data analysis is to infer, from the observables, the degree of donor-acceptor interactions in a way that is interpretable in terms of molecular interactions and independent of instrument-specific parameters and photobleaching. The degree of interaction can be defined in various ways depending on the purpose of an experiment, and we call such a user-defined degree of interaction a FRET index *E. B-FRET is a computational framework to infer the FRET index E in a statistically optimal manner*. For concreteness, we consider the most general case of bi-molecular FRET, in which the donor and acceptor are on different carrier molecules and hence the stoichiometry of acceptor to donor is not fixed, and compare results with those obtained using the widely used E-FRET method^12^ which corrects for both spectral crosstalk and photobleaching (Online Methods and Supplementary Note 3). Note, however, that B-FRET is not restricted to bi-molecular FRET; the same idea and algorithm can be used for uni-molecular FRET systems, where the donor and acceptor are attached to the same carrier, so that the conformational changes of the carrier molecule are encoded in fluorescence signals (Supplementary Note 1 and 2).

**Figure 1.**
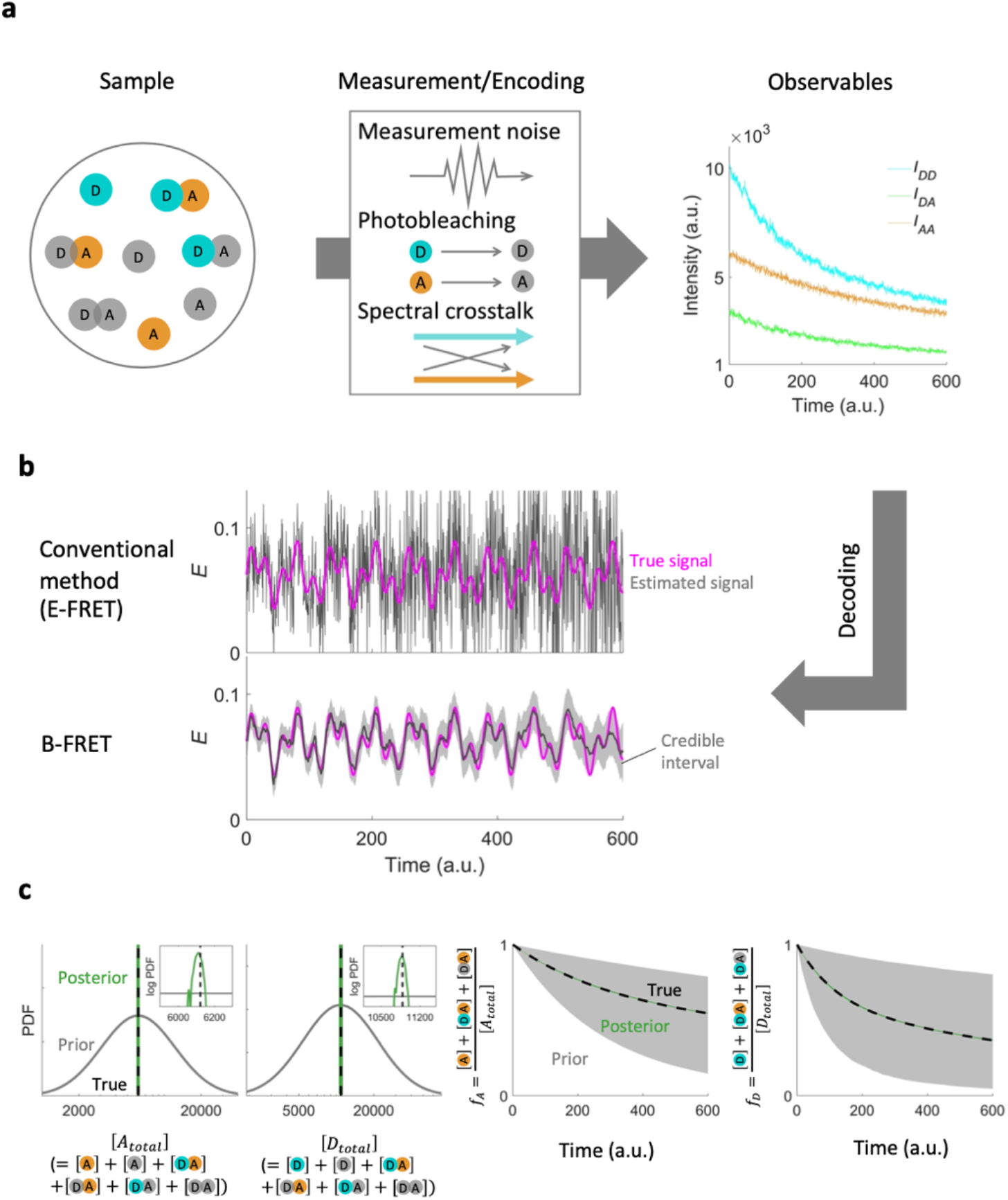
Optimal decoding of the FRET index *E* from FRET data. (a) A schematic of 3-cube time-lapse FRET imaging. For both bi-molecular and uni-molecular FRET systems, a FRET sample (left) consists of 8 chemical species (the schematic represents the case of bi-molecular FRET, where the donor and acceptor can be isolated from each other): fluorescent donor (D; cyan) and acceptor (A; orange), photobleached D and A (grey) and 4 different D-A complexes. A time-lapse FRET measurement encodes the molecular information into the three fluorescence time series (right), *I_AA_* (acceptor emissions during acceptor excitation), *I_DD_* (donor emission during donor excitations), and *I_DA_* (acceptor emission during donor excitations). The encoding subjects to measurement noise, photobleaching, and spectral crosstalk (middle). The relationship between the sample state and the observables can be expressed by a photophysical model with unknown parameters, which are learned from the observables via B-FRET. (b) Decoding the information of molecular interactions measured by the FRET index *E* from the synthetic data shown in a both by the E-FRET (top) and B-FRET (bottom). True (magenta) and estimated FRET index *E* (grey line) are shown. Implementing an optimal decoding, B-FRET estimates the true signal more precisely. For B-FRET, 95% credible intervals are shown by grey shade. (c) Prior and posterior distributions of the unknown parameters in the photophysical model. From left to right: the total concentrations of acceptor ([*A_total_*]), donor ([*D_total_*]), the fraction of intact acceptor (*f_A_*), and donor (*f_D_*). Insets in the left two panels are magnified posterior distributions (green) plotted in log scale for both X and Y axes. In the right two panels, ranges of priors and posteriors of *f_A_* and *f_D_* (from 2.5 to 97.5 percentile) are shown. In all cases, posteriors are highly confined, implying the presence of rich information about the model parameters in the observables shown in a.

In a bimolecular FRET system, the donor (D) and acceptor (A), fused to two different molecules, can form a molecular complex leading to FRET from the donor to the acceptor. The system is characterized by the concentrations of eight chemical species: D*, D, A*, A, D*A*, D*A, DA*, and DA, where fluorescent and non-fluorescent molecules are indicated by the presence and absence of a star (*) respectively, and donors and acceptors can be free (e.g., D*) or complexed (e.g., D*A*) (Fig. 1a). In general, inferring the temporal evolution of all the eight chemical concentrations from data of a smaller number of time series is ill-conditioned and intractable. Accordingly, all quantitative FRET analysis methods to date depend on simplifying assumptions (e.g., the conservation of the donor and acceptor concentrations during a measurement) that are satisfied in a standard experimental FRET setup. Consistent with this, we restrict the scope of this paper to a set of standard assumptions – specifically, that which underlies the E-FRET method^12^. See Online Methods and Supplementary Note 1 for the assumptions we make for both bi- and uni-molecular FRET. Note, however, that the B-FRET framework described below is generic and does not depend on any particular set of assumptions (see Discussions).

The applicability of B-FRET is also independent of the specific definition of *E* (see Discussions). For concreteness, here we consider a standard measure of the degree of interaction^12,14,22,23^:

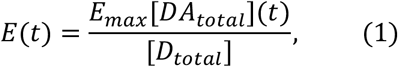

where [*D_total_*] ≡ [*D*\*](*t*) + [*D*](*t*) + [*D*\**A*\*](*t*) + [*D*\**A*](*t*) + [*DA*\*](*t*) + [*DA*](*t*) is the total concentration of the donor molecule, which we assume to be constant, and [*DA_total_*](*t*) ≡ [*D*\**A*\*](*t*) + [*D*\**A*](*t*) + [*DA*\*](*t*) + [*DA*](*t*) is the total concentration of the donor-acceptor complex, which can change over time; *E_max_* is the specific FRET efficiency of the complex, defined as the probability of energy transfer from the donor to acceptor in the donor-acceptor complex per donor excitation event and is constant given a FRET pair and an experimental condition^2,5^. The FRET index *E* defined by Eq. 1 is independent of instrument-specific parameters and of the degree of the photobleaching of the fluorescent molecules, and is linearly dependent on the fractional occupancy of the donor, making it an ideal measure of the degree of molecular interaction^12^.

At its core, B-FRET is a direct application of Bayesian inference for so-called *state space models*^20,21^. In this framework, one infers the temporal evolution of hidden (i.e., unobservable), dynamical state variables from noisy observations. A state space model consists of a *dynamic model*, which describes the temporal evolution of hidden state variables, and a *measurement model*, which is a static function mapping the hidden variables at time t to observables at time t. We discuss these in turn.

In B-FRET, the hidden dynamic variable is the product of the specifc FRET efficiency and the total concentration of the complex, i.e., *χ*(*t*) = *E_max_* [*DA_total_*](*t*) (Online Methods). The dynamic model links *χ*(*t*) at two consecutive times via a probability distribution called the *process noise* described by a parametric probability distribution such as a Gaussian distribution (Online Methods; Supplementary Note 2). The assumption of a dynamical model is a central feature of B-FRET: it allows us to exploit temporal correlations in the hidden variable over small times^20,21^, which algebraic methods such as E-FRET neglects. On the other hand, process noise introduces additional parameters, (e.g., the standard deviation of a Gaussian distribution). These are estimated as part of the B-FRET algorithm, as described below and in more detail in Online Methods and Supplementary Note 2.

In addition to the dynamic model, B-FRET requires a measurement model, which describes the photophysical processes by which the hidden dynamic variable *χ*(*t*) is converted into observables^13,24^. In the standard 3-cube FRET imaging setup, the data consists of three time series of fluorescence intensities *I_AA_, I_DD_*, and *I_DA_* (Fig. 1a). These are, respectively, fluorescence measured at the acceptor emission band during excitation of the acceptor band; fluorescence measured at the donor emission band during excitation of the donor band; and fluorescence measured at the acceptor emission band during excitation of the donor band. Other than excitations of and emissions from fluorescent proteins, the photophysical processes involved in a fluorescence measurement include photobleaching of fluorescent proteins, spectral crosstalk (i.e., bleed-through of donor emission to the acceptor emission band and cross-excitation of the acceptor at the donor excitation band), energy transfer from the donor to acceptor due to FRET, and measurement noise (Fig. 1a). All of these effects are incorporated into a single probabilistic model that linearly maps the hidden variables *χ*(*t*) into the three observables *I_AA_*(*t*), *I_DD_*(*t*), and *I_DA_*(*t*) – this is our measurement model (Online Methods and Supplementary Note 1). Like the dynamic model, the measurement model is another central feature of B-FRET: this model makes all the assumptions involved in the decoding process mathematically explicit, whereas they can be implicit or even undefined in algebraic methods. As with the process noise in the dynamical model, this linear map has unknown parameters, one of which is [*D_total_*] in the equation 1. We estimate them as described below and in more detail in Online Methods and Supplementary Note 2.

Given these two ingredients – the dynamical model and measurement model – our goal is to estimate the FRET index *E*(*t*) = *χ*(*t*)/[*D_total_*] (Eq. 1) from observables. In the framework of Bayesian inference^18,19^, this amounts to computing the posterior distribution of *E*(*t*), 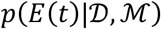, which quantitatively describes how well the possible values of *E*(*t*) are confined given all the data 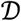 and the model 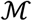. Since this distribution contains all the information one can theoretically have, computing the distribution ensures the statistically optimal inference of the FRET index. Because model parameters are also unknown, they must also be inferred from data. Thus, the computation of the posterior distribution of the FRET index is decomposed into the evaluations of two distributions: the posterior distribution of *E*(*t*) *given* specific model parameter values, and the posterior distribution of the model parameters themselves (Online Methods). These two distributions can be evaluated using Bayesian smoothing and filtering theory, respectively (Supplementary Note 2). Once these distributions are determined, the posterior distribution 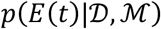 is computed using a Monte Carlo approach (Online Methods).

### B-FRET efficiently learns from data

To see how much the B-FRET algorithm improves the SNR of the estimated FRET index, we compared the FRET index *E* computed by B-FRET with that computed by the conventional E-FRET method. We first generated a synthetic (bimolecular) FRET data set by simulating oscillatory dynamics of FRET signals and all the confounding factors present in real data, namely spectral crosstalk, photobleaching and measurement noise (Fig 1a; Supplementary Note 4). With relatively large measurement noise, the oscillatory FRET dynamics are hard to see in the raw time-series data (Fig. 1a). Consequently, the FRET index computed by E-FRET is highly noisy, and the true oscillatory dynamics are obscured (Fig. 1b). However, we found that the FRET index computed by B-FRET estimates the true signal substantially more precisely, as evidenced by comparison of estimation errors (Fig. 2b). Furthermore, unlike E-FRET, B-FRET naturally provides statistical uncertainty of the estimated FRET index as a credible interval (CI) at each time point. As expected, the width of 95% CIs (Online Methods for definition) increases over time because of the decreasing data quality resulting from photobleaching (grey shadow in Fig. 1b). Consistent with precise estimation of *E*, the posterior distributions of the model parameters are highly confined around their true values used to generate the synthetic data (Fig. 1c). These observations demonstrate that the raw fluorescence time-series data, despite high levels of noise, contain rich information about molecular interactions, and B-FRET successfully exploits the available information to better constrain the possible values of the FRET index and model parameters.

**Figure 2.**
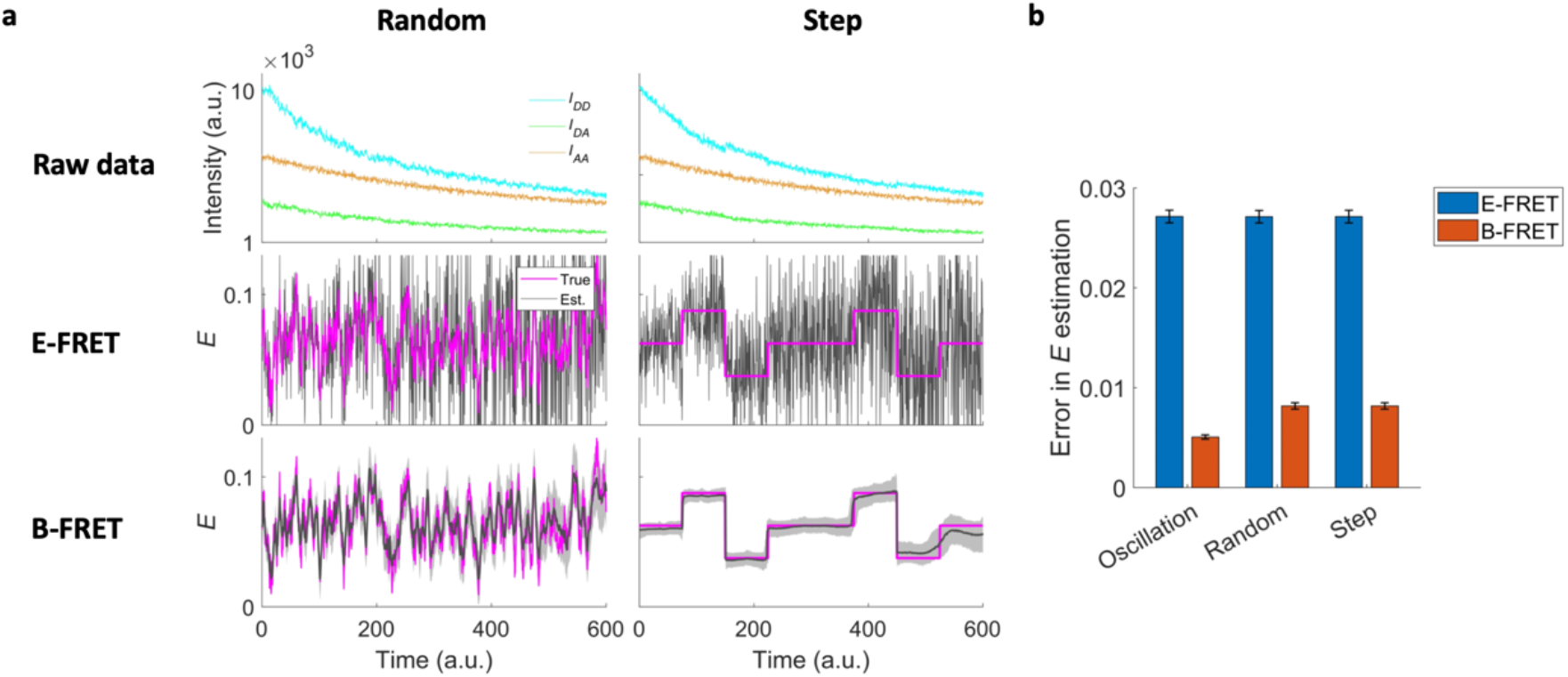
B-FRET performance is robust for different FRET dynamics. (**a**) Synthetic data with random (left) and step (right) FRET signals. (Top) Raw fluorescence timeseries. (Middle) The FRET index *E* computed by E-FRET (grey; *E_est_*) and its true values (magenta; *E_true_*). (Bottom) The FRET index *E* computed by B-FRET (grey; *E_est_*) and its true values (magenta). The shade shows 95% credible interval. (**c**) A bar chart quantifying the error in *E* estimation defined as 〈|*E_est_* – *E_true_*|〉, where the angle bracket is temporal average. The error bars are standard deviation over 5 data sets with identical FRET signal dynamics but different realizations of measurement noise.

### B-FRET is robust to the variation in FRET temporal patterns

To see how much the precision of FRET-index estimation is affected by the underlying temporal pattern of FRET signals, we next generated synthetic data in which the FRET signal exhibits random dynamics (Supplementary Note 4). Unlike the case of oscillatory dynamics (Fig. 1), the random signal is aperiodic and contains a broad range of frequencies, including those comparable to or higher than the data-sampling frequency, which precludes algorithms that exploit regular patterns in a signal. Despite this, we found that the FRET index computed by B-FRET is more precise and less noisy than that computed by E-FRET (Fig. 2a and Supplementary Fig.1).

The above two cases, oscillatory and random, were successfully analyzed with a Gaussian process noise, a standard choice for the process noise for its flexibility in capturing a broad class of dynamics^20,21^. However, for highly non-Gaussian dynamics, e.g., ones that remains unchanged most of the time but exhibit abrupt step changes only occasionally, it is known that non-Gaussian process noise can perform better than Gaussian process noise^21^. Although B-FRET is computationally cheaper with Gaussian process noise since many calculations can be executed analytically, the algorithm can be adapted to other process noise statistics by replacing the analytical calculations with numerical ones (Supplementary Note 2). To test the performance of B-FRET for non-Gaussian dynamics, we generated a synthetic FRET signal consisting of discrete steps (Supplementary Note 4), and modeled the process noise using a Student’s *t*-distribution (Online Methods). Indeed, the FRET index computed by B-FRET precisely captures the dynamics.

We note that B-FERT, combined with the framework of model selection, does not require a user to know in advance which model (e.g., a Gaussian or non-Gaussian process noise) to use to analyze a set of data. By computing the Bayes Information Criterion (BIC; Online Methods), B-FRET enables a user to automatically select a model that is best evidenced by the data. Applying this, we confirmed that the step data supports the choice of Non-Gaussian process noise, while the oscillatory and random data do not (Supplementary Fig. 2)

### B-FRET outperforms conventional methods irrespective of the measurement conditions

To see how the relative performance of B-FRET to E-FRET depends on the specific conditions of time-lapse imaging, such as the levels of measurement noise and sampling intervals, we investigated the signal-estimation errors of both methods for various measurement conditions. We first generated sets of synthetic data in which the degree of donor-acceptor interaction follows Gaussian random statistics over time with a correlation time *τ_c_* and fluorescence signals *I_AA_, I_DD_*, and *I_DA_* were measured with many different sampling frequencies *τ_c_*/Δ*t* and levels of measurement noise (Supplementary Note 4). For each data set, we then estimated the FRET index *E* using both B-FRET and E-FRET methods.

Fig. 3a shows representative results for B-FRET. As the SNR of raw fluorescent signals increases, B-FRET detects more subtle changes in *E* with lower statistical uncertainties (95% CIs are shown by grey shades in Fig. 3a). Also, the higher the sampling frequency relative to the FRET dynamics, the more precise the B-FRET estimation. This is because faster sampling increases the correlation between successive fluorescence signals, and these correlations are exploited by B-FRET. Meanwhile, the effect of reduced sampling frequency (Fig. 3a; lower left) can be compensated by increasing the data SNR (Fig. 3a; upper left).

**Figure 3.**
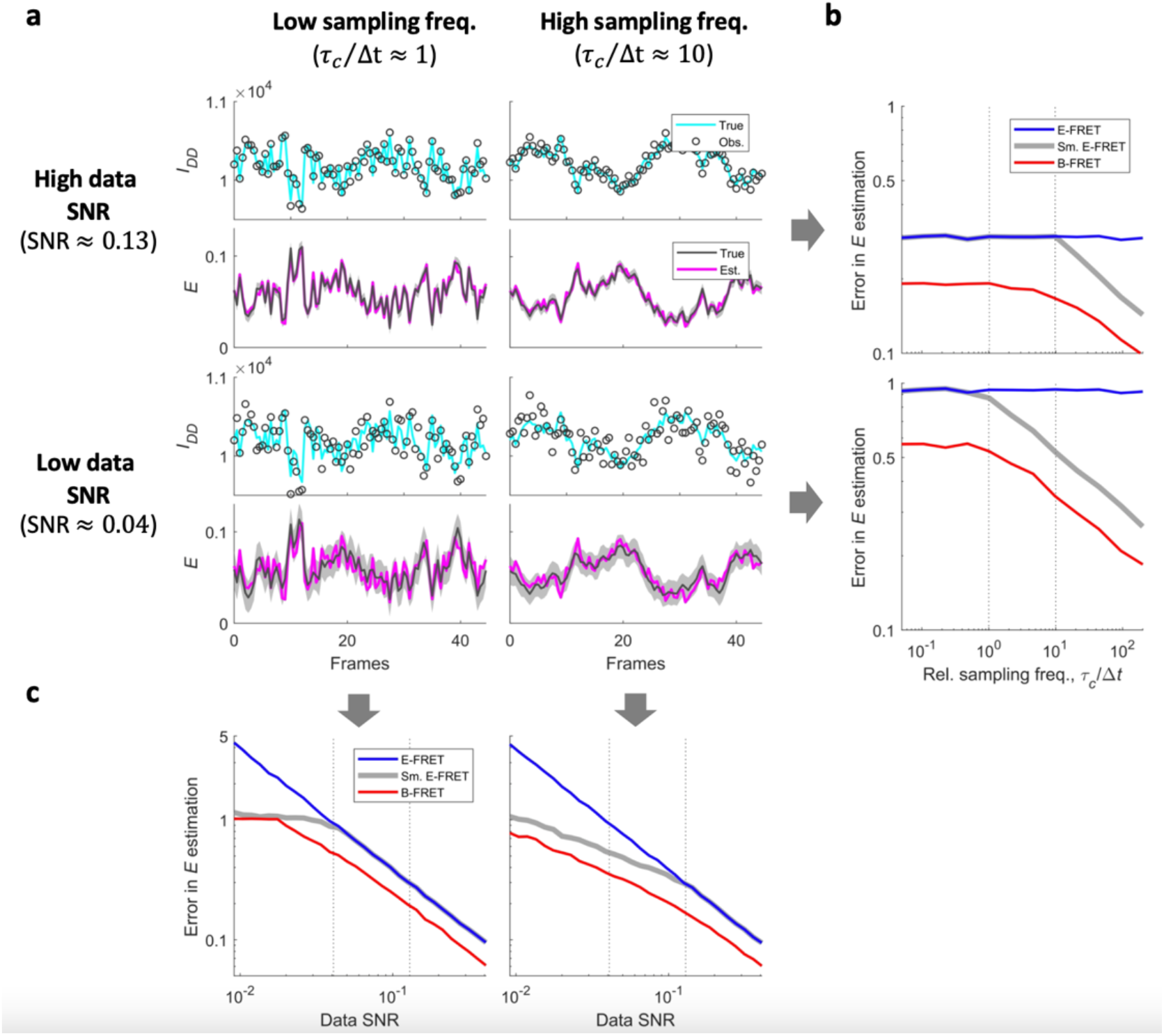
B-FRET outperforms conventional methods irrespective of measurement conditions. (**a**) Representative simulated data – *I_DD_* with (dark grey) and without (cyan) measurement noise – and estimated (dark grey) and true (magenta) values of the FRET index *E* in 4 measurement conditions are shown. FRET signals were changed randomly with a correlation time *τ_c_*, and data were sampled every Δ*t*; two sampling frequencies *τ_c_*/Δ*t* ≈ 1 (under-sampling regime) and 10 (over-sampling regime) are shown. Different levels of measurement noise were simulated without changing the expected values of the observables; two different data SNRs (Supplementary Note 4) are shown. (**b**) Errors in *E* estimation defined as 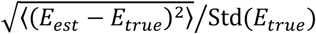 for B-FRET (red), E-FRET (blue) and E-FRET combined with optimal median filtering (grey) were plotted against sampling frequency, *τ_c_*/Δ*t*. Cases of high data SNR (≈ 0.13; top) and low data SNR (≈ 0.04; bottom) are shown. Optimal median filtering requires knowledge about true signals, which is not accessible, and hence cannot be implemented in practice. Thus, the grey line gives the minimum achievable error by the combination of E-FRET and median filtering. B-FRET requires no knowledge about true signals, and yet outperforms E-FRET in all explored conditions. (**c**) Error in *E* estimation plotted against data SNR. Cases of under-sampling (*τ_c_*/Δ*t* ≈ 1) and over-sampling (*τ_c_*/Δ*t* ≈ 10) are shown. Again, E-FRET outperforms E-FRET in all explored conditions.

We quantified the average error in FRET signal estimation as the root-mean-square error normalized by the magnitude of the fluctuation of the true signal, 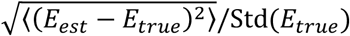, for both E-FRET and B-FRET in various measurement conditions (Fig. 3b and c). B-FRET (red) outperforms E-FRET (blue) in all conditions explored; importantly, even if E-FRET signals are smoothed with median filters with an optimal window size in terms of error reduction (grey) – which requires knowledge of the true FRET signal that an experimenter does not usually have access to – E-FRET still significantly underperforms compared to B-FRET. This can be understood by noting that, in the E-FRET method, some information about the true FRET signals contained in the raw fluorescent time series is already lost upon the algebraic computation to obtain *E*, and no degree of smoothing after that computation can recover the lost information. B-FRET, on the other hand, exploits the larger amount of information in the raw observables, including temporal correlations, and achieves more precise estimation of *E* without requiring any knowledge about FRET dynamics.

### B-FRET improves signal estimation of real data

To test the performance of B-FRET on real data, we applied the method to a previously developed bi-molecular FRET system that reports the kinase activity of the *E. coli* chemotaxis signaling pathway^22,23,25^. Recent FRET analyses of this pathway at the single-cell level have revealed fundamental features of cell signaling that are inaccessible by a population-level assay, such as spontaneous fluctuation in the pathway activity^25^, environment-dependent dynamic modulation of the degree of cell-to-cell variability^26^, and the high efficiency with which cells use information acquired by the pathway^27^. However, the FRET data from single *E. coli* cells are noisy: this is firstly because the small size of bacterial cells limits the number of fluorescent molecules per cell volume, and increasing the illumination power induces more photobleaching and phototoxicity. This has limited further characterizations of the signaling pathway.

The *E. coli* chemotaxis signaling pathway is a two-component signal transduction system^28^, where the receptor associated kinase CheA phosphorylates the response regulator CheY, which is then dephosphorylated by the phosphatase CheZ. Binding of chemoattractant molecules to the receptors changes the propensity for the receptor, and hence the kinase, to be active. Opposing this propensity is feedback regulation by methylation and demethylation enzymes. These two mechanisms together produce a steady-state kinase activity that is independent of the background chemoattractant concentration, a ubiquitous phenomenon in cell signaling called response adaptation^29–32^. The activity of the pathway can be read out by quantifying the FRET between the donor (mYFP) fused to CheZ and the acceptor (mRFP) fused to CheY, which binds to CheZ when phosphorylated by CheA. It has been well established that, upon a step increase in a chemoattractant concentration, the kinase activity and the concentration of phosphorylated CheY (and hence the level of FRET) decreases rapidly before response adaptation, while a step decrease in a chemoattractant concentration causes the opposite response^22,25^.

We measured fluorescence signals, *I_AA_, I_DA_*, and *I_DD_* from single *E. coli* cells using a 3-cube FRET measurement setup (Fig. 4a; Online Methods). In this setup, we delivered fast-switching (~ 0.1 s) step-like changes of *α*-methyl-aspartate (MeAsp), a non-metabolizable analog of the chemoattractant aspartate, using a recently-developed microfluidic system^26,27^. Large step changes in MeAsp (100% changes or higher) were delivered to cells at the beginning and end of the measurement to define the dynamic range (i.e., minimal and maximal FRET levels) of each cell. Several small step changes in MeAsp (20% changes) that cause sub-saturating responses, on average, were also applied in the middle of the measurement (Fig. 4b). First, we extracted the FRET index *E* using the E-FRET method (Fig. 4b left). As expected, the large noise prevented us from discerning single responses to the sub-saturating (20%) step stimuli. Quantifying responses from such noisy data requires some form of data averaging, as was done before^26,27^; however it unavoidably masks properties of individual responses. Next, we analyzed the same set of data using the B-FRET method (Fig. 4b right). B-FRET drastically improved the SNR and disclosed the cell-to-cell and temporal variations in the signaling dynamics more vividly: some cells respond to small step signals faithfully, whereas other cells neglect the same signals; some cells fluctuate vigorously, whereas some cells are more stable. Such variations could be functionally important for a cell population to deal with environmental uncertainties as recent studies have suggested^33–35^. Furthermore, B-FRET not just make some subsaturating responses clearly discernible by eye; it also enables us to tell whether the changes in FRET are statistically significant or not (red boxes in Fig. 4b). Finally, as with synthetic data (Fig. 1c), the posterior distributions of the model parameters are highly confined (Fig. 4c), demonstrating that real experimental data also contain sufficient information to confine the photophysical model. Together, these results demonstrate that B-FRET can greatly improve the quality of extracted FRET signals, and therefore help experimenters reveal novel dynamic features of cellular processes.

**Figure 4.**
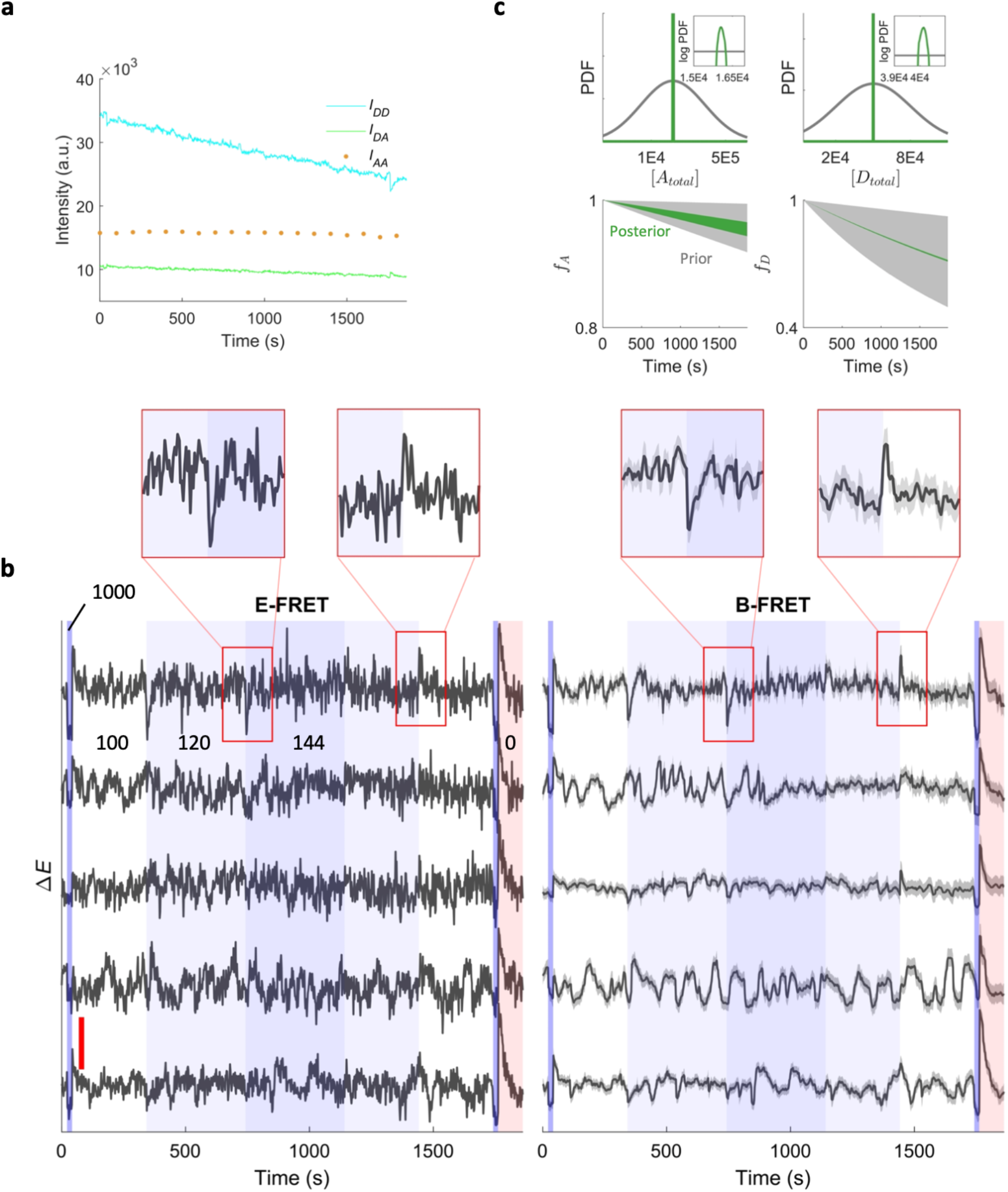
B-FRET infers the FRET index precisely in real FRET data. (**a**) Three observables, *I_AA_, I_DD_*, and *I_DA_*, were acquired from a bimolecular FRET system in single *E. coli* cells. The change s in FRET reports the changes in the activity of a kinase that governs chemotaxis behavior of *E. coli*. (**b**) Using FRET data sets from individual *E. coli* cells, the FRET index *E* was estimated by E-FRET (left) and B-FRET (right). Five representative cells are shown. Blues and red in the background indicate different concentrations (numbers in the unit of μM) of a chemoattractant MeAsp delivered to the cells. The red vertical line in the left panel corresponds to the change in the FRET index Δ*E* = 0.05. For B-FRET, 95% credible intervals are shown by the grey shade. The noise reduction by B-FRET reveals temporal and cell-to-cell variation in the FRET dynamics, while they are mostly obscured in the noise in the E-FRET results. Regions enclosed by the red boxes are expanded above. (**c**) Prior and posterior distributions of the model parameters. Despite the relatively high noise of the FRET data, the posterior distributions are highly confined, suggesting the efficient usage of information contained in the raw data by B-FRET. The data in panel a and c are from the cell shown at the bottom in panel **b**.

## Discussions

Inefficient decoding of the information about molecular interactions from FRET data amounts to wasting acquired photons. Here, we propose a computational framework, B-FRET, to decode the FRET index time series with theoretically maximal efficiency. A conventional way to improve SNR in live FRET imaging has been to aggregate signals from many samples (e.g., cells) and compute their average^22,23^; however, this method fails to capture variations and asynchronous dynamics across samples. B-FRET reduces the need for such averaging, as we demonstrated here by analyzing signaling dynamics in single bacterial cells (Fig. 4), and thereby providing a powerful aid to studies of biological variation – both across cells within a population, and across time within a single cell – that would be lost in averaging. B-FRET is of practical use even to experimenters who do not necessarily need to reduce SNR: to achieve a given SNR, B-FRET requires fewer photons, reducing the need for high-power illumination and therefore the unwanted effects of photobleaching and phototoxicity. Thus, B-FRET *computationally* extends the scope of FRET analyses by increasing the SNR and/or by requiring less photons, in much the same way as brighter FRET pairs or more sensitive photodetectors *experimentally* enhance FRET.

Although a method has been developed that systematically incorporates measurement noise and infers molecular interactions for *snapshot* FRET data^13^, developing a similar method for *time-lapse* FRET data has remained a challenge due to additional complications, such as photobleaching and temporal changes in the degree of FRET. Past approaches to the analysis of time-lapse FRET data have neglected measurement noise. B-FRET, by performing Bayesian inference on state space models, deals with measurement noise and other confounding factors in a principled manner. Furthermore, the statistical uncertainties of the observables are systematically converted to that of the inferred FRET index, which is expressed as the (posterior) distribution of the FRET index. This enables the experimenter to assess the uncertainty of the estimate and evaluate whether a change in a FRET index is statistically significant or not.

Another important advantage of B-FRET is that it decouples the assumptions involved in the inference problem from the definition of the FRET index, whereas these are inherently coupled in the past algebraic approaches. For example, E-FRET, under a certain set of assumptions, gives an algebraic formula (Online Methods) to compute the FRET index, which can be interpreted in molecular terms as 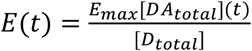 (Eq. 1). But the method gives no clue about how to compute different FRET indexes such as e.g. 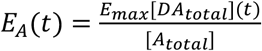. B-FRET provides a single algorithm for computing *any* FRET index, once it is defined. In B-FRET, therefore, inferring *E_A_*(*t*) is as straightforward as *E*(*t*), once the necessary parameters are provided (See Supplementary Note 1). In principle B-FRET can incorporate any set of assumptions into the inference procedure: different assumptions constrain the relationship between variables differently, but the core inference algorithm in B-FRET is independent of the constraints (Online Methods). However, this flexibility should not lead to the presumption that B-FRET enables sufficiently precise estimation of FRET indices in *any* condition: instead, B-FRET gives the statistically optimal inference under a certain set of assumptions. It is possible that the ‘optimal’ result does not meet the demand of an experimenter, if the experiment is not constrained enough. It is beyond the scope of the current study to explore non-standard experimental conditions because they are highly variable among experiments. In future studies, it will be important to investigate what experimental conditions better confine a FRET index, and how it depends on different FRET indices.

As a result of B-FRET’s ability to easily incorporate different sets of assumptions, one can analyze both bimolecular and unimolecular FRET using the same framework; only a slightly different set of assumptions needs to be made for unimolecular FRET due to the fixed stoichiometry of the donor and acceptor. In Supplementary Note 1, we derive the photophysical model for unimolecular FRET systems. Based on this model, we analyze unimolecular FRET data obtained from eukaryotic cells and demonstrate that it improves the SNR of an estimated FRET index (Supplementary Note 5 and Supplementary Fig. 3).

As with all other quantitative FRET methods, the applications of B-FRET are naturally limited by our understanding of the photophysical processes involved in a FRET measurement. For example, photoconversion of fluorescent proteins, which were reported to occur upon excitation for some fluorescence proteins in certain conditions^2,36,37^, can produce another chemical species that are dissimilar to both donor and acceptor during a measurement. If such secondary processes are significant but not taken into consideration in the photophysical model (Online Methods), B-FRET can yield misleading results. Although it is possible to incorporate such processes into our photophysical model once they are characterized, B-FRET does not alleviate the necessity for careful selection of a FRET pair and for control experiments to validate the basic assumptions involved in the data analysis. A distinct advantage of B-FRET is that its assumptions are explicit – helping experimenters identify necessary controls and tailor their experiments accordingly.

Finally, we comment on time-lapse FRET imaging with sub-compartment resolution. Previously, to quantify the dynamics of the degree of molecular interaction with sub-compartment resolution, methods like E-FRET were applied on a pixel-by-pixel basis and FRET indices were computed for individual pixels^12^. Nothing prevents applying B-FRET to data obtained from individual pixels, in principle. However, quantitative interpretation of FRET indices computed in such a way is not as straightforward, because assumptions that can be readily validated at the compartment level may not be validated at the sub-compartment level. For example, the total concentrations of fluorescent proteins are often assumed constant at the compartment level for E-FRET (and in the examples in this paper). This is valid, at least to a first approximation, at the compartment level as far as the measurement duration is sufficiently shorter than the time scale of factors that can change the protein concentrations, e.g., gene expression. However, this does not necessarily mean that it also holds at the sub-compartment level – the spatial distributions of proteins may dynamically change within a compartment without changing the total concentration. We are not aware of any quantitative FRET method that generally provides molecular interpretation at the sub-compartment level. Given the general demand to resolve molecular interactions at the subcellular level in cell biology, it will be an important future direction to develop such a quantitative FRET method.

## Online methods

### Strains and plasmids for the bimolecular FRET experiment

The *E. coli* strain used for the bimolecular FRET experiments is a derivative of *E. coli* K-12 strain RP437 (HCB33), and described in detail elsewhere^25,26^. In brief, the FRET acceptor-donor pair (CheY-mRFP and CheZ-mYFP) is expressed in tandem from plasmid pSJAB106^25^ under an isopropyl β-D-thiogalactopyranoside (IPTG)-inducible promoter. The glass-adhesive mutant of FliC (FliC*) was expressed from a sodium salicylate (NaSal)-inducible pZR1 plasmid. The plasmids are transformed in VS115, a cheY cheZ fliC mutant of RP437 (gift of V. Sourjik). The crosstalk coefficient for spectral bleedthrough was measured using a strain expressing CheZ-YFP from a plasmid, and that for cross-excitation was measured using a strain expressing CheY-mRFP from a plasmid (Supplementary Note 3).

### Cell preparation and bimolecular FRET measurement in a microfluidic device

Single-cell FRET microscopy and cell culture was carried out essentially as described previously^25–27^. In brief, cells were picked from a frozen stock at −80°C and inoculated in 2 mL of Tryptone Broth (TB; 1% bacto tryptone, 0.5 % NaCl) and grown overnight to saturation at 30°C and shaken at 250 RPM. Cells from a saturated overnight culture were diluted 100X in 10 mL TB and grown to OD600 0.45-0.47 in the presence of 100 μg/ml ampicillin, 34 μg/ml chloramphenicol, 50 μM IPTG and 3 μM NaSal, at 33.5°C and 250 RPM shaking. Cells were collected by centrifugation (5 min at 5000 rpm, or 4080 RCF) and washed twice with motility buffer (10 mM KPO4, 0.1 mM EDTA, 1 μM methionine, 10 mM lactic acid, pH 7), and then were resuspended in 2 mL motility buffer. Cells were left for 2 hours before starting a measurement to let all fluorescent proteins mature. Cells in motility buffer do not synthesize new proteins due to auxotrophic limitation. All experiments were performed at 22-23°C. Microfluidic devices for the FRET experiments were constructed from polydimethylsiloxane (PDMS) and used to control stimulus levels delivered to cells following exactly the same protocol as before^26,27^.

### Single-cell bimolecular FRET imaging system

FRET imaging in the microfluidic device was performed using an inverted microscope (Eclipse Ti-E; Nikon) equipped with an oil-immersion objective lens (CFI Apo TIRF 60X Oil; Nikon). YFP was illuminated by an LED illumination system (SOLA SE, Lumencor) through an excitation bandpass filter (FF01-500/24-25; Semrock) and a dichroic mirror (F01-542/27-25F; Semrock). The fluorescence emission was led into an emission image splitter (OptoSplit II; Cairn) and further split into donor and acceptor channels by a second dichroic mirror (FF580-FDi01-25×36; Semrock). The emission was then collected through emission bandpass filters (FF520-Di02-25×36 and FF593-Di03-25×36; Semrock) by a sCMOS camera (ORCA-Flash4.0 V2; Hamamatsu). RFP was illuminated in the same way as YFP except that an excitation bandpass filter (FF01-575/05-25; Semrock) and a dichroic mirror (FF593-Di03-25×36; Semorock) were used. An additional excitation filter (59026x; Chroma) was used in front of the excitation filters. To synchronize image acquisition and the delivery of stimulus solutions, a custom-made MATLAB program controlled both the imaging system (through the API provided by Micro-Manager^38^) and the states of the solenoid valves.

### Photophysical model

Here, we consider the case of bi-molecular FRET systems discussed in the main text; however, essentially the same argument applies to uni-molecular FRET systems (Supplementary Note 1).

First, we define the FRET data set. Time-lapse measurements of *I_DD_, I_DA_*, and *I_AA_* are conducted at discrete time points. We assume that FRET from the donor to acceptor affects *I_DD_* and *I_DA_*, but not *I_AA_* (See below and Supplementary Note 1). Thus, the sampling frequency of *I_DD_* and/or *I_DA_* limits the temporal resolution of an estimated FRET signal. In practice, *I_DD_* and *I_DA_* are measured (almost) simultaneously to better exploit the FRET-induced changes in *I_DD_* and *I_DA_*. Thus, we designate the same time points for the *I_DD_* and *I_DA_* measurements, and the set of the time points are written as 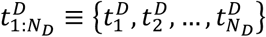, where *N_D_* is the total number of measurements. *I_AA_* is generally acquired at different time points from *I_DD_* and *I_DA_*, and thus we designate the time points for *I_AA_* as 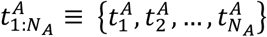, where *N_A_* is the total number of measurements, and generally *N_D_* ≠ *N_A_*. The entire set of the time-lapse fluorescence intensity data is 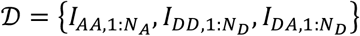, where

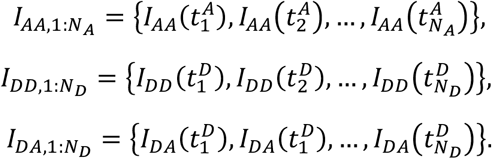

Next, we construct a photophysical model 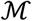 to be learned from the data 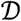. Under a standard 3-cube FRET-microscopy setup, the (background-subtracted) observables *I_AA_, I_DD_* and *I_DA_* are generally linked to the concentrations of the chemical species as follows:

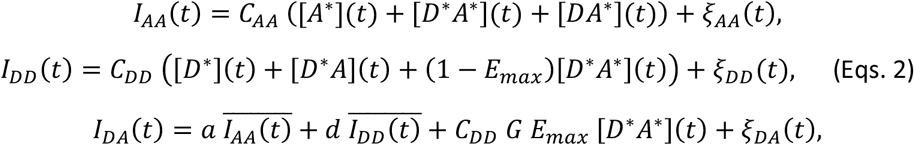

where *ξ_AA_, ξ_DD_* and *ξ_DA_* describe the measurement noise of corresponding fluorescent channels, and we assume they follow the zero-mean Gaussian distributions, i.e., 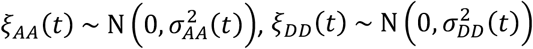, and 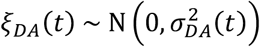, where 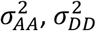, and 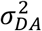 are time-dependent variances and are determined from the data (Supplementary Note 3; Note parametric noise models other than Gaussian distributions can also be used if necessary); 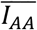 and 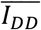 are respectively the expectation values of *I_AA_* and *I_DD_*, and thus 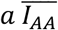 and 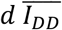 respectively represent the cross-excitation of the acceptor by the donor excitation wavelengths and the bleedthrough of the donor emission into the acceptor emission filter^4,10,11^; *C_AA_, C_DD_, a, d*, and *G* are parameters dependent on imaging systems and the photophysical properties of the donor and acceptor, which are defined as

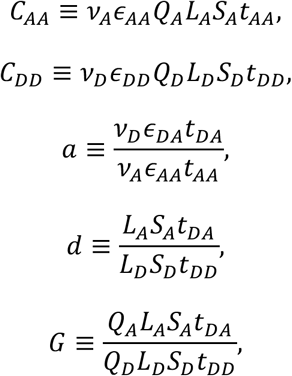

where, *v_D_* (*v_A_*) is the intensity of illumination reaching the sample through the donor (acceptor) excitation filter, *ϵ_DD_* the absorption coefficient of the donor, *ϵ_DA_* (*ϵ_AA_*) the absorption coefficient of the acceptor at the donor-excitation (acceptor-excitation) wavelength, *Q_D_* (*Q_A_*) the quantum yield of donor (acceptor), *L_D_* (*L_A_*) the throughput of the donor (acceptor) emission light-path, *S_D_* (*S_A_*) the quantum sensitivity of the camera for donor (acceptor) emission, and *t_DA_, t_AA_*, and *t_DD_* respectively the exposure time for the FRET, acceptor, and donor channels^12,13,24^. The parameters *a, d* and *G* can be determined by independent measurements^12,27^. *C_DD_* and *C_AA_* do not necessarily need to be determined as explained below. The model (Eqs. 2) is general, only assuming that the acceptor fluorescence is not detectable through the donor emission filter and that the acceptor excitation light does not excite the donor, which are easily achieved by selecting appropriate filter sets^12,27^.

We introduce the following set of assumptions, which are satisfied in a typical FRET experiment and used also in E-FRET^12^ (see also Supplementary Note 1). (i) The total amount of donor and acceptor molecules are conserved during the course of a measurement. (ii) The photobleaching locally follows a first-order decay process, i.e., the rate of change of the amount of intact (i.e., fluorescent) donor (acceptor) is proportional to its concentration, although the proportionality constants can change over time. (iii) The system is in a quasi-steady state at each time point with the timescale of photobleaching is much larger than other relevant timescales (e.g., that of binding and unbinding of the fluorescently-labeled proteins). See Supplementary Note 1 for how these are expressed mathematically.

Without loss of generality we set *C_AA_* = *C_DD_* = 1, because these parameters only affect the units of the concentrations of the chemical species (see Discussion). Also, we introduce a new label *χ*(*t*) = *E_max_* [*DA_total_*](*t*) because we are not necessarily interested in decomposing *E_max_* and [*DA_total_*](*t*), which only appear as a product in our definition of the FRET index *E*. Under these assumptions, Equations 2 is reduced to

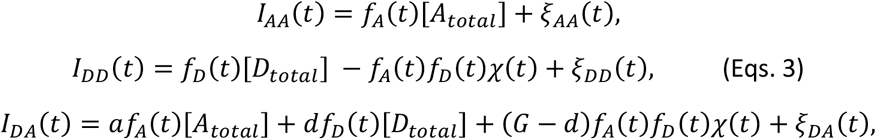

where [*A*_total_] and [*D_total_*] are the total concentrations of the acceptor and donor respectively; *f_A_* (*t*) and *f_D_* (*t*) are the intact fractions of the acceptor and donor at time *t* respectively, and hence take values between 0 and 1 (see Supplementary Note 1 for derivation). To learn the model from data, *f_A_* (*t*) and *f_D_* (*t*) need to be expressed by parametric functions, whose parameters, as well as other parameters, are estimated by the inference algorithm described below. Any parametric functions can be used depending on the data in principle. See Supplementary Note 5 for the specific functions used to analyze data used in this paper.

The presence of the hidden variable *χ*(*t*) in the equations for *I_DD_* and *I_DA_* (Eqs. 3) makes the learning of the model less straightforward. To deal with this, we rewrite the equations for *I_DD_* and *I_DA_* using the framework of the state-space model^20,21^:

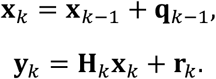

The first line, the dynamic model, describes the time evolution of the state 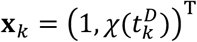. The process noise **q**_*k*-1_ governs the transition between two consecutive states. For example, Gaussian process noise can be written as:

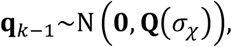

where the covariance matrix **Q**(*σ_χ_*) is defined as

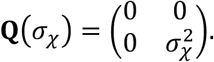

For non-Gaussian dynamics, one can use the Student’s t-distribution^19,21^, which can be written as

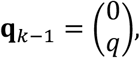

and

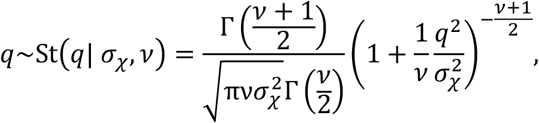

where *σ_x_* is the scale parameter and *v* > 0 is called the degree of freedom. When *v* = 1, the t-distribution reduces to the Cauchy or Lorentz distribution, while for *v* ≫ 5 it approaches a Gaussian distribution 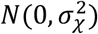.

The second line, the measurement model, describes the relationship between the observables 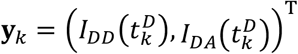 given the state **x**_*k*_. The measurement model matrix **H**_*k*_ at time 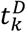 is defined as

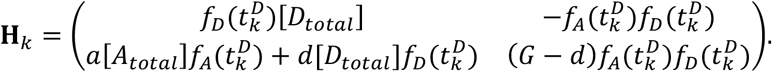

The Gaussian measurement noise **r**_*k*_ at time 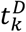 is written as

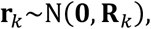

where the covariance matrix **R**_*k*_ is defined as

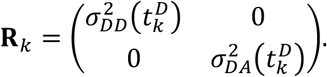

The variances of measurement noise can be determined from data (Supplementary Note 3).

### Learning algorithm

To compute the posterior distribution of the FRET index *E_k_*, 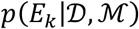, where 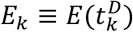, we rewrite it in term of the model parameters **θ**:

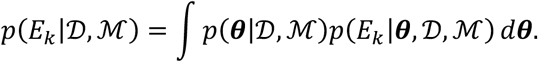

The right-hand side is the expectation of a function of model parameters **θ**, 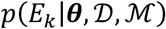, with respect to the (posterior) distribution of ***θ***, 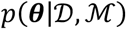. Thus, we evaluate the integral on the righthand side by a Monte Carlo approach, drawing many samples from the two probability distributions as described below.

First, we evaluate 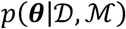. Using Bayes’ rule, this can be written as

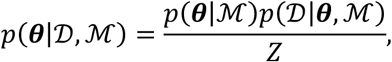

where 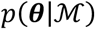 is the prior distribution of the model parameters, which are usually wide in width to express one’s ignorance about the parameter values (See Supplementary Note 2 for how to design prior distributions for each model parameter and Supplementary Note 5 for the actual distributions used to analyze data presented in this paper); 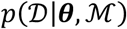 is the likelihood function, which describes the probability of observed data as a function of model parameters ***θ***; and *Z* is the normalization constant, which one does not have to evaluate for the purpose of drawing samples form 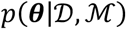. The prior distribution 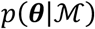 is given by the user of B-FRET and the likelihood function 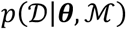 is evaluated by using the Bayesian filtering algorithm^20,21^ (Supplementary Note 2). Then, using a sampling method^18,19^, one can draw a set of samples 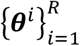 from the distribution, where 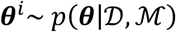 and *R* (≫ 1) is the number of samples. Samples were drawn either directly from the distribution using a Markov chain Monte Carlo (MCMC) method (e.g., slice sampling^18,19^), or from an approximated Gaussian distribution obtained by Laplace’s method^18,19^. In drawing many samples, the latter is computationally much cheaper, and thus we adopted it upon confirmation that the bias introduced by the approximation is negligible (Supplementary Fig. 4).

Second, we evaluate the (posterior) distribution of the FRET index 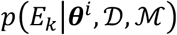 based on Bayesian smoothing algorithm^20,21^, using the sampled parameter set 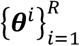 (Supplementary Note 2). This enables to draw samples 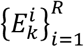 from the distribution, where 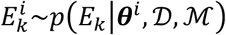. Using the samples, we can evaluate the integral as

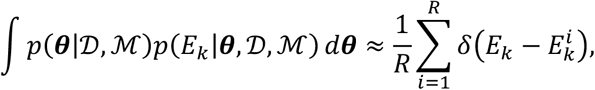

where *δ*(*x*) is the Dirac delta function. With sufficiently large *R* samples from 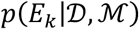, one can quantify any properties of the distribution 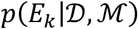. We used the median of the sample as representative values of the estimates, and the interval between 2.27 and 97.73 percentiles, each of which corresponds to *μ* ± 2*σ* respectively for a Gaussian distribution N(*μ, σ*), as a measure of the statistical uncertainty of the estimation and called it a ‘95% credible interval (CI)’.

### E-FRET method and the effect of error in optical parameter estimation

The E-FRET method^12^ provides a formula for a FRET index *E_corr_* that gives an estimate of Eq. 1. This reads

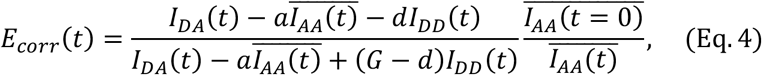

Where the optical parameters *a, d*, and *G* are defined in **Photophysical model**. For the variables with bars, e.g., 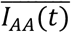, their expected (or smoothed) values can be used as opposed to raw intensity values. It can be shown that under the assumptions described in **Photophysical model** and in the limit of zero measurement noise, this quantity converges to the FRET index defined by Eq. 1 (Supplementary Note 1). The optical parameters *a, d*, and *G* are measured from independent measurements, but only with finite precision. The errors in the estimations of these parameters introduce some biases in the computed FRET index, whose effect grows as more fluorescent proteins are photobleached, which can be corrected under some assumptions^27^ (See Supplementary Note 3 for more detail).

### Model selection

In case a user of B-FRET is not sure about what model to use (e.g., Gaussian or non-Gaussian process noise), the framework of model selection enables to select, among a set of candidate models, a model that is best evidenced by a set of data. For this purpose, B-FRET computes the Bayesian information criterion (BIC) defined as

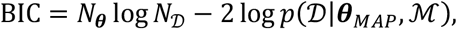

where *N_θ_* and 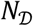 are the numbers of the model parameters and data points, respectively and ***θ**_MAP_* is the parameter values that maximize the likelihood function 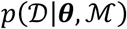. A model with the lowest BIC value is selected as the best model among a set of candidates (Supplementary Fig. 2).

## Acknowledgements

We thank H. H. Mattingly and Y. Kojima for commenting on the text. We acknowledge R. Gomez-Sjoberg, Microfluidics Lab, for providing information and software to control the solenoid valves in the microfluidic setup. This works was supported by NIH awards R01GM106189 and R01GM138533 (KK and TE); JST, PRESTO Grant Number JPMJPR21E4, Japan (KK); a postdoctoral fellowship through the Swartz Foundation for Theoretical Neuroscience and postdoctoral fellowships NIH F32MH118700 and NIH K99DC019397 (NK); JSPS KAKENHI Grants, no. 19H05798 (AK);

## Author information

### Contributions

KK designed the research. KK developed the algorithm and implemented it in MATLAB with inputs from NK. NK implemented the algorithm in Python. KK performed the experiments in *E. coli.* KK, TE and TSS analyzed the results. KA performed the experiments with HeLa cells and KA and KK analyzed the results. KK, NK, and TE wrote the manuscript. All authors edited and approved the manuscript.

## Data availability

Data will be made available upon publication.

## Code availability

Codes written in MATLAB and Python will be made available on the Emonet lab git website (github.com/emonetlab) upon publication.

## References

1. Tsien, R. Y. Building and breeding molecules to spy on cells and tumors. FEBS Lett. 579, 927–932 (2005).

2. Miyawaki, A. Development of Probes for Cellular Functions Using Fluorescent Proteins and Fluorescence Resonance Energy Transfer. Annu. Rev. Biochem. 80, 357–373 (2011).

3. Miyawaki, A. & Niino, Y. Molecular Spies for Bioimaging—Fluorescent Protein-Based Probes. Mol. Cell 58, 632–643 (2015).

4. Miyawaki, A. & Tsien, R. Y. Monitoring protein conformations and interactions by fluorescence resonance energy transfer between mutants of green fluorescent protein. in Methods in Enzymology vol. 327 472–500 (Elsevier, 2000).

5. Bajar, B., Wang, E., Zhang, S., Lin, M. & Chu, J. A Guide to Fluorescent Protein FRET Pairs. Sensors 16, 1488 (2016).

6. Zhang, J., Campbell, R. E., Ting, A. Y. & Tsien, R. Y. Creating new fluorescent probes for cell biology. Nat. Rev. Mol. Cell Biol. 3, 906–918 (2002).

7. Lam, A. J. et al. Improving FRET dynamic range with bright green and red fluorescent proteins. Nat. Methods 9, 1005–1012 (2012).

8. Lichtman, J. W. & Conchello, J.-A. Fluorescence microscopy. Nat. Methods 2, 910–919 (2005).

9. Sanderson, M. J., Smith, I., Parker, I. & Bootman, M. D. Fluorescence Microscopy. Cold Spring Harb. Protoc. 2014, pdb.top071795 (2014).

10. Youvan, D. C. et al. Calibration of Fluorescence Resonance Energy Transfer in Microscopy Using Genetically Engineered GFP Derivatives on Nickel Chelating Beads. 18.

11. Gordon, G. W., Berry, G., Liang, X. H., Levine, B. & Herman, B. Quantitative Fluorescence Resonance Energy Transfer Measurements Using Fluorescence Microscopy. Biophys. J. 74, 2702–2713 (1998).

12. Zal, T. & Gascoigne, N. R. J. Photobleaching-Corrected FRET Efficiency Imaging of Live Cells. Biophys. J. 86, 3923–3939 (2004).

13. Lichten, C. A. & Swain, P. S. A Bayesian method for inferring quantitative information from FRET data. BMC Biophys. 4, 10 (2011).

14. Zeug, A., Woehler, A., Neher, E. & Ponimaskin, E. G. Quantitative Intensity-Based FRET Approaches—A Comparative Snapshot. Biophys. J. 103, 1821–1827 (2012).

15. Berney, C. & Danuser, G. FRET or No FRET: A Quantitative Comparison. Biophys. J. 84, 3992–4010 (2003).

16. Vogel, S. S., Thaler, C. & Koushik, S. V. Fanciful FRET. Sci. STKE Signal Transduct. Knowl. Environ. 2006, re2 (2006).

17. Miyawaki, A. et al. Fluorescent indicators for Ca2+based on green fluorescent proteins and calmodulin. Nature 388, 882–887 (1997).

18. MacKay, D. J. C. Information Theory, Inference, and Learning Algorithms. 640.

19. Bishop, C. M. Pattern recognition and machine learning. (Springer, 2006).

20. Sarkka, S. Bayesian Filtering and Smoothing. (Cambridge University Press, 2013). doi:10.1017/CBO9781139344203.

21. Kitagawa, G. Introduction to time series modeling. (Chapman and Hall/CRC, 2010).

22. Sourjik, V. & Berg, H. C. Receptor sensitivity in bacterial chemotaxis. Proc. Natl. Acad. Sci. 99, 123–127 (2002).

23. Sourjik, V., Vaknin, A., Shimizu, T. S. & Berg, H. C. In Vivo Measurement by FRET of Pathway Activity in Bacterial Chemotaxis. in Methods in Enzymology vol. 423 365–391 (Elsevier, 2007).

24. Wlodarczyk, J. et al. Analysis of FRET Signals in the Presence of Free Donors and Acceptors. Biophys. J. 94, 986–1000 (2008).

25. Keegstra, J. M. et al. Phenotypic diversity and temporal variability in a bacterial signaling network revealed by single-cell FRET. eLife 6, e27455 (2017).

26. Kamino, K., Keegstra, J. M., Long, J., Emonet, T. & Shimizu, T. S. Adaptive tuning of cell sensory diversity without changes in gene expression. Sci. Adv. 6, eabc1087 (2020).

27. Mattingly, H. H., Kamino, K., Machta, B. B. & Emonet, T. Escherichia coli chemotaxis is information limited. Nat. Phys. 17, 1426–1431 (2021).

28. Parkinson, J. S., Hazelbauer, G. L. & Falke, J. J. Signaling and sensory adaptation in Escherichia coli chemoreceptors: 2015 update. Trends Microbiol. 23, 257–266 (2015).

29. Alon, U., Surette, M. G., Barkai, N. & Leibler, S. Robustness in bacterial chemotaxis. 397, 4 (1999).

30. Kamino, K., Fujimoto, K. & Sawai, S. Collective oscillations in developing cells: Insights from simple systems: Collective oscillations in developing cells. Dev. Growth Differ. 53, 503–517 (2011).

31. Kamino, K. & Kondo, Y. Rescaling of Spatio-Temporal Sensing in Eukaryotic Chemotaxis. PLOS ONE 18 (2016).

32. Kamino, K. et al. Fold-change detection and scale invariance of cell–cell signaling in social amoeba. Proc. Natl. Acad. Sci. 114, E4149–E4157 (2017).

33. Moore, J. P., Kamino, K. & Emonet, T. Non-Genetic Diversity in Chemosensing and Chemotactic Behavior. Int. J. Mol. Sci. 22, 6960 (2021).

34. Karin, O. & Alon, U. Temporal fluctuations in chemotaxis gain implement a simulated-tempering strategy for efficient navigation in complex environments. iScience 24, 102796 (2021).

35. Frankel, N. W. et al. Adaptability of non-genetic diversity in bacterial chemotaxis. eLife 3, e03526 (2014).

36. Malkani, N. & Schmid, J. A. Some Secrets of Fluorescent Proteins: Distinct Bleaching in Various Mounting Fluids and Photoactivation of Cyan Fluorescent Proteins at YFP-Excitation. PLoS ONE 6, e18586 (2011).

37. Kirber, M. T., Chen, K. & Keaney, J. F. YFP photoconversion revisited: confirmation of the CFP-like species. Nat. Methods 4, 767–768 (2007).

38. Edelstein, A., Amodaj, N., Hoover, K., Vale, R. & Stuurman, N. Computer Control of Microscopes Using μManager. Curr. Protoc. Mol. Biol. 92, (2010).

