## Supplementary Information for "Optimal inference of molecular interactions in live FRET imaging"

### Supplementary Note 1

#### Deriving photophysical models of FRET measurements

Keita Kamino, Nirag Kadakia, Kazuhiro Aoki, Thomas S. Shimizu, and Thierry Emonet

Here we derive the photophysical models of FRET measurements (i.e., the measurement model in the state-space model). Both bimolecular and unimolecular FRET systems are considered. We also derive the E-FRET formula so that the underlying assumptions are clearly seen.

**General measurement model for bimolecular FRET systems**

A general photophysical model that links the concentrations of chemical species and (background-subtracted) fluorescence intensities is written as:

$$I_{AA}(t) = C_{AA}\left([A^*](t) + [D^*A^*](t) + [DA^*](t)\right) + \xi_{AA}(t),$$

$$I_{DD}(t) = C_{DD}\left([D^*](t) + [D^*A](t) + (1 - E_{max})[D^*A^*](t)\right) + \xi_{DD}(t), \quad \text{(Eqs. 1-1)}$$

$$I_{DA}(t) = a\,\overline{I_{AA}(t)} + d\,\overline{I_{DD}(t)} + C_{DD}\,G\,E_{max}\,[D^*A^*](t) + \xi_{DA}(t).$$

See Online Methods for definitions of each term and parameter. We reiterate that $\xi_{AA}(t)$, $\xi_{DD}(t)$ and $\xi_{DA}(t)$ are zero-mean stochastic variables – typically Gaussian distributions – describing measurement noise of respective fluorescence signals, and we assume that their magnitudes are estimated independently from the B-FRET algorithm (Supplementary Note 3). Also, the parameters that depend on the imaging system, $a, d$ and $G$ are determined from independent measurements (Supplementary Note 3). This general model (Eqs. 1-1) contains five latent variables (i.e., $[A^*](t)$, $[D^*](t)$, $[DA^*](t)$, $[D^*A](t)$ and $[D^*A^*](t)$), and thus the problem of inferring these variables by observing three variables (i.e., $I_{AA}(t)$, $I_{DD}(t)$, and $I_{DA}(t)$) is inherently ill-posed. However, in typical FRET experiments, experimental design can be used to constrain the degrees of freedom of the system and obtain interpretable results. Below, we consider the typical set of constraints satisfied by most FRET experiments – the same set of assumptions used by E-FRET – and how they constrain the general model (Eqs. 1-1).

**A reduced measurement model for bimolecular FRET systems**

We first assume that the total concentrations of the donor and acceptor are conserved during a measurement:

$$[A_{total}] = [A^*](t) + [A](t) + [D^*A^*](t) + [D^*A](t) + [DA^*](t) + [DA](t),$$

$$[D_{total}] = [D^*](t) + [D](t) + [D^*A^*](t) + [D^*A](t) + [DA^*](t) + [DA](t),$$

where $[A_{\mathrm{total}}]$ and $[D_{total}]$ are the total concentrations of the acceptor and donor respectively.

Next, we assume that the photobleaching rate is a first-order decay process. Combined with the assumption that there's no synthesis of new fluorescent proteins during a measurement, this leads to:

$$\frac{d([A^*](t) + [D^*A^*](t) + [DA^*](t))}{dt} = -\alpha(t)([A^*](t) + [D^*A^*](t) + [DA^*](t)),$$

$$\frac{d([D^*](t) + [D^*A^*](t) + [D^*A](t))}{dt} = -\delta(t)([D^*](t) + [D^*A^*](t) + [D^*A](t)),$$

where $\alpha(t)$ and $\delta(t)$ are the bleaching rates of the acceptor and donor at time $t$. Solving these, we get:

$$[A^*](t) + [D^*A^*](t) + [DA^*](t) = [A_{total}]\, e^{-\int_0^t \alpha(t')dt'} = [A_{total}] f_A(t, \boldsymbol{\theta}_A),$$

$$[D^*](t) + [D^*A^*](t) + [D^*A](t) = [D_{total}]\, e^{-\int_0^t \delta(t')dt'} = [D_{total}] f_D(t, \boldsymbol{\theta}_D),$$

where we introduced the functions $f_A(t, \boldsymbol{\theta}_A) = e^{-\int_0^t \alpha(t')dt'}$ and $f_D(t, \boldsymbol{\theta}_D) = e^{-\int_0^t \delta(t')dt'}$ parameterized by $\boldsymbol{\theta}_A$ and $\boldsymbol{\theta}_D$ respectively. $f_A(t, \boldsymbol{\theta}_A)$ and $f_D(t, \boldsymbol{\theta}_D)$ represent the intact fractions of the acceptor and donor at time $t$, respectively.

Lastly, we assume that the system is in a quasi-steady state at each time point. Namely, the timescale of photobleaching is sufficiently longer than the following two timescales: (i) the timescale of the binding and unbinding of the molecules $\mathrm{X}$ and $\mathrm{Y}$, to which the donor and acceptor are fused respectively, and (ii) the timescale of diffusion of either the donor or acceptor over the compartment that enclose the molecules. With this assumption, the fraction of *each free or complexed species* that is intact decays exponentially. In particular, we get:

$$\frac{[A^*](t)}{[A^*](t) + [A](t)} = f_A(t, \boldsymbol{\theta}_A),$$

$$\frac{[D^*](t)}{[D^*](t) + [D](t)} = f_D(t, \boldsymbol{\theta}_D).$$

Furthermore, defining $\gamma(t)$ as the binding affinity constant between $\mathrm{X}$ and $\mathrm{Y}$ at time $t$:

$$[D^*A^*](t) = \gamma(t)\ [D^*](t)[A^*](t)$$

$$= \gamma(t) \Big(\big([D^*](t) + [D](t)\big) f_D(t, \boldsymbol{\theta}_D)\Big) \Big(\big([A^*](t) + [A](t)\big) f_A(t, \boldsymbol{\theta}_A)\Big)$$

$$= \gamma(t)([X](t) f_D(t, \boldsymbol{\theta}_D))\big([Y](t) f_A(t, \boldsymbol{\theta}_A)\big)$$

$$= [DA_{total}]\ (t) f_A(t, \boldsymbol{\theta}_A) f_D(t, \boldsymbol{\theta}_D),$$

where $[X](t) = [D^*](t) + [D](t)$ and $[Y](t) = [A^*](t) + [A](t)$, and $[DA_{total}]\ (t) = [D^*A^*](t) + [D^*A](t) + [DA^*](t) + [DA](t)$.

Under these assumptions, the general model (Eqs. 1-1) becomes

$$I_{AA}(t) = C_{AA}\, f_A(t, \boldsymbol{\theta}_A)[A_{total}] + \xi_{AA}(t),$$

$$I_{DD}(t) = C_{DD}(f_D(t, \boldsymbol{\theta}_D)[D_{total}] - f_A(t, \boldsymbol{\theta}_A) f_D(t, \boldsymbol{\theta}_D) E_{max}\,[DA_{total}](t)) + \xi_{DD}(t), \qquad \text{(Eqs. 1-2)}$$

$$I_{\mathrm{DA}}(t) = a\,\overline{I_{AA}(t)} + d\,\overline{I_{DD}(t)} + C_{DD} f_A(t, \boldsymbol{\theta}_A) f_D(t, \boldsymbol{\theta}_D) G\, E_{max}\,[DA_{total}](t) + \xi_{DA}(t).$$

Without loss of generality, we set $C_{AA} = C_{DD} = 1$. This is because these parameters only determine the units of the concentrations of the chemical species. Although setting $C_{AA} = C_{DD} = 1$ means that we use different concentration units for $[A_{total}]$ and $[D_{total}]$ and $[DA_{total}]$, this doesn't affect the estimation of the FRET index $E = \frac{E_{max}[DA_{total}]}{[D_{total}]}$ because both the numerator and denominator of $E$ are measured by the same concentration unit. Also, since we are not necessarily interested in estimating $E_{max}$ and $[DA_{total}]$ separately, we assign a single time-dependent variable to the product, i.e., $E_{max}\,[DA_{total}](t) = \chi(t)$ . Note, however, one can easily incorporate measurements of the parameters $C_{AA}, C_{DD}$ and $E_{max}$, and infer different FRET indices (e.g., $E_1 = \frac{[DA_{total}]}{[D_{total}]}$ or $E_2 = \frac{[DA_{total}]}{[A_{total}]}$) within the B-FRET framework.

Finally, we discretize the model in time since actual measurements are conducted at discrete time points. For an effective FRET measurement, $I_{DD}$ and $I_{DA}$ have to be measured simultaneously or consecutively with sufficiently small time interval compared to the timescale of the changes in the donor-acceptor interaction, $\gamma(t)$; so, we designate the time points for $I_{DD}$ and $I_{DA}$ as $t^D_{1:N_D} \equiv \{t^D_1, t^D_2, \dots, t^D_{N_D}\}$. $I_{AA}$ can be measured independently from – and often less frequently than – the measurements of $I_{DD}$ and $I_{DA}$, because $I_{AA}$ does not depend on the FRET between the donor and acceptor. Accordingly, we define the $I_{AA}$ measurement time points as $t^A_{1:N_A} \equiv \{t^A_1, t^A_2, \dots, t^A_{N_A}\}$. Generally $N_D \neq N_A$, and the measurement intervals do not have to be constant.

Using these assumptions and the notation above, we get the probabilistic measurement model of the observables for the bimolecular FRET system:

$$I_{AA}(t^A_j) = f_A(t^A_j, \boldsymbol{\theta}_A)[A_{total}] + \xi_{AA}(t^A_j),$$

$$I_{DD}(t^D_k) = f_D(t^D_k, \boldsymbol{\theta}_D)[D_{total}] - f_A(t^A_k, \boldsymbol{\theta}_A) f_D(t^D_k, \boldsymbol{\theta}_D)\chi_k + \xi_{DD}(t^D_k), \qquad \text{(Eqs. 1-3)}$$

$$I_{\mathrm{DA}}(t^D_k) = a f_A(t^A_k, \boldsymbol{\theta}_A)[A_{total}] + d f_D(t^D_k, \boldsymbol{\theta}_D)[D_{total}] + (G-d) f_A(t^A_k, \boldsymbol{\theta}_A) f_D(t^D_k, \boldsymbol{\theta}_D)\chi_k + \xi_{DA}(t^D_k),$$

where $\chi_k \equiv \chi(t^D_k)$.

**General measurement model for unimolecular FRET systems**

A unimolecular FRET-sensor molecule consists of a donor and acceptor domains and a sensor domain. The donor and acceptor domains flank the sensor domain. The sensor domain changes its conformation upon binding to a cognate molecule, which causes the change in the distance between the donor and acceptor and thus the level of FRET. Unimolecular FRET systems are different from bimolecular FRET systems in that: (i) the donor-acceptor stoichiometry is fixed to 1:1 in a unimolecular system, while it is variable in a bimolecular system, and (ii) in an unimolecular system, there can be finite basal FRET between the donor and acceptor even when the sensor is in a 'off' state, while in a bimolecular FRET system there is essentially no FRET between free donor and acceptor unless the concentrations of the fluorescent proteins are very high (in which case we can easily incorporate the effect in the measurement model). We consider the FRET sensor in a unimolecular FRET system as a two-state molecule that can be either in active or inactive state when the donor and acceptor are close or distant to each other. When both donor and acceptor are intact, we assume that an active sensor shows high specific FRET efficiency $E_{max}$, and an inactive sensor low specific FRET efficiency $E_{min}$. The system consists of the following eight chemical species: $\mathrm{D^*A^*}, \mathrm{D^*A}, \mathrm{DA^*}, D^*\cdot A^*, D^*\cdot A, D\cdot A^*$, and $D\cdot A$, where a chemical species with a dot '·' is in an inactive state, and without a dot an active state. Similarly to the bimolecular-FRET model, the observables $I_{AA}, I_{DD}$ and $I_{DA}$ are linked to the concentrations of the chemical species as follows:

$$I_{AA}(t) = C_{AA}\,([D^*A^*](t) + [DA^*](t) + [D\cdot A^*](t) + +[D^*\cdot A^*](t)) + \xi_{AA}(t),$$

$$I_{DD}(t) = C_{DD}\left([D^*A](t) + [D^*\cdot A](t) + (1 - E_{max})[D^*A^*](t) + (1 - E_{min})[D^*\cdot A^*](t)\right) + \xi_{DD}(t), \text{ (Eqs. 1-4)}$$

$$I_{DA}(t) = a\,\overline{I_{AA}(t)} + d\,\overline{I_{DD}(t)} + C_{DD}\,G\left(E_{max}\,[D^*A^*](t) + E_{min}\,[D^*\cdot A^*](t)\right) + \xi_{DA}(t).$$

The parameters are defined in the same way as the bimolecular FRET model. Again, $\xi_{AA}(t)$, $\xi_{DD}(t)$ and $\xi_{DA}(t)$ are zero-mean stochastic variables describing measurement noise of respective fluorescence signals, and we assume that their magnitudes are estimated independently from the B-FRET algorithm.

**A reduced measurement model for unimolecular FRET systems**

We first assume that the total concentration of the FRET sensor molecule does not change during a measurement:

$$C_{total} = [D^*A^*](t) + [D^*A](t) + [DA^*](t) + [DA](t) + [D^*\cdot A^*](t) + [D^*\cdot A](t) + [D\cdot A^*](t) + [D\cdot A](t).$$

We also introduce variables to describe the total concentrations of active- and inactive-state sensors respectively:

$$[DA_{total}](t) = [D^*A^*](t) + [D^*A](t) + [DA^*](t) + [DA](t),$$

$$[D\cdot A_{total}](t) = [D^*\cdot A^*](t) + [D^*\cdot A](t) + [D\cdot A^*](t) + [D\cdot A](t).$$

Next, we assume that the photobleaching rate is a first-order decay process:

$$\frac{d([D^*A^*](t) + [DA^*](t) + [D^*\cdot A^*](t) + [D\cdot A^*](t))}{dt} = -\alpha(t)([D^*A^*](t) + [DA^*](t) + [D^*\cdot A^*](t) + [D\cdot A^*](t)),$$

$$\frac{d([D^*A^*](t) + [D^*A](t) + [D^*\cdot A^*](t) + [D^*\cdot A](t))}{dt} = -\delta(t)([D^*A^*](t) + [D^*A](t) + [D^*\cdot A^*](t) + [D^*\cdot A](t)),$$

where $\alpha(t)$ and $\delta(t)$ are the bleaching rates of the acceptor and donor and at time $t$. Solving these, we get:

$$[D^*A^*](t) + [DA^*](t) + [D^*\cdot A^*](t) + [D\cdot A^*](t) = C_{total}\,e^{-\int_0^t \alpha(t')dt'} = C_{total}\,f_A(t,\boldsymbol{\theta}_A),$$

$$[D^*A^*](t) + [D^*A](t) + [D^*\cdot A^*](t) + [D^*\cdot A](t) = C_{total}\,e^{-\int_0^t \delta(t')dt'} = C_{total}\,f_D(t,\boldsymbol{\theta}_D),$$

where we, in the same way as bimolecular FRET, introduced the functions $f_A(t,\boldsymbol{\theta}_A) = e^{-\int_0^t \alpha(t')dt'}$ and $f_D(t,\boldsymbol{\theta}_D) = f_D(t,\boldsymbol{\theta}_D)$ parameterized by $\boldsymbol{\theta}_A$ and $\boldsymbol{\theta}_D$, respectively. $f_A(t,\boldsymbol{\theta}_A)$ and $f_D(t,\boldsymbol{\theta}_D)$ represent the intact fractions of the donor and acceptor at time $t$ respectively.

Lastly, we assume that the system is in a quasi-steady state at each time point, i.e., the time scale of the active-inactive transition is sufficiently shorter than that of photobleaching. Therefore, at a certain time $t$, the intact fractions of the acceptor and donor molecules of all active-state (or inactive-state) sensor molecules are given by $f_A(t,\boldsymbol{\theta}_A)$ and $f_D(t,\boldsymbol{\theta}_D)$. This gives us the following relationships:

$$[D^*A^*](t) = f_A(t,\boldsymbol{\theta}_A)f_D(t,\boldsymbol{\theta}_D)[DA_{total}](t),$$

$$[D^*\cdot A^*](t) = f_A(t,\boldsymbol{\theta}_A)f_D(t,\boldsymbol{\theta}_D)[D\cdot A_{total}](t).$$

Under these assumptions, the general model (Eqs. 1-4) is reduced to:

$$I_{AA}(t) = C_{AA}\,f_A(t,\boldsymbol{\theta}_A)C_{total} + \xi_{AA}(t),$$

$$I_{DD}(t) = C_{DD}\left(f_D(t,\boldsymbol{\theta}_D)C_{total} - f_A(t,\boldsymbol{\theta}_A)f_D(t,\boldsymbol{\theta}_D)(E_{max}\,[DA_{total}](t) + E_{min}\,[D\cdot A_{total}](t))\right) + \xi_{DD}(t), \text{ (Eqs. 1-5)}$$

$$I_{\mathrm{DA}}(t) = a\,\overline{I_{AA}(t)} + d\,\overline{I_{DD}(t)} + C_{DD} f_A(t,\boldsymbol{\theta}_A) f_D(t,\boldsymbol{\theta}_D) G\,(E_{max}\,[DA_{total}](t) + E_{min}\,[D\cdot A_{total}](t)) + \xi_{DA}(t).$$

To simplify, we reparametrize the model. Setting $C_{AA} = C_{DD} = 1$ as in the bimolecular FRET model, we get:

$$[A_{total}] = C_{AA}C_{total} = C_{total},$$

$$[D_{total}] = C_{DD}C_{total} = C_{total},$$

$$\chi(t) = C_{DD}(E_{max}\,[DA_{total}](t) + E_{min}\,[D\cdot A_{total}](t))$$
$$= (E_{max}\,[DA_{total}](t) + E_{min}\,[D\cdot A_{total}](t)).$$

Note that because generally $C_{AA} \neq C_{DD}$, the concentration of $[A_{total}]$ is measured by a different unit from those of $[D_{total}]$ and $\chi(t)$; However, this is not an issue as far as we denominate the FRET index by $[D_{total}]$, in the same way as the bimolecular case.

By discretizing the model in time in the same way as the bimolecular FRET, we get

$$I_{AA}(t_j^A) = f_A(t_j^A,\boldsymbol{\theta}_A)[A_{total}] + \xi_{AA}(t_j^A),$$

$$I_{DD}(t_k^D) = f_D(t_k^D,\boldsymbol{\theta}_D)[D_{total}] - f_A(t_k^D,\boldsymbol{\theta}_A)f_D(t_k^D,\boldsymbol{\theta}_D)\chi_k + \xi_{DD}(t_k^D), \quad \text{(Eqs. 1-6)}$$

$$I_{DA}(t_k^D) = af_A(t_k^D,\boldsymbol{\theta}_A)[A_{total}] + df_D(t_k^D,\boldsymbol{\theta}_D)[D_{total}] + (G-d)f_A(t_k^D,\boldsymbol{\theta}_A)f_D(t_k^D,\boldsymbol{\theta}_D)\chi_k + \xi_{DA}(t_k^D).$$

Note that the apparent form of the model is identical to that of the bimolecular-FRET model, although the interpretation of $\chi(t_k) \equiv \chi_k$ is different: for bimolecular FRET $\chi(t)$ represents $E_{max}\,[DA_{total}](t)$ whereas for monomolecular FRET $\chi(t)$ represents $(E_{max}\,[DA_{total}](t) + E_{min}\,[D\cdot A_{total}](t))$. Once the parameters and $\chi_k$ are inferred from the data, one can quantify the FRET signal as

$$E_k = \frac{\chi_k}{[D_{total}]} = \frac{E_{max}\,[DA_{total}](t_k^D) + E_{min}\,[D\cdot A_{total}](t_k^D)}{[D_{total}]},$$

which has a clear interpretation given by the last expression.

**On the parameterized photobleaching functions in the measurement models**

In principle, we can use any parameterized photobleaching functions for $f_A(t,\boldsymbol{\theta}_A)$ and $f_D(t,\boldsymbol{\theta}_D)$, which describe the temporal evolutions of the fractions of intact acceptor and donor respectively, depending on the FRET experiment. The photobleaching dynamics of the fluorophores can be usually described by simple functions such as linear, single-exponential, or bi-exponential functions. Which function is more appropriate depends primarily on the degree of photobleaching, which depends on the length of a measurement and excitation intensity. For some data sets, the appropriate functional form can be unknown. In this case, different models can be compared using a model-selection criterion such as the Bayesian information criterion (Online Methods) to find out which model is best evidenced by the data.

**Derivation of E-FRET formula for bimolecular FRET**

Here, based on the original work, we re-derive the E-FRET formula (Online methods), so that assumptions involved can be seen more clearly. First, the E-FRET formula reads

$$E_{corr}(t) = \frac{I_{DA}(t) - a\overline{I_{AA}(t)} - dI_{DD}(t)}{I_{DA}(t) - a\overline{I_{AA}(t)} + (G-d)I_{DD}(t)} \frac{\overline{I_{AA}(t=0)}}{\overline{I_{AA}(t)}}.$$

The E-FRET methods asserts that this formula gives an estimate of the following FRET index,

$$E(t) = \frac{E_{max}[DA_{total}](t)}{[D_{total}]}.$$

For a unimolecular FRET system with non-zero minimum FRET efficiency, the FRET index is (see 'A reduced measurement model for unimolecular FRET systems'),

$$E(t) = \frac{E_{max}\ [DA_{total}](t) + E_{min}\ [D \cdot A_{total}](t)}{[D_{total}]}.$$

In general, $E_{corr}(t) \neq E(t)$; however, under certain assumptions, one can shown $E_{corr}(t) = E(t)$, which we show below. For brevity, we only consider the case of bimolecular FRET (i.e., $E = E_{max}\ [DA_{total}](t)/[D_{total}]$), but one can easily derive the same formula for the case of unimolecular FRET with non-zero minimum FRET.

In the above section entitled 'A reduced measurement model for bimolecular FRET systems', we assumed (i) the conservation of the total fluorescent protein concentrations, (ii) the photobleaching is a first-order decay process, and (iii) the system is in a quasi-steady state at each time point. Mathematically, these assumptions were expressed as

$$[A^*](t) + [D^*A^*](t) + [DA^*](t) = [A_{total}]f_A(t, \boldsymbol{\theta}_A),$$

$$[D^*](t) + [D^*A^*](t) + [D^*A](t) = [D_{total}]f_D(t, \boldsymbol{\theta}_D),$$

$$[D^*A^*](t) = [DA_{total}]\ (t)f_A(t, \boldsymbol{\theta}_A)f_D(t, \boldsymbol{\theta}_D).$$

Using these, the general equation that link chemical species to fluorescence intensities (Eqs. 1-1) becomes the reduced model (Eqs. 1-2), which reads

$$I_{AA}(t) = C_{AA}\ f_A(t, \boldsymbol{\theta}_A)[A_{total}]\ + \xi_{AA}(t),$$

$$I_{DD}(t) = C_{DD}(f_D(t, \boldsymbol{\theta}_D)[D_{total}]\ - f_A(t, \boldsymbol{\theta}_A)f_D(t, \boldsymbol{\theta}_D)E_{max}\ [DA_{total}](t)) + \xi_{DD}(t),$$

$$I_{\mathrm{DA}}(t) = a\ \overline{I_{AA}(t)} + d\ \overline{I_{DD}(t)} + C_{DD}f_A(t, \boldsymbol{\theta}_A)f_D(t, \boldsymbol{\theta}_D)G\ E_{max}\ [DA_{total}](t)\ + \xi_{DA}(t).$$

By plugging these expressions into the E-FRET formula, and assuming zero measurement noise ($\xi_{AA} = \xi_{DD} = \xi_{DA} = 0$), one gets

$$E_{corr}(t) = \frac{I_{DA}(t) - a\overline{I_{AA}(t)} - dI_{DD}(t)}{I_{DA}(t) - a\overline{I_{AA}(t)} + (G-d)I_{DD}(t)} \frac{\overline{I_{AA}(t=0)}}{\overline{I_{AA}(t)}}$$

$$= \frac{C_{DD}\ G\ E_{max}\ [DA_{total}](t)f_A(t, \boldsymbol{\theta}_A)f_D(t, \boldsymbol{\theta}_D)}{C_{DD}\ G\ E_{max}\ [DA_{total}](t)f_A(t, \boldsymbol{\theta}_A)f_D(t, \boldsymbol{\theta}_D) + C_{DD}G\left([D_{total}]f_D(t, \boldsymbol{\theta}_D) - E_{max}[DA_{total}](t)f_A(t, \boldsymbol{\theta}_A)f_D(t, \boldsymbol{\theta}_D)\right)} \frac{1}{f_A(t, \boldsymbol{\theta}_A)}$$

$$= \frac{E_{max}\ [DA_{total}](t)}{[D_{total}]} = E.$$

Thus, the E-FRET formula gives the estimation of the FRET index defined as $E = E_{max}\ [DA_{total}](t)/[D_{total}]$ when the three assumptions are satisfied.

### Supplementary Note 2

#### Learning algorithm and prior distributions

Keita Kamino, Nirag Kadakia, Kazuhiro Aoki, Thomas S. Shimizu, and Thierry Emonet

**Overview**

The goal of the B-FRET learning algorithm is to make an information-theoretically optimal inference of a user-defined FRET index $E$ at each time point, given a model $\mathcal{M}$ and a set of data $\mathcal{D}$.

A data set can be described as $\mathcal{D} = \{I_{AA,1:N_A}, I_{DD,1:N_D}, I_{DA,1:N_D}\}$, where

$$I_{AA,1:N_A} = \{I_{AA}(t_1^A), I_{AA}(t_2^A), \ldots, I_{AA}(t_{N_A}^A)\},$$

$$I_{DD,1:N_D} = \{I_{DD}(t_1^D), I_{DD}(t_2^D), \ldots, I_{DD}(t_{N_D}^D)\},$$

$$I_{DA,1:N_D} = \{I_{DA}(t_1^D), I_{DA}(t_1^D), \ldots, I_{DA}(t_{N_D}^D)\}.$$

Note that measurement time points for $I_{AA}$ are generally different from those of $I_{DD}$ and $I_{DA}$ (Online methods).

A model $\mathcal{M}$ can be written as

$$I_{AA}(t_j^A) = f_A(t_j^A, \boldsymbol{\theta}_A)[A_{total}] + \xi_{AA}(t_j^A), \quad (\text{Eq. } 2-1)$$

$$\mathbf{x}_k = \mathbf{x}_{k-1} + \mathbf{q}_{k-1}, \quad (\text{Eq. } 2-2)$$

$$\mathbf{y}_k = \mathbf{H}_k(\boldsymbol{\theta}_m)\mathbf{x}_k + \mathbf{r}_k. \quad (\text{Eq. } 2-3)$$

See Online Methods and Supplementary Note 1 for the derivation and the set of assumptions involved. The equation for $I_{AA}$ is separated from the state-space representation of the equations for $\mathbf{y}_k = \left(I_{DD}(t_k^D), I_{DA}(t_k^D)\right)^{\mathrm{T}}$ (Eqs. 2-2 and 2-3), because only $I_{DD}$ and $I_{DA}$ are dependent on the latent variable $\mathbf{x}_k = (1, \chi_k)^{\mathrm{T}}$, where $\chi_k \equiv \chi(t_k^D)$. We call Eq. 2-2 a dynamic model and Eq. 2-3 a measurement model. $\mathbf{q}_{k-1} = (0, q)^{\mathrm{T}}$ is the process noise, where the stochastic variable $q$ follows a zero-mean probability distribution parameterized by $\boldsymbol{\theta}_{\boldsymbol{q}}$, $q \sim p(q|\boldsymbol{\theta}_{\boldsymbol{q}})$. $\xi_{AA}$ and $\mathbf{r}_k$ describes the measurement noise of $I_{AA}$, $I_{DD}$ and $I_{DA}$, respectively. We assume zero-mean Gaussian measurement noise, i.e.:

$$\xi_{AA}(t_k^A) \sim \mathrm{N}\left(0, \sigma_{AA}^2(t_k^D)\right),$$

$$\mathbf{r}_k \sim \mathrm{N}\left(\mathbf{0}, \begin{pmatrix} \sigma_{DD}^2(t_k^D) & 0 \\ 0 & \sigma_{DA}^2(t_k^D) \end{pmatrix}\right).$$

The variances of the measurement noise as functions of time $\sigma_{AA}^2(t_k^A)$, $\sigma_{DD}^2(t_k^D)$, and $\sigma_{DA}^2(t_k^D)$ are determined independently from the B-FRET algorithm (Supplementary Note 3). The measurement model matrix $\mathbf{H}_k(\boldsymbol{\theta}_m)$ is defined as

$$\mathbf{H}_k(\boldsymbol{\theta}_m) = \begin{pmatrix} f_D(t_k^D, \boldsymbol{\theta}_D)[D_{total}] & -f_A(t_k^D, \boldsymbol{\theta}_A) f_D(t_k^D, \boldsymbol{\theta}_D) \\ a f_A(t_k^D, \boldsymbol{\theta}_A)[A_{total}] + d f_D(t_k^D, \boldsymbol{\theta}_D)[D_{total}] & (G-d) f_A(t_k^D, \boldsymbol{\theta}_A) f_D(t_k^D, \boldsymbol{\theta}_D) \end{pmatrix},$$

where $a, d$ and $G$ are imaging-system parameters determined by independent measurements (Online Methods; Supplementary Note 3) and $\boldsymbol{\theta}_m = \{[A_{total}], [D_{total}], \boldsymbol{\theta}_A, \boldsymbol{\theta}_D\}$ are unknown parameters of the measurement matrix $\mathbf{H}_k$ (Online Methods for definitions). We label the set of all unknown parameters in the model by $\boldsymbol{\theta}$, i.e.,

$$\boldsymbol{\theta} = \{\boldsymbol{\theta}_m, \boldsymbol{\theta}_q\}.$$

Both bimolecular and unimolecular FRET systems follow the same model equation, although the interpretations of the parameters and variables are different (Supplementary Note 1).

Making the optimal inference of $E_k \equiv E(t_k^D)$ amounts to computing the posterior probability distribution of $E_k$ given a set of data $\mathcal{D}$ and a model $\mathcal{M}$, $p(E_k|\mathcal{D}, \mathcal{M})$; with the distribution at hand, one can obtain, e.g., the most probable value of $E_k$ quantified by the mode of the distribution and the uncertainty of the estimation quantified by, e.g., the standard deviation of the distribution. As written, this posterior distribution hides the influence of model parameters. To make this explicit, we expand the distribution $p(E_k|\mathcal{D}, \mathcal{M})$ over the model parameters $\boldsymbol{\theta}$:

$$p(E_k|\mathcal{D}, \mathcal{M}) = \int p(E_k, \boldsymbol{\theta}|\mathcal{D}, \mathcal{M})\, d\boldsymbol{\theta}$$

$$= \int p(\boldsymbol{\theta}|\mathcal{D}, \mathcal{M}) p(E_k|\boldsymbol{\theta}, \mathcal{D}, \mathcal{M})\, d\boldsymbol{\theta}. \quad \text{(Eq. 2} - 4)$$

This decomposition illustrates how we evaluate $p(E_k|\mathcal{D}, \mathcal{M})$ in practice: first, evaluate (or draw samples from) the posterior distribution of the model parameters $p(\boldsymbol{\theta}|\mathcal{D}, \mathcal{M})$; second evaluate the posterior distribution of the FRET index $p(E_k|\boldsymbol{\theta}, \mathcal{D}, \mathcal{M})$ given the sampled parameter. With a sufficient number of samples, the integral of the posterior distribution of the FRET index $p(E_k|\mathcal{D}, \mathcal{M})$ is approximated straightforwardly by a Monte Carlo approach (Online methods). In B-FRET, below, we first describe how the respective distributions are evaluated. We then briefly discuss the prior distributions of the parameters.

**Evaluating the posterior distribution of the model parameters** $p(\boldsymbol{\theta}|\mathcal{D}, \mathcal{M})$

Using the Bayes' rule, the posterior distribution of the parameters $\boldsymbol{\theta}$ given the data $\mathcal{D}$, $\log p(\boldsymbol{\theta}|\mathcal{D})$ (hereafter, we omit the conditioning by the model $\mathcal{M}$ to make the expressions less cluttered) can be written as

$$\log p(\boldsymbol{\theta}|\mathcal{D}) = \log p(\mathcal{D}|\boldsymbol{\theta}) + \log p(\boldsymbol{\theta}) + C$$

$$= \log p\left(I_{DD,1:N_D}, I_{DA,1:N_D}\middle|\boldsymbol{\theta}\right) + \log p\left(I_{AA,1:N_A}\middle|\boldsymbol{\theta}\right) + \log p(\boldsymbol{\theta}) + C$$

$$= \log p\left(\mathbf{y}_{1:N_D}\middle|\boldsymbol{\theta}\right) + \log p\left(I_{AA,1:N_A}\middle|\boldsymbol{\theta}\right) + \log p(\boldsymbol{\theta}) + C$$

where $\log p(\boldsymbol{\theta})$ is the log prior distributions of the parameters (see below) and $C$ is the normalization constant. Note that, given the model parameters, $\log p(\mathcal{D}|\boldsymbol{\theta}) = \log p\left(I_{DD,1:N_D}, I_{DA,1:N_D}\middle|\boldsymbol{\theta}\right) + \log p\left(I_{AA,1:N_A}\middle|\boldsymbol{\theta}\right)$ because $I_{AA}$ is independent of the hidden variable $\{\chi_k\}$ and the measurement noise of $I_{AA}$ is independent of that of $I_{DD}$ and $I_{DA}$.

The log-likelihood function of the parameters $\{[A_{total}], \boldsymbol{\theta}_A\}$, $\log p\left(I_{AA,1:N_A}\middle|\boldsymbol{\theta}\right)$ is evaluated as

$$\log p\left(I_{AA,1:N_A}\middle|\boldsymbol{\theta}\right) = \log p\left(I_{AA,1:N_A}\middle|\{[A_{total}],\boldsymbol{\theta}_A\}\right)$$

$$= \log\left(\prod_{k=1}^{N_A}\frac{1}{\sqrt{2\pi\sigma_{AA}^2(t_k^A)}}\exp\left(-\frac{\left(I_{AA}(t_k^A)-f_A(t_k^A,\boldsymbol{\theta}_A)[A_{total}]\right)^2}{2\sigma_{AA}^2(t_k^A)}\right)\right)$$

$$= -\sum_{k=1}^{N_A}\frac{\left(I_{AA}(t_k^A)-f_A(t_k^A,\boldsymbol{\theta}_A)[A_{total}]\right)^2}{2\sigma_{AA}^2(t_k^A)} + \text{Const.} \quad (\text{Eq. }2-5)$$

Evaluating the log-likelihood function $\log p\left(\mathbf{y}_{1:N_D}\middle|\boldsymbol{\theta}\right)$ is less straightforward due to the involvement of the hidden variable $\mathbf{x}_k$. This can be written as

$$\log p\left(\mathbf{y}_{1:N_D}\middle|\boldsymbol{\theta}\right) = \sum_{k=1}^{N_D}\log p(\mathbf{y}_k|\mathbf{y}_{1:k-1},\boldsymbol{\theta})$$

$$= \sum_{k=1}^{N_D}\log\left(\int p(\mathbf{y}_k|\mathbf{x}_k,\boldsymbol{\theta})p(\mathbf{x}_k|\mathbf{y}_{1:k-1},\boldsymbol{\theta})d\mathbf{x}_k\right), \quad (\text{Eq. }2-6)$$

where we define $p(\mathbf{y}_1|\mathbf{y}_{1:0},\boldsymbol{\theta}) \equiv p(\mathbf{y}_1|\boldsymbol{\theta})$. The distribution $p(\mathbf{y}_k|\mathbf{x}_k)$ is specified by the measurement model (Eq. 2-3). Thus, we need to evaluate the predictive distribution of the state at time point $k$, $\mathbf{x}_k$, given certain parameter values, $\boldsymbol{\theta}$, and the observables up to $k-1$, $\mathbf{y}_{1:k-1}$, i.e., $p(\mathbf{x}_k|\mathbf{y}_{1:k-1},\boldsymbol{\theta})$. This can be written as

$$p(\mathbf{x}_k|\mathbf{y}_{1:k-1},\boldsymbol{\theta}) = \int p(\mathbf{x}_k,\mathbf{x}_{k-1}|\mathbf{y}_{1:k-1},\boldsymbol{\theta})d\mathbf{x}_{k-1}$$

$$= \int p(\mathbf{x}_k|\mathbf{x}_{k-1},\mathbf{y}_{1:k-1},\boldsymbol{\theta})p(\mathbf{x}_{k-1}|\mathbf{y}_{1:k-1},\boldsymbol{\theta})d\mathbf{x}_{k-1}$$

$$= \int p(\mathbf{x}_k|\mathbf{x}_{k-1},\boldsymbol{\theta})p(\mathbf{x}_{k-1}|\mathbf{y}_{1:k-1},\boldsymbol{\theta})d\mathbf{x}_{k-1}, \quad (\text{Eq. }2-7)$$

where we used $p(\mathbf{x}_k|\mathbf{x}_{k-1},\mathbf{y}_{1:k-1},\boldsymbol{\theta}) = p(\mathbf{x}_k|\mathbf{x}_{k-1},\boldsymbol{\theta})$. The distribution $p(\mathbf{x}_k|\mathbf{x}_{k-1},\boldsymbol{\theta})$ is specified by the dynamic model (Eq. 2-2). Thus, we need to evaluate the filtering (or posterior) distribution of the hidden state $\mathbf{x}_{k-1}$, given certain parameter values, $\boldsymbol{\theta}$, and the observables up to the same time point $k-1$, $\mathbf{y}_{1:k-1}$, i.e., $p(\mathbf{x}_{k-1}|\mathbf{y}_{1:k-1},\boldsymbol{\theta})$. This can be written, using Bayes' rule, as

$$p(\mathbf{x}_k|\mathbf{y}_{1:k},\boldsymbol{\theta}) = \frac{1}{Z_k}p(\mathbf{y}_k|\mathbf{x}_k,\mathbf{y}_{1:k-1},\boldsymbol{\theta})p(\mathbf{x}_k|\mathbf{y}_{1:k-1},\boldsymbol{\theta})$$

$$= \frac{1}{Z_k}p(\mathbf{y}_k|\mathbf{x}_k,\boldsymbol{\theta})p(\mathbf{x}_k|\mathbf{y}_{1:k-1},\boldsymbol{\theta}), \quad (\text{Eq. }2-8)$$

where we used $p(\mathbf{y}_k|\mathbf{x}_k,\mathbf{y}_{1:k-1}) = p(\mathbf{y}_k|\mathbf{x}_k)$, which is specified by the measurement model (Eq. 2-3). $Z_k$ is the normalization constant.

With these expressions, the likelihood function (Eq. 2-6) can be evaluated by the following recursive method. First, by providing the filtering distribution of the time point one step before the initial time point $p(\mathbf{x}_0|\boldsymbol{\theta})$ $\left(= p(\mathbf{x}_0|\mathbf{y}_{1:0},\boldsymbol{\theta})\right)$ as a prior, Eq. 2-7 gives the predictive distribution of the initial time point $p(\mathbf{x}_1|\mathbf{y}_{1:0},\boldsymbol{\theta}) = p(\mathbf{x}_1|\boldsymbol{\theta})$. Given this predictive distribution, Eq. 2-8 gives the filtering distribution of the initial time point $p(\mathbf{x}_1|\mathbf{y}_1,\boldsymbol{\theta})$. This enables to evaluate, through Eq. 2-7, the predictive distribution of the next time point $p(\mathbf{x}_2|\mathbf{y}_1,\boldsymbol{\theta})$, which can be fed into Eq. 2-8 to obtain the filtering distribution $p(\mathbf{x}_2|\mathbf{y}_2, \boldsymbol{\theta})$. By repeating this procedure, the predictive distributions $p(\mathbf{x}_k|\mathbf{y}_{1:k-1}, \boldsymbol{\theta})$ at all the following time points can be obtained. This enables to compute the likelihood function (Eq. 2-6).

The predictive and filtering distributions have closed-form expressions when i) the model is linear, which is true for FRET measurements, and ii) the process noise is Gaussian. Since Gaussian dynamic model is able to capture a broad range of dynamics[1,2], and therefore has direct relevance to FRET-data analysis, we first discuss the linear-Gaussian case below. However, there can be situations where the dynamics of a system is inherently non-Gaussian (e.g., the step-like dynamics described in the main text). To better capture the dynamics in such cases, one needs to assume a non-Gaussian process noise, which necessitates evaluating the predictive and filtering distributions numerically. We thus discuss the non-Gaussian case next.

<u>Gaussian process noise</u>

Assuming Gaussian process noise, the predictive and filtering distributions can be written as[1,2]

$$p(\mathbf{x}_k|\mathbf{y}_{1:k-1}, \boldsymbol{\theta}) = \mathrm{N}\big(\mathbf{x}_k \big| \mathbf{m}_{\boldsymbol{k}}^{-}(\boldsymbol{\theta}), \mathbf{P}_k^{-}(\boldsymbol{\theta})\big),$$

$$p(\mathbf{x}_k|\mathbf{y}_{1:k}, \boldsymbol{\theta}) = \mathrm{N}\big(\mathbf{x}_k \big| \mathbf{m}_{\boldsymbol{k}}(\boldsymbol{\theta}), \mathbf{P}_k(\boldsymbol{\theta})\big),$$

where the parameters of the Gaussian distributions above can be computed with the following Kalman filter prediction and update steps.

The prediction step is

$$\mathbf{m}_{\boldsymbol{k}}^{-}(\boldsymbol{\theta}) = \mathbf{m}_{k-1}(\boldsymbol{\theta}),$$

$$\mathbf{P}_k^{-}(\boldsymbol{\theta}) = \mathbf{P}_{k-1}(\boldsymbol{\theta}) + \mathbf{Q}_{k-1},$$

where $\mathbf{Q}_{k-1}$ is the variance-covariance matrix of the process noise, i.e., $\mathbf{q}_{k-1} \sim \mathrm{N}(\mathbf{0}, \mathbf{Q}_{k-1})$, and $\mathbf{Q}_{k-1} = \begin{pmatrix} 0 & 0 \\ 0 & \sigma_\chi^2 \end{pmatrix}$.

The update step is

$$\begin{aligned}
\mathbf{v}_k(\boldsymbol{\theta}) &= \mathbf{y}_k - \mathbf{H}_k(\boldsymbol{\theta})\mathbf{m}_k^{-}(\boldsymbol{\theta}),\\
\mathbf{S}_k(\boldsymbol{\theta}) &= \mathbf{H}_k(\boldsymbol{\theta})\mathbf{P}_k^{-}(\boldsymbol{\theta})\mathbf{H}_k^{\mathrm{T}}(\boldsymbol{\theta}) + \mathbf{R}_k,\\
\mathbf{K}_k(\boldsymbol{\theta}) &= \mathbf{P}_k^{-}(\boldsymbol{\theta})\mathbf{H}_k^{\mathrm{T}}(\boldsymbol{\theta})\mathbf{S}_{\boldsymbol{k}}^{-\mathbf{1}}(\boldsymbol{\theta}),\\
\mathbf{m}_{\boldsymbol{k}}(\boldsymbol{\theta}) &= \mathbf{m}_k^{-}(\boldsymbol{\theta}) + \mathbf{K}_k(\boldsymbol{\theta})\mathbf{v}_k(\boldsymbol{\theta}),\\
\mathbf{P}_k(\boldsymbol{\theta}) &= \mathbf{P}_k^{-}(\boldsymbol{\theta}) - \mathbf{K}_k(\boldsymbol{\theta})\mathbf{S}_k(\boldsymbol{\theta})\mathbf{K}_k^{\mathrm{T}}(\boldsymbol{\theta}).
\end{aligned}$$

The recursion is started from the prior mean $\mathbf{m}_0$ and covariance $\mathbf{P}_0$ of the distribution of $\mathbf{x}_0$ $p(\mathbf{x}_0|\mathbf{y}_{1:0}, \boldsymbol{\theta}) \equiv \mathrm{N}(\mathbf{x}_0|\mathbf{m}_0,\ \mathbf{P}_0)$, which is given as

$$\mathbf{m}_0 = \begin{pmatrix} 1 \\ \chi_0 \end{pmatrix},$$

$$\mathbf{P}_0 = \begin{pmatrix} \epsilon & 0 \\ 0 & \sigma_{\chi_0}^2 \end{pmatrix}.$$

The parameters $\chi_0$ and $\sigma_{\chi_0}$ are chosen to reflect our ignorance about the initial state. Typically, an arbitrary value of $\chi_0$ (e.g., $\chi_0 = 0$) and a sufficiently large variance $\sigma_{\chi_0}^{\mathbf{2}}$ are used. The parameter $\epsilon$ ($>$

0) needs to be sufficiently small but nonzero, e.g., $\epsilon = 10^{-10}$, for the numerical stability in computing $\mathbf{P}_0^{-1}(\boldsymbol{\theta})$. The exact choices of these parameters do not affect the result.

Using these, the log posterior distribution (Eq. 2-6) can be written as

$$\log p(\mathbf{y}_{1:N_D}|\boldsymbol{\theta}) = \sum_{k=1}^{N_D} \log\left(\int p(\mathbf{y}_k|\mathbf{x}_\mathrm{k}) p(\mathbf{x}_\mathrm{k}|\mathbf{y}_{1:k-1},\boldsymbol{\theta}) d\mathbf{x}_k\right)$$

$$= \sum_{k=1}^{N_D} \log\left(\int \mathrm{N}(\mathbf{y}_k|\mathbf{H}_k(\boldsymbol{\theta})\mathbf{x}_k, \mathbf{R}_k)\mathrm{N}\left(\mathbf{x}_k|\mathbf{m}_k^-(\boldsymbol{\theta}), \mathbf{P}_k^-(\boldsymbol{\theta})\right) d\mathbf{x}_k\right)$$

$$= \sum_{k=1}^{N_D} \log \mathrm{N}\left(\mathbf{y}_k\middle|\mathbf{H}_k(\boldsymbol{\theta})\mathbf{m}_k^-(\boldsymbol{\theta}), \mathbf{S}_k(\boldsymbol{\theta})\right)$$

$$= -\sum_{k=1}^{N_D}\left(\frac{1}{2}\log|2\pi\mathbf{S}_k(\boldsymbol{\theta})| + \frac{1}{2}\mathbf{v}_k^\mathrm{T}(\boldsymbol{\theta})\mathbf{S}_k^{-1}(\boldsymbol{\theta})\mathbf{v}_k(\boldsymbol{\theta})\right).$$

Thus, the posterior distribution of the parameters is given by

$$\log p(\boldsymbol{\theta}|\mathcal{D}) = \log p\left(\mathbf{y}_{1:N_D}|\boldsymbol{\theta}\right) + \log p\left(I_{AA,1:N_A}|\boldsymbol{\theta}\right) + \log p(\boldsymbol{\theta}) + \mathrm{Const.}$$

$$= -\sum_{k=1}^{N_D}\left(\frac{1}{2}\log|2\pi\mathbf{S}_k(\boldsymbol{\theta})| + \frac{1}{2}\mathbf{v}_k^T(\boldsymbol{\theta})\mathbf{S}_k^{-1}(\boldsymbol{\theta})\mathbf{v}_k(\boldsymbol{\theta})\right) - \sum_{k=1}^{N_A}\frac{(I_{AA}(t_k^A) - f_A(t_k^A, \boldsymbol{\theta}_A)[A_{total}])^2}{2\sigma_{AA}^2(t_k^A)} + \log p(\boldsymbol{\theta}) + \mathrm{Const.} \quad (\mathrm{Eq.}\,2-9)$$

<u>Non-Gaussian process noise</u>

To compute the filtering and predictive distributions numerically, we approximate functions by a step function[1]. Since the first element of $\mathbf{x}_\mathrm{k}$ is fixed to 1, the predictive distribution in practice is one-dimensional, i.e., $p\left(\mathbf{x}_\mathrm{k} = (1, \chi_k)^\mathrm{T}\middle|\mathbf{y}_{1:k-1},\boldsymbol{\theta}\right) = p(\chi_k|\mathbf{y}_{1:k-1},\boldsymbol{\theta})$. Rewriting the predictive distribution, we have

$$p(\chi_k|\mathbf{y}_{1:k-1},\boldsymbol{\theta}) = \int p(\chi_k|\chi_{k-1},\boldsymbol{\theta})p(\chi_{k-1}|\mathbf{y}_{1:k-1},\boldsymbol{\theta})d\chi_{k-1}.$$

To approximate the distribution $p(\chi_k|\mathbf{y}_{1:k-1},\boldsymbol{\theta})$ by a step function, we first restrict the domain of the function to a finite interval $x_0 \le \chi_k \le x_d$, where $x_0$ and $x_d$ are sufficiently small and large numbers respectively. Then we divide the interval into d sub-intervals $x_0 < x_1 < \cdots < x_d$ with a uniform interval $\Delta x = x_{i+1} - x_i$ for $i = 0, \ldots, d-1$. The predictive distribution $p(\chi_k|\mathbf{y}_{1:k-1},\boldsymbol{\theta}) \equiv \tilde{p}(\chi_k)$ is then specified by $\{x_1, \ldots, x_d; \tilde{p}_1, \ldots, \tilde{p}_d\}$, where $\tilde{p}_i = \tilde{p}(x_i)$ for $i = 1, \ldots, d$. In the same way, the filtering distribution $p(\chi_{k-1}|\mathbf{y}_{1:k-1},\boldsymbol{\theta}) \equiv \tilde{f}(\chi_{k-1})$ is specified by $\{x_1, \ldots, x_d; \tilde{f}_1, \ldots, \tilde{f}_d\}$, where $\tilde{f}_i = \tilde{f}(x_i)$. The process noise $q_{k-1}$ that appears in the dynamic model, $\chi_k = \chi_{k-1} + q_{k-1}$, follows a distribution $p(q|\boldsymbol{\theta}) \equiv \tilde{Q}(q)$, and is specified, with the same discretization interval $\Delta x$ and a sufficiently large number $x_D = x_d - x_0$, by $\{x_{-D}, \ldots, x_D; \tilde{Q}_{-D}, \ldots, \tilde{Q}_D\}$, where $\tilde{Q}_i = \tilde{Q}(x_i)$ for $i = -D, -D+1, \ldots, D$. Using this notation, the approximated prediction distribution can be written as, for $i = 1, \ldots, d$,

$$\tilde{p}_i = \tilde{p}(x_i) = \int_{x_0}^{x_d} \tilde{Q}(x_i - x)\tilde{f}(x)dx$$

$$= \sum_{j=1}^{d} \int_{x_{j-1}}^{x_j} \tilde{Q}(x_i - x)\, \tilde{f}(x) dx$$

$$\approx \Delta x \sum_{j=1}^{d} \tilde{Q}_{i-j} \tilde{f}_j$$

$$= \Delta x \big[\tilde{\mathbf{f}}\tilde{\mathbf{Q}}\big]_i, \quad (\text{Eq. } 2-10)$$

where $\tilde{\mathbf{f}} = (\tilde{f}_1, \tilde{f}_2, \dots, \tilde{f}_d)$ and $\tilde{\mathbf{Q}} = \begin{pmatrix} \tilde{Q}_0 & \cdots & \tilde{Q}_{d-1} \\ \vdots & \ddots & \vdots \\ \tilde{Q}_{1-d} & \cdots & \tilde{Q}_0 \end{pmatrix}$. After this is computed, $\tilde{p}_i$ rescaled to $\frac{\tilde{p}_i}{\Delta x (\sum_{l=1}^{d} \tilde{p}_l)}$ to properly normalize the approximated distribution.

Eq. 2 – 10 approximates the predictive distribution at time $t_k$, $p(\chi_k | \mathbf{y}_{1:k-1}, \boldsymbol{\theta})$, as a piecewise continuous function. Given this, the filtering distribution at time $t_k$,

$$p(\chi_k | \mathbf{y}_{1:k}, \boldsymbol{\theta}) = \frac{1}{Z_k} p(\mathbf{y}_k | \chi_k, \boldsymbol{\theta}) p(\chi_k | \mathbf{y}_{1:k-1}, \boldsymbol{\theta})$$

is approximated by multiplying the discrete values $\tilde{p}_i$ by $p(\mathbf{y}_k | \chi_k, \boldsymbol{\theta}) = \mathrm{N}(\mathbf{y}_k | \mathbf{H}_k(\boldsymbol{\theta})(1, \chi_k)^{\mathrm{T}}, \mathbf{R}_k)$, evaluated at the same timepoints. Thus, the values of the steps in filtering distribution are:

$$\tilde{f}_i = \tilde{f}(x_i) = \frac{\mathrm{N}(\mathbf{y}_k | \mathbf{H}_k(\boldsymbol{\theta})(1, x_i)^{\mathrm{T}}, \mathbf{R}_k) \tilde{p}_i}{C}, \quad (\text{Eq. } 2-11)$$

where the normalization constant $C$ is given by

$$C = \int_{x_0}^{x_d} \mathrm{N}\big(\mathbf{y}_k \big| \mathbf{H}_k(\boldsymbol{\theta})(1, x_i)^{\mathrm{T}}, \mathbf{R}_k\big) \tilde{p}(x_i) dx_i$$

$$= \sum_{j=1}^{d} \int_{x_{j-1}}^{x_j} \mathrm{N}\big(\mathbf{y}_k \big| \mathbf{H}_k(\boldsymbol{\theta})(1, x_i)^{\mathrm{T}}, \mathbf{R}_k\big) \tilde{p}(x_i) dx_i$$

$$= \Delta x \sum_{j=1}^{d} \mathrm{N}\left(\mathbf{y}_k \middle| \mathbf{H}_k(\boldsymbol{\theta})(1, x_j)^{\mathrm{T}}, \mathbf{R}_k\right) \tilde{p}_j.$$

In the same way as the Gaussian case, the predictive and filtering distributions at all time points can be computed in a recursive manner.

Once we have the predictive distribution at each time point $t = t_k^D$, we can evaluate the log-likelihood function as

$$\log p\big(\mathbf{y}_{1:N_D} \big| \boldsymbol{\theta}\big) = \sum_{k=1}^{N_D} \log\left(\int p(\mathbf{y}_k | \chi_k, \boldsymbol{\theta}) p(\chi_k | \mathbf{y}_{1:k-1}, \boldsymbol{\theta}) d\chi_k\right)$$

$$= \sum_{k=1}^{N_D} \log\left(\int_{x_0}^{x_d} \mathrm{N}\big(\mathbf{y}_k \big| \mathbf{H}_k(\boldsymbol{\theta})(1, x)^{\mathrm{T}}, \mathbf{R}_k\big) \tilde{p}(x) dx\right)$$

$$= \sum_{k=1}^{N_D} \log\left( \sum_{j=1}^{d} \int_{x_{j-1}}^{x_j} \mathrm{N}\left(\mathbf{y}_k \middle| \mathbf{H}_k(\boldsymbol{\theta})(1,x)^{\mathrm{T}}, \mathbf{R}_k\right) \tilde{p}(x)\, dx \right)$$

$$= \sum_{k=1}^{N_D} \log\left( \Delta x \sum_{j=1}^{d} \mathrm{N}\left(\mathbf{y}_k \middle| \mathbf{H}_k(\boldsymbol{\theta})\left(1,x_j\right)^{\mathrm{T}}, \mathbf{R}_k\right) \tilde{p}_j \right). \quad (\mathrm{Eq.}\,2-12)$$

Since the likelihood function $p\left(I_{AA,1:N_A} \middle| \boldsymbol{\theta}\right)$ and the prior distribution $p(\boldsymbol{\theta})$ can be evaluated in the same way as the Gaussian case, the posterior distribution of the parameters is given by

$$\log p(\boldsymbol{\theta}|\mathcal{D}) = \log p\left(\mathbf{y}_{1:N_D} \middle| \boldsymbol{\theta}\right) + \log p\left(I_{AA,1:N_A} \middle| \boldsymbol{\theta}\right) + \log p(\boldsymbol{\theta}) + C$$

$$= \sum_{k=1}^{N_D} \log\left( \Delta x \sum_{j=1}^{d} \mathrm{N}\left(\mathbf{y}_k \middle| \mathbf{H}_k(\boldsymbol{\theta})\left(1,x_j\right)^{\mathrm{T}}, \mathbf{R}_k\right) \tilde{p}_j \right) - \sum_{k=1}^{N_A} \frac{\left(I_{AA}(t_k^A) - f_A(t_k^A, \boldsymbol{\theta}_A)[A_{total}]\right)^2}{2\sigma_{AA}^2(t_k^A)} + \log p(\boldsymbol{\theta}) + \mathrm{Const.}\,(\mathrm{Eq.}\,2-13)$$

**Approximating the posterior distribution** $\log p(\boldsymbol{\theta}|\mathcal{D})$

Samples $\{\boldsymbol{\theta}_i\}$ can be drawn from this posterior distribution Eq. 2-9 or Eq. 2-13, by using a Markov chain Monte Carlo (MCMC) method (e.g., Slice sampling[3,4]) – this approach is exact in the limit of large number of samples. However, since this method is computationally costly, in the B-FRET algorithm, we implemented the option to approximate the distribution by a lognormal distribution (the Laplace approximation[3,4]); we chose a lognormal distribution rather than a Gaussian distribution since all parameters in our model take positive values. To make this approximation, we first find the mode of the log posterior distribution as a function of $\log p\ (\log \boldsymbol{\theta}\ |\mathcal{D})$, which we label by $\log \boldsymbol{\theta}_{MAP}$, using an optimization algorithm, and then compute the Hessian matrix at the mode defined by

$$\mathbf{A} = -\nabla_{\log \boldsymbol{\theta}} \nabla_{\log \boldsymbol{\theta}} \log p(\log \boldsymbol{\theta}\ |\mathcal{D})\ |_{\log \boldsymbol{\theta} = \log \boldsymbol{\theta}_{MAP}}.$$

Them, the posterior distribution is approximated by

$$p(\boldsymbol{\theta}|\mathcal{D}) \approx \mathrm{Lognormal}(\boldsymbol{\theta}|\log \boldsymbol{\theta}_{MAP}, \mathbf{A}^{-1}),$$

where $\mathbf{A}^{-1}$ is the inverse of $\mathbf{A}$. The difference in the performance between the two methods are negligible (Supplementary Fig. 4), so we used the Laplace approximation unless otherwise indicated.

**Evaluating the posterior distribution of the state given a set of parameters** $p(E_k|\boldsymbol{\theta}, \mathcal{D}, \mathcal{M})$

Since a user-defined FRET index $E_k$ is dependent only on the hidden state $\mathbf{x}_k$ and model parameters $\boldsymbol{\theta}$ (e.g., $E_k = \frac{\chi_k}{[D_{total}]} = E_{max} \frac{[DA_{total}]}{[D_{total}]}$, where $[D_{total}]$ is a part of the model parameter $\boldsymbol{\theta}$), evaluating the (smoothing) distribution of the FRET index conditioned by parameters $p(E_k|\boldsymbol{\theta}, \mathcal{D})$ is equivalent to evaluating the distribution of the hidden state conditioned by the parameters $p(\mathbf{x}_k|\boldsymbol{\theta}, \mathcal{D})$. Note that, once the parameters $\boldsymbol{\theta}$ are given, the only relevant data to the inference of $\mathbf{x}_k$ is $\mathbf{y}_{1:N_D}$ since $I_{AA}$ is independent from $\{\mathbf{x}_k\}$, and thus $p(\mathbf{x}_k|\boldsymbol{\theta}, \mathcal{D}) = p\left(\mathbf{x}_k \middle| \boldsymbol{\theta}, \mathbf{y}_{1:N_D}\right)$. The distribution $p\left(\mathbf{x}_k \middle| \boldsymbol{\theta}, \mathbf{y}_{1:N_D}\right)$ can be evaluated by the following Bayesian smoothing equation[1,2]:

$$p\left(\mathbf{x}_k \middle| \boldsymbol{\theta}, \mathbf{y}_{1:N_D}\right) = \int p\left(\mathbf{x}_k, \mathbf{x}_{k+1} \middle| \mathbf{y}_{1:N_D}, \boldsymbol{\theta}\right) d\mathbf{x}_{k+1}$$

$$= \int p(\mathbf{x}_{k+1}|\mathbf{y}_{1:N_D}, \boldsymbol{\theta}) p(\mathbf{x}_k|\mathbf{x}_{k+1}, \mathbf{y}_{1:N_D}, \boldsymbol{\theta}) \, d\mathbf{x}_{k+1}$$

$$= \int p(\mathbf{x}_{k+1}|\mathbf{y}_{1:N_D}, \boldsymbol{\theta}) p(\mathbf{x}_k|\mathbf{x}_{k+1}, \mathbf{y}_{1:k}, \boldsymbol{\theta}) \, d\mathbf{x}_{k+1}$$

$$= \int p(\mathbf{x}_{k+1}|\mathbf{y}_{1:N_D}, \boldsymbol{\theta}) \frac{p(\mathbf{x}_k|\mathbf{y}_{1:k}, \boldsymbol{\theta}) p(\mathbf{x}_{k+1}|\mathbf{x}_k, \mathbf{y}_{1:k}, \boldsymbol{\theta})}{p(\mathbf{x}_{k+1}|\mathbf{y}_{1:k}, \boldsymbol{\theta})} d\mathbf{x}_{k+1}$$

$$= p(\mathbf{x}_k|\mathbf{y}_{1:k}, \boldsymbol{\theta}) \int \frac{p(\mathbf{x}_{k+1}|\mathbf{y}_{1:N_D}, \boldsymbol{\theta}) p(\mathbf{x}_{k+1}|\mathbf{x}_k, \boldsymbol{\theta})}{p(\mathbf{x}_{k+1}|\mathbf{y}_{1:k}, \boldsymbol{\theta})} d\mathbf{x}_{k+1} \quad (\text{Eq. } 2-14),$$

where from the second to the third line we used

$$p(\mathbf{x}_k|\mathbf{x}_{k+1}, \mathbf{y}_{1:N_D}, \boldsymbol{\theta}) = p(\mathbf{x}_k|\mathbf{x}_{k+1}, \mathbf{y}_{1:k}, \mathbf{y}_{k+1:N_D}, \boldsymbol{\theta})$$

$$= \frac{p(\mathbf{y}_{k+1:N_D}|\mathbf{x}_k, \mathbf{x}_{k+1}, \mathbf{y}_{1:k}, \boldsymbol{\theta}) p(\mathbf{x}_k|\mathbf{x}_{k+1}, \mathbf{y}_{1:k}, \boldsymbol{\theta})}{p(\mathbf{y}_{k+1:N_D}|\mathbf{x}_{k+1}, \mathbf{y}_{1:k}, \boldsymbol{\theta})}$$

$$= \frac{p(\mathbf{y}_{k+1:N_D}|\mathbf{x}_{k+1}, \mathbf{y}_{1:k}, \boldsymbol{\theta}) p(\mathbf{x}_k|\mathbf{x}_{k+1}, \mathbf{y}_{1:k}, \boldsymbol{\theta})}{p(\mathbf{y}_{k+1:N_D}|\mathbf{x}_{k+1}, \mathbf{y}_{1:k}, \boldsymbol{\theta})}$$

$$= p(\mathbf{x}_k|\mathbf{x}_{k+1}, \mathbf{y}_{1:k}, \boldsymbol{\theta}).$$

The Bayesian smoothing equation (Eq. 2-14) has a closed form expression when the process noise is Gaussian. However, when the process noise is non-Gaussian, we need to resort to a numerical method. We discuss both cases below.

Gaussian process noise

The closed-form expression[1,2] for the smoothed distribution is

$$p(\mathbf{x}_k|\mathbf{y}_{1:N_D}, \boldsymbol{\theta}) = \mathrm{N}(\mathbf{x}_k|\mathbf{m}_k^{\mathrm{s}}(\boldsymbol{\theta}), \mathbf{P}_k^{\mathrm{s}}(\boldsymbol{\theta})),$$

where the parameters of the Gaussian distributions can be computed by the following RTS(Rauch-Tung-Striebel) smoother:

$$\mathbf{m}_{k+1}^{-}(\boldsymbol{\theta}) = \mathbf{m}_k(\boldsymbol{\theta}),$$

$$\mathbf{P}_{k+1}^{-}(\boldsymbol{\theta}) = \mathbf{P}_k(\boldsymbol{\theta}) + \mathbf{Q}_k,$$

$$\mathbf{G}_k(\boldsymbol{\theta}) = \mathbf{P}_k(\boldsymbol{\theta})[\mathbf{P}_{k+1}^{-}(\boldsymbol{\theta})]^{-1},$$

$$\mathbf{m}_k^{\mathrm{s}}(\boldsymbol{\theta}) = \mathbf{m}_k(\boldsymbol{\theta}) + \mathbf{G}_k(\boldsymbol{\theta})[\mathbf{m}_{k+1}^{\mathrm{s}}(\boldsymbol{\theta}) - \mathbf{m}_{k+1}^{-}(\boldsymbol{\theta})],$$

$$\mathbf{P}_k^{s}(\boldsymbol{\theta}) = \mathbf{P}_k(\boldsymbol{\theta}) + \mathbf{G}_k(\boldsymbol{\theta})[\mathbf{P}_{k+1}^{\mathrm{s}}(\boldsymbol{\theta}) - \mathbf{P}_{k+1}^{-}(\boldsymbol{\theta})]\mathbf{G}_k^{\mathrm{T}}(\boldsymbol{\theta}).$$

Here, $\mathbf{m}_k^{-}(\boldsymbol{\theta})$, $\mathbf{P}_k^{-}(\boldsymbol{\theta})$, $\mathbf{m}_k(\boldsymbol{\theta})$, and $\mathbf{P}_k(\boldsymbol{\theta})$ are the mean and covariance of the predictive and smoothed distribution computed above, i.e., $p(\mathbf{x}_k|\mathbf{y}_{1:k-1}, \boldsymbol{\theta}) = \mathrm{N}(\mathbf{x}_k|\mathbf{m}_k^{-}(\boldsymbol{\theta}), \mathbf{P}_k^{-}(\boldsymbol{\theta}))$ and $p(\mathbf{x}_k|\mathbf{y}_{1:k}, \boldsymbol{\theta}) = \mathrm{N}(\mathbf{x}_k|\mathbf{m}_k(\boldsymbol{\theta}), \mathbf{P}_k(\boldsymbol{\theta}))$. $\mathbf{Q}_k$ is the variance-covariance matrix of the process noise, i.e., $\mathbf{q}_k \sim \mathrm{N}(\mathbf{0}, \mathbf{Q}_k)$. The recursion is initiated from the last timepoint $k = N_D$, with $\mathbf{m}_{N_D}^{\mathrm{s}}(\boldsymbol{\theta}) = \boldsymbol{m}_{N_D}(\boldsymbol{\theta})$ and $\mathbf{P}_{N_D}^{\mathrm{s}}(\boldsymbol{\theta}) = \mathbf{P}_{N_D}(\boldsymbol{\theta})$.

Non-Gaussian process noise

We approximate the smoothed distribution $p(\mathbf{x}_k|\mathbf{y}_{1:N_D})$ by step functions[1]. Rewriting the smoothed distribution using $\chi(t_{D,k}) \equiv \chi_k$, we get

$$p(\chi_k|\mathbf{y}_{1:N_D},\boldsymbol{\theta}) = p(\chi_k|\mathbf{y}_{1:k},\boldsymbol{\theta}) \int \frac{p(\chi_{k+1}|\mathbf{y}_{1:N_D},\boldsymbol{\theta})p(\chi_{k+1}|\chi_k,\boldsymbol{\theta})}{p(\chi_{k+1}|\mathbf{y}_{1:k},\boldsymbol{\theta})} d\chi_{k+1}.$$

In the same way as the step-function approximation of the predictive and filtering distributions, we restrict the domain of the function to a finite interval $x_0 \le \chi_k \le x_d$, and divide the interval into d sub-intervals $x_0 < x_1 < \cdots < x_d$ with a uniform interval $\Delta x = x_{i+1} - x_i$ for $i = 1, \dots, d-1$ . The smoothed distribution $p(\chi_k|\mathbf{y}_{1:N_D},\boldsymbol{\theta}) \equiv \tilde{s}(\chi_k)$ is then specified by $\{x_0, \dots, x_d; \tilde{s}_1, \dots, \tilde{s}_d\}$, where $\tilde{s}_i = \tilde{s}(x_i)$. Using this notation, the approximated smoothed distribution can be written as, for $i = 1, \dots, d$,

$$\begin{aligned}
\tilde{s}_i = \tilde{s}(x_i) &= \tilde{f}(x_i) \int_{x_0}^{x_d} \frac{\tilde{s}(x)\tilde{Q}(x_i - x)}{\tilde{p}(x)} dx \\
&= \tilde{f}(x_i) \sum_{j=1}^{d} \int_{x_{j-1}}^{x_j} \frac{\tilde{s}(x)\tilde{Q}(x_i - x)}{\tilde{p}(x)} dx \\
&= \Delta x \tilde{f}_i \sum_{j=1}^{d} \tilde{Q}_{i-j} \frac{\tilde{s}_j}{\tilde{p}_j} \\
&= \Delta x \tilde{f}_i \big[\tilde{\boldsymbol{S}}\tilde{\boldsymbol{Q}}\big]_i,
\end{aligned}$$

where $\tilde{\mathbf{S}} = \left(\frac{\tilde{s}_1}{\tilde{p}_1}, \frac{\tilde{s}_2}{\tilde{p}_2}, \dots, \frac{\tilde{s}_d}{\tilde{p}_d}\right)$ and $\tilde{\mathbf{Q}} = \begin{pmatrix} \tilde{Q}_0 & \cdots & \tilde{Q}_{d-1} \\ \vdots & \ddots & \vdots \\ \tilde{Q}_{1-d} & \cdots & \tilde{Q}_0 \end{pmatrix}$. The predictive distribution $\{\tilde{p}_i\}$ and filtering distribution $\{\tilde{f}_i\}$ and are obtained by the recursive filtering algorithm described above. After this is computed, $s_i$ is modified to $\frac{s_i}{\Delta x(\sum_{l=1}^{d} s_l)}$ to normalize the approximated distribution.

**Prior distributions of the model parameters**

The choice of prior distributions for the model parameters $\boldsymbol{\theta} = \{[A_{total}], [D_{total}], \boldsymbol{\theta}_A, \boldsymbol{\theta}_D, \boldsymbol{\theta}_q\}$ depends on the detail of the FRET experiment and how much an experimenter has knowledge about the parameters in advance. As demonstrated in the main text, however, it is not necessary to know the values of these parameters in advance because the fluorescence time series obtained by a typical FRET measurement contain enough information to confine these parameters. Here, we discuss some examples and practical tips on constructing priors, without assuming any knowledge about these parameter values.

First, for a parameter whose value is restricted within a range of $[a, b]$ by definition (e.g., the wight parameter $\delta$ in the bi-exponential photobleaching function, $f_D(t) = \delta \exp\left(-\frac{t}{\tau_{D1}}\right) + (1 - \delta) \exp\left(-\frac{t}{\tau_{D2}}\right)$, is bounded by [0,1]), one can use, e.g., a uniform distribution $p(x|a, b)$, which takes $\frac{1}{b-a}$ for $x \in [a, b]$ and 0 otherwise.

Other parameters that appear in the model can only take positive values, i.e., left-bounded by zero, by definition (e.g., the total concentration of acceptor $[A_{total}]$). For those parameters, we used log-normal distributions:

$$p(x) = \mathrm{lognormal}(x|\mu,\sigma^2) = \frac{1}{x\sigma\sqrt{2\pi}}\exp\left(-\frac{(\ln x-\mu)^2}{2\sigma^2}\right).$$

Now we discuss how we chose $\mu$ and $\sigma^2$ for each parameter. The basic idea here is that before executing the B-FRET algorithm we roughly estimate each parameter and set $\mu$ and $\sigma^2$ so that "true" value of the parameter is certainly included in the support of the prior function.

First, $[A_{total}]$ and $\boldsymbol{\theta}_A$ can be estimated relatively precisely without B-FRET because $I_{AA}$ is not dependent on the hidden variable $\{\chi_k\}$ (see Eq. 2-1). The estimation is done by simply fitting the equation for $I_{AA}$, $I_{AA} = [A_{total}]f_A(t|\boldsymbol{\theta}_A)$ (Eq. 2-1; here we neglect the measurement-noise term), to the data $\{I_{AA}(t_k^A)\}$. Namely,

$$\{[A_{total}]_{est}, \boldsymbol{\theta}_{A,est}\} = \underset{\{[A_{total}],\boldsymbol{\theta}_A\}}{\mathrm{argmin}} \sum_{k=1}^{N_A}\left([A_{total}]f_A(t|\boldsymbol{\theta}_A) - I_{AA}(t_k^A)\right)^2.$$

Thus, for the priors of $[A_{total}]$ and $\boldsymbol{\theta}_A$, we chose $\mu$ and $\sigma$ such that the mode of the log-normal distribution ($\exp(\mu-\sigma^2)$) matches the estimated parameter values, after manually selecting relatively small $\sigma=\sigma_0$ to reflect our relatively high confidence about the estimated parameter value. Note the standard deviation of a log-normal distribution $\mathrm{lognormal}(x|\mu,\sigma^2)$ with respect to $\log x$ is $\sigma$, so if one thinks an estimated parameter could be off roughly by $X_{err}$ fold, s/he can set, e.g., $\sigma = \log(X_{err})$. Thus, the priors for these parameters are:

$$p([A_{total}]) = \mathrm{Lognormal}([A_{total}]\,|\log[A_{total}]_{est} + \sigma_0^2, \sigma_0^2),$$

$$p(\boldsymbol{\theta}_{A,i}) = \mathrm{Lognormal}(\boldsymbol{\theta}_{A,i}\,|\log\boldsymbol{\theta}_{A,est,i} + \sigma_0^2, \sigma_0^2),$$

where $\boldsymbol{\theta}_{A,i}$ is the i-th element of $\boldsymbol{\theta}_A$.

The parameters $[D_{total}]$ and $\boldsymbol{\theta}_D$ cannot be estimated in the same way because both $I_{DD}$ and $I_{DA}$ are dependent on the hidden variable $\{\chi_k\}$, which one cannot access before applying B-FRET. To roughly estimate $[D_{total}]$, we note that the equations for $I_{DD}$ and $I_{DA}$ can be written, neglecting the measurement-noise term, as

$$I_{DD}(t_k^D) \approx f_D(t_k^D,\boldsymbol{\theta}_D)[D_{total}] + -f_A(t_k^D,\boldsymbol{\theta}_A)f_D(t_k^D,\boldsymbol{\theta}_D)\chi_k,$$

$$I_{DA}(t_k^D) \approx a[A_{total}]f_A(t_k^D,\boldsymbol{\theta}_D) + d[D_{total}]f_D(t_k^D,\boldsymbol{\theta}_D) + (G-d)f_A(t_k^D,\boldsymbol{\theta}_A)f_D(t_k^D,\boldsymbol{\theta}_D)\chi_k.$$

At the first time point of a measurement, $f_D(t_1^D,\boldsymbol{\theta}_D) = f_A(t_1^D,\boldsymbol{\theta}_A) = 1$ by definition (there's no photobleaching), and $I_{AA}(t_1^A) \approx [A_{total}]$. Deleting $\chi_k$ from the two equations, we get

$$[D_{total}]_{est} \approx \frac{I_{DA}(t_1^D) + \left((G-d)I_{DD}(t_1^D) - aI_{AA}(t_1^A)\right)}{G}.$$

The right-hand side of this equations are all observables ($I_{AA}(t_1^A)$, $I_{DD}(t_1^D)$, and $I_{DA}(t_1^D)$) or imaging-system parameters ($a, d$, and $G$), and thus computable without knowing $\{\chi_k\}$. Using this, we set the prior of $[D_{total}]$ as

$$p([D_{total}]) = \mathrm{Lognormal}([D_{total}]\,|\log[D_{total}]_{est} + \sigma_0^2, \sigma_0^2).$$

Since $[D_{total}]_{est}$ only gives a crude estimation, we set $\sigma_0^2$ to a relatively high value. To roughly estimate $\boldsymbol{\theta}_D$, we fit the function of $f_D(t|\boldsymbol{\theta}_D)$ to a normalized time series of $I_{DD}$, $\{I_{DD}(t_k^D)/I_{DD}(t_1^D)\}$, i.e.,

$$\boldsymbol{\theta}_{D,est} = \underset{\boldsymbol{\theta}_D}{\operatorname{argmin}} \sum_{k=1}^{N_D} \left(f_D(t|\boldsymbol{\theta}_D) - I_{DD}(t_k^D)/I_{DD}(t_1^D)\right)^2.$$

Note that this is a rather crude estimation because the observed $\{I_{DD}(t_k^D)\}$ depends on the degree of FRET between the donor and acceptor, which we neglect here. However, since the effect of FRET on the intensity $I_{DD}$ is generally small (i.e., $f_D(t|\boldsymbol{\theta}_D)[D_{total}] > f_A(t|\boldsymbol{\theta}_A)f_D(t|\boldsymbol{\theta}_A)\chi(t)$), so this still gives an order-of-magnitude estimation. We can reflect the relatively high uncertainty by a relatively high $\sigma_0^2$, and set the prior as

$$p(\boldsymbol{\theta}_{D,i}) = \text{Lognormal}(\boldsymbol{\theta}_{D,i} | \log \boldsymbol{\theta}_{D,est,i} + \sigma_0^2, \sigma_0^2),$$

where $\boldsymbol{\theta}_{D,i}$ is the i-th element of $\boldsymbol{\theta}_D$.

The estimation of $\boldsymbol{\theta}_q$, which dictates the process noise of the hidden state $q \sim p(q|\boldsymbol{\theta}_q)$, is more challenging and requires a sophisticated inference algorithm like B-FRET. Still, a rough, order-of-magnitude estimation can be done for the purpose of constructing a prior distribution. First, we compute $\chi_k = E_{corr}(t_k^D)[D_{total}]_{est}$, where $E_{corr}(t_k^D)$, which gives an estimation of $\frac{E_{max}[DA_{total}]}{[D_{total}]}$ is obtained from the E-FRET formula (Online Methods) and the estimation of $[D_{total}]$, $[D_{total}]_{est}$, was obtained above. Although this is a highly noisy estimate of $\chi_k$ based on the information-inefficient E-FRET formula, this allows us to compute the distribution of $\Delta\chi_k = \chi_k - \chi_{k-1}$, which gives an estimation of $p(q|\boldsymbol{\theta}_{\boldsymbol{q}})$, enabling to obtain $\boldsymbol{\theta}_{q,est}$. Using, a large $\sigma_0^2$, we set the prior as

$$p(\boldsymbol{\theta}_{q,i}) = \text{Lognormal}(\boldsymbol{\theta}_{q,i} | \log \boldsymbol{\theta}_{q,est,i} + \sigma_0^2, \sigma_0^2),$$

where $\boldsymbol{\theta}_{D,i}$ is the i-th element of $\boldsymbol{\theta}_D$.

**References**


1. Kitagawa, G. *Introduction to time series modeling*. (Chapman and Hall/CRC, 2010).
2. Sarkka, S. *Bayesian Filtering and Smoothing*. (Cambridge University Press, 2013). doi:10.1017/CBO9781139344203.
3. MacKay, D. J. C. Information Theory, Inference, and Learning Algorithms. 640.
4. Bishop, C. M. *Pattern recognition and machine learning*. (Springer, 2006).

### Supplementary Note 3

#### Determining imaging-system parameters and measurement-noise levels

Keita Kamino, Nirag Kadakia, Kazuhiro Aoki, Thomas S. Shimizu, and Thierry Emonet

B-FRET assumes the knowledge of imaging-system dependent parameters, $a$, $b$, and $G$ (see Online Methods or below for definitions), which have been routinely measured in 3-cube FRET measurement setups[1–3]. Also assumed are the levels of measurement noise associate with the fluorescence signals $I_{AA}$, $I_{DD}$, and $I_{DA}$ as functions of frame number (or time). Here, we describe how we determined these.

**Measurements of imaging system parameters**

The measurements of imaging-systems parameters $a$, $b$, and $G$ were described elsewhere[1,2] in detail, and so here we describe them only briefly. First, the cross-talk coefficients $a$ and $b$ can be estimated by observing the fluorescent signals from strains that express only the acceptor or the donor because

$$a \equiv \frac{\nu_D \epsilon_{DA} t_{DA}}{\nu_A \epsilon_{AA} t_{AA}} \simeq \frac{I_{DA(A)}}{I_{AA(A)}}, \quad (\text{Eq. } 3-1)$$

$$d \equiv \frac{L_A S_A t_{DA}}{L_D S_D t_{DD}} \simeq \frac{I_{DA(D)}}{I_{DD(D)}}, \quad (\text{Eq. } 3-2)$$

where, $\nu_D$ ($\nu_A$) is the intensity of illumination reaching the sample through the donor (acceptor) excitation filter, $\epsilon_{DA}$ ($\epsilon_{AA}$) the absorption coefficient of the acceptor at the donor-excitation (acceptor-excitation) wavelength, $L_D$ ($L_A$) the throughput of the donor (acceptor) emission light-path, $S_D$ ($S_A$) the quantum sensitivity of the camera for donor (acceptor) emission, and $t_{DA}$, $t_{AA}$, and $t_{DD}$ respectively the exposure time for the FRET, acceptor, and donor channels; $A$ and $D$ in the parentheses in the lower index indicates that the corresponding fluorescent signals are obtained from the strain that only expresses the acceptor and the donor respectively. The approximations above become exact in the limit of zero measurement noise. This can be shown by noting that

$$I_{DA(A)} = [A^*]\nu_D \epsilon_{DA} Q_A L_A S_A t_{DA} + \xi_{DA(A)},$$

$$I_{AA(A)} = [A^*]\nu_A \epsilon_{AA} Q_A L_A S_A t_{AA} + \xi_{AA(A)},$$

$$I_{DA(D)} = [D^*]\nu_D \epsilon_{DD} Q_D L_A S_A t_{DA} + \xi_{DA(D)},$$

$$I_{DD(D)} = [D^*]\nu_D \epsilon_{DD} Q_D L_D S_D t_{DD} + \xi_{DD(D)},$$

where $\xi_{DA(A)}$, $\xi_{AA(A)}$, $\xi_{DA(D)}$ and $\xi_{DD(D)}$ represent measurement noise, $\epsilon_{DD}$ the absorption coefficient of the donor, and $Q_D$ ($Q_A$) the quantum yield of donor (acceptor). As has been done before[1–3], we obtained the estimates of $a$ by measuring $I_{DA(A)}$ and $I_{AA(A)}$ (Eq. 3-1) and $d$ by measuring $I_{DA(D)}$ and $I_{DD(D)}$ (Eq. 3-2) from many cells and linear least-squares fitting the data:

$$a = \underset{a'}{\operatorname{argmin}} \sum_i \left(a' I_{AA(A),i} - I_{DA(A),i}\right)^2,$$

$$d = \underset{d'}{\mathrm{argmin}} \sum_i \left(d' I_{DD(D),i} - I_{DA(D),i}\right)^2,$$

where subscripts $i$ indicates different cells. The values we obtained were $a = 0.3369\ (\pm 0.0006)$ and $d = 0.0891\ (\pm 0.0001)$ for the measurement system used for the *E. coli* chemotaxis pathway[2] (SE shown in parentheses), and $a = 0.346\ (0.331 - 0.357)$ and $d = 0.606\ (0.601 - 0.609)$ for the system used for the HeLa cAMP responses (Supplementary Fig. 3; 95% confidence intervals in the parentheses).

The parameter $G$ quantifies the change in sensitized emission define as $F_c = I_{DA} - aI_{AA} - dI_{DD}$ (Zal et al) per unit change in $I_{DD}$ due to FRET, namely $G \equiv \left|\frac{dF_c}{dI_{DD}}\right|$, which, using optical parameters, can be written as $G \equiv \frac{Q_A L_A S_A t_{DA}}{Q_D L_D S_D t_{DD}}$. This is in principle can be measured from a FRET strain expressing both donor and acceptor and by measuring fluorescence intensities before and after acceptor photobleaching:

$$G \simeq \frac{F_c'}{I_{DD}^{post} - I_{DD}'}, \quad (\mathrm{Eq.}\,3-3)$$

where $I_{DD}^{post}$ is the intensity of donor fluorescence after the acceptor is photobleached, and $F_c'$ and $I_{DD}'$ correspond to $F_c$ and $I_{DD}$, respectively, in the absence of photobleaching[1]. Again, the approximation is exact in the limit of zero measurement noise. The relation can be shown by noting

$$F_C' = [D^*A^*]v_D\epsilon_{DD}E_{max}Q_A L_A S_A t_{DA} + \xi_{F_c},$$

$$I_{DD}^{post} - I_{DD}' = [D^*A^*]v_D\epsilon_{DD}E_{max}Q_D L_D S_D t_{DD} + \xi_{DD}',$$

where $\xi_{F_c}$ and $\xi_{DD}'$ represent measurement noise. However, acceptor photobleaching can cause confounding effects such as photoconversion[4], and thus Eq. 3-3 is not very useful as an equation to empirically estimate $G$. Alternatively, if one can induce changes in FRET by, e.g., external stimuli, $G$ can be measured without acceptor photobleaching[2]. To see this, consider two states ($j = 1$ or 2) of a FRET sample that have different FRET levels. For each state, the equation 3-3 holds:

$$G \simeq \frac{F_{c,j}'}{I_{DD}^{post} - I_{DD,j}'},$$

where $F_{c,j}'$ and $I_{DD,j}'$ are $F_c'$ and $I_{DD}'$ when the sample state is $j$ ($= 1$ or 2). Note $G$ and $I_{DD}^{post}$ are common in the two states. By deleting $I_{DD}^{post}$ from the two equations $G$ can be expressed as

$$G \simeq \frac{\left|F_{c,2}' - F_{c,1}'\right|}{\left|I_{DD,2}' - I_{DD,1}'\right|}.$$

Based on this expression, we estimated the value of $G$ by least-squares fitting the fluorescence signals from multiple cells, i.e.,

$$G = \underset{G'}{\mathrm{argmin}} \sum_i \left(G'\left|I_{DD,2}' - I_{DD,1}'\right| - \left|F_{c,2}' - F_{c,1}'\right|\right)^2,$$

where subscripts $i$ indicates different cells. The value obtained was $G = 0.3497\ (\pm 0.0018)$ for the system used for the *E. coli* chemotaxis pathway[2] (SE shown in parentheses), and $G = 1.316\ (1.253 - 1.396)$ for the system used for the HeLa cAMP responses (Supplementary Fig. 3; 95% confidence intervals in the parentheses).

#### Effects of parameter-estimation error on the FRET signal

The imaging-system parameters $a, d$ and $G$ determined above can only be measured with finite precision; therefore the determined values inevitably contain some error. The FRET index we estimate $E = E_{max}[DA_{total}]/[D_{total}]$, when expressed by using observables, is a function of these parameters, as can be seen in the E-FRET formula[1] (Online Methods):

$$E_{corr}(t) = \frac{I_{DA}(t) - a\overline{I_{AA}(t)} - dI_{DD}(t)}{I_{DA}(t) - a\overline{I_{AA}(t)} + (G-d)I_{DD}(t)} \frac{\overline{I_{AA}(t=0)}}{\overline{I_{AA}(t)}},$$

therefore the errors in $a, d$ and $G$ necessarily bias the estimation of $E$. How these errors affect the estimation of $E$ was quantitatively investigated previously[2], and we reproduce the discussion below to make our argument self-contained. Briefly, the conclusions are the following. (i) The bias in the *absolute level* of estimated $E$ grows exponentially as more acceptors and donors are photobleached; this generates an increasing or decreasing trend – the sign depends on the sign of the errors – over time in the estimated $E$, even if the actual degree of molecular interactions remains unchanged over time. However, (ii) the bias in the *changes* in the estimated $E$ that occurs at a time scale faster than the (generally slow) photobleaching is small. This means that changes in the estimated $E$ around the slowly increasing or decreasing trend are reliable signals. Because of these properties, for both E-FRET and B-FRET results, we subtracted slowly increasing or decreasing trends by fitting a linear or exponential function[2].

Here, we show the effects of the error in the imaging-system-parameter estimations[2]. The estimated values of the parameters can be written as

$$a_{est} = a + \Delta a,$$

$$d_{est} = d + \Delta d,$$

$$G_{est} = G + \Delta G,$$

where true values of the parameters are denoted by $a$, $d$, and $G$ and the deviations from them by $\Delta a$, $\Delta d$, and $\Delta G$. First, using $F_c = I_{DA}(t) - a\overline{I_{AA}(t)} - dI_{DD}(t)$, we note that $E_{corr}$ can be approximated as

$$E_{corr} = \frac{\frac{F_c}{I_{DD}}}{\frac{F_c}{I_{DD}} + G} \frac{I_{AA}(0)}{I_{AA}}$$

$$\approx \frac{I_{AA}(0)}{G} \frac{F_c}{I_{DD} I_{AA}}$$

$$= \frac{I_{AA}(0)}{G} \frac{I_{DA} - d\, I_{DD} - a\, I_{AA}}{I_{DD} I_{AA}}.$$

In the second line, we used $\frac{F_c}{I_{DD}} \ll G$ to simplify the following calculation, although it is not essential. This assumption is valid in a typical low FRET-efficiency experiment where the value of $F_c$ is sufficiently lower than $I_{DD}$ $\left(\frac{F_c}{I_{DD}} \ll 1\right)$ and yet, to be able to detect FRET signals, the parameter $G$ needs to be $\sim\mathcal{O}(1)$. For example, in our setup for bi-molecular FRET, $\frac{F_c}{I_{DD}} \lesssim 0.05$ and $G \simeq 0.35$ (ref.[1])

The error in $E_{corr}$ due to the error in the estimated parameters $\Delta a$, $\Delta d$, and $\Delta G$ can be written as

$$\Delta E_{corr} = E_{corr}(a+\Delta a, d+\Delta \mathrm{d}, G+\Delta G) - E_{corr}(a,d,G)$$

$$\simeq \frac{\partial E_{corr}(a,d,G)}{\partial a}\Delta a + \frac{\partial E_{corr}(a,d,G)}{\partial d}\Delta d + \frac{\partial E_{corr}(a,d,G)}{\partial G}\Delta G$$

$$\simeq -\frac{I_{AA}(0)}{GI_{DD}}\Delta a - \frac{I_{AA}(0)}{GI_{AA}}\Delta d - \frac{E_{corr}}{G}\Delta G$$

Thus, the fraction of error in $E_{corr}$ can be written as

$$\frac{\Delta E_{corr}}{E_{corr}} = -\frac{I_{AA}\Delta a}{F_c} - \frac{I_{DD}\Delta d}{F_c} - \frac{\Delta G}{G}.$$

As derived in Supplementary Note 1, the observables $I_{DD}$ and $I_{AA}$ and the sensitized emission $F_c$ can be written as

$$I_{DD} \simeq C_{DD}\left([X_{total}]\, e^{-\int_0^t \delta(t')dt'} - E_{max}\,[XY]\, e^{-\int_0^t \alpha(t')+\delta(t')dt'}\right) \sim C_{DD}\,[X_{total}]\, e^{-\int_0^t \delta(t')dt'}$$

$$I_{AA} \simeq C_{AA}\,[Y_{total}]\, e^{-\int_0^t \alpha(t')dt'}$$

$$F_c(t) \simeq C_{DD}\, G\, E_{max}\,[XY]\, e^{-\int_0^t \alpha(t')+\delta(t')dt'},$$

where $\delta(t) > 0$ and $\alpha(t) > 0$ are, respectively, the (time-dependent) rates of photobleaching of the donor and acceptor, and the final approximation for $I_{DD}$ is valid under the assumption $\frac{F_c}{I_{DD}} \ll G_E$.

Using these expressions, we get

$$\frac{\Delta E_{corr}}{E_{corr}} \sim A\, e^{\int_0^t \delta(t')dt'}\Delta a + D\, e^{\int_0^t \alpha(t')dt'}\Delta d - \frac{\Delta G}{G},$$

where $A = \frac{C_{AA}\,[Y_{total}]}{C_{DD}\,G\,E_{max}\,[XY]} > 0$ and $D = \frac{[X_{total}]}{GE_{max}\,[XY]} > 0$. The first and the second terms grow quasi-exponentially as the fluorescent proteins photobleach; thus, the measured value of $E_{corr}$, at baseline levels of molecular interaction, changes over time. Note that the time scale of this change is governed by the time scale of photobleaching.

The remaining question is how uncertainty in the parameters $a$, $d$, and $G$, in the presence of photobleaching, affects the mapping between *changes* in molecular interactions and the corresponding *change* in $E_{corr}$. To address this, we analyze the sensitivity of $E_{corr}$ to the change in the degree of molecular interaction and its dependence on photobleaching.

The degree of molecular interaction is dictated by the time-dependent binding affinity $\gamma(t)$ between the two target molecules X and Y. Therefore, the sensitivity of $E_{corr}$ to changes in $\gamma$ at a given time can be quantified by $\frac{\partial E_{corr}(\gamma|a,d,G)}{\partial \gamma}$. With errors in the parameters, this quantity can be written as

$$\frac{\partial E_{corr}(\gamma|a+\Delta a, d+\Delta d, G+\Delta G)}{\partial \gamma} = \frac{\partial E_{corr}(\gamma|a,d,G)}{\partial \gamma} + \frac{\partial \Delta E_{corr}(\gamma)}{\partial \gamma}$$

$$= \frac{\partial E_{corr}(\gamma|a,d,G)}{\partial \gamma}\left(1 + \frac{\frac{\partial \Delta E_{corr}(\gamma)}{\partial \gamma}}{\frac{\partial E_{corr}(\gamma|a,d,G)}{\partial \gamma}}\right) \equiv \frac{\partial E_{corr}(\gamma|a,d,G)}{\partial \gamma}(1+\Delta).$$

Thus, $\Delta \equiv \frac{\frac{\partial \Delta E_{corr}}{\partial \gamma}}{\frac{\partial E_{corr}(\gamma|a,d,G)}{\partial \gamma}}$ characterizes the bias error, and the question is how this quantity behaves with photobleaching. To compute this, we note

$$\Delta = \frac{\frac{\partial}{\partial \gamma}\left(-\frac{I_{AA}(0)}{GI_{DD}(\gamma)}\Delta a - \frac{I_{AA}(0)}{GI_{AA}}\Delta d - \frac{E_{corr}(\gamma)}{G}\Delta G\right)}{\frac{\partial E_{corr}(\gamma)}{\partial \gamma}}$$

$$= \frac{-\frac{I_{AA}(0)\Delta a}{G}\frac{\partial}{\partial \gamma}\left(\frac{1}{I_{DD}(\gamma)}\right) - \frac{\Delta G}{G}\frac{\partial E_{corr}}{\partial \gamma}}{\frac{\partial E_{corr}}{\partial \gamma}}$$

$$= \frac{-\frac{I_{AA}(0)\Delta a}{G}\frac{\partial}{\partial \gamma}\left(\frac{1}{I_{DD}(\gamma)}\right)}{\frac{\partial E_{corr}}{\partial \gamma}} - \frac{\Delta G}{G},$$

where we used the fact that $I_{AA}$ is independent of $\gamma$, i.e., $\frac{\partial I_{AA}}{\partial \gamma} = 0$. We note

$$\frac{\partial E_{corr}}{\partial \gamma} \simeq \frac{\partial}{\partial \gamma}\left(\frac{1}{G}\frac{F_c(\gamma)}{I_{DD}(\gamma)}\frac{I_{AA}(0)}{I_{AA}}\right)$$

$$= \frac{I_{AA}(0)}{GI_{AA}}\frac{\partial}{\partial \gamma}\left(\frac{F_c}{I_{DD}}\right)$$

$$= \frac{I_{AA}(0)}{GI_{AA}}\left(\frac{1}{I_{DD}}\frac{\partial F_c}{\partial \gamma} - \frac{F_c}{I_{DD}^2}\frac{\partial I_{DD}}{\partial \gamma}\right)$$

$$= -\frac{I_{AA}(0)}{GI_{AA}}\frac{1}{I_{DD}}\frac{\partial I_{DD}}{\partial \gamma}\left(-\frac{\frac{\partial F_c}{\partial \gamma}}{\frac{\partial I_{DD}}{\partial \gamma}} + \frac{F_c}{I_{DD}}\right)$$

$$= -\frac{I_{AA}(0)}{GI_{AA}}\frac{1}{I_{DD}}\frac{\partial I_{DD}}{\partial \gamma}\left(G + \frac{F_c}{I_{DD}}\right),$$

where at the final step we used $-\frac{\partial F_c}{\partial \gamma}/\frac{\partial I_{DD}}{\partial \gamma} = |\Delta F_c|/|\Delta I_{DD}| = G$. By plugging this to the expression for $\Delta$, we get

$$\Delta = \frac{\frac{I_{AA}(0)\Delta a}{G}\frac{1}{I_{DD}^2}\frac{\partial I_{DD}}{\partial \gamma}}{-\frac{I_{AA}(0)}{GI_{AA}}\frac{1}{I_{DD}}\frac{\partial I_{DD}}{\partial \gamma}\left(G + \frac{F_c}{I_{DD}}\right)} - \frac{\Delta G}{G}$$

$$= \frac{-I_{AA}\Delta a_E}{I_{DD}\left(G + \frac{F_c}{I_{DD}}\right)} - \frac{\Delta G}{G}$$

$$\simeq -\frac{I_{AA}\,\Delta a_E}{I_{DD}\,G} - \frac{\Delta G}{G}$$

$$= H\, e^{\int_0^t \delta(t') - \alpha(t') dt'}\Delta a - \frac{\Delta G}{G},$$

where $H = \frac{C_{AA}\,[Y_{total}]}{C_{DD}\,[X_{total}]G} > 0$ and in the third line we used $\frac{F_c}{I_{DD}} \ll G$. This expression tells us that the relative error in the mapping from molecular interaction to $E_{corr}$, $\Delta$, is small if $\Delta a$ and $\Delta G$ are small. Furthermore, this relative error grows slower than the relative error in the baseline of $E_{corr}$, $\frac{\Delta E_{corr}}{E_{corr}}$, because only the difference between the donor and acceptor photobleaching rates appears in the exponential. Additionally, the coefficient $H$ is typically smaller than the coefficients in $\frac{\Delta E_{corr}}{E_{corr}}$, $A$ and $D$. In fact, assuming $C_{AA} \approx C_{DD}$ and $[Y_{total}] \approx [X_{total}]$, one can show that both $H/A$ and $H/D$ are bounded by $\frac{E_{max}\,[XY]}{[X_{total}]} < 1$.

**Measurement noise estimation**

We assume Gaussian measurement noise $\xi_{AA}(t)$, $\xi_{DD}(t)$, and $\xi_{DA}(t)$ for the fluorescence signals $I_{AA}(t)$, $I_{DD}(t)$, and $I_{DA}(t)$, respectively (Online Methods). Thus, the measurement noise can be written as

$$\xi_{AA}(t) \sim \mathrm{N}\left(0, \sigma_{AA}^2(t)\right),$$

$$\xi_{DD}(t) \sim \mathrm{N}\left(0, \sigma_{DD}^2(t)\right),$$

$$\xi_{DA}(t) \sim \mathrm{N}\left(0, \sigma_{DA}^2(t)\right).$$

We would like to estimate the time-dependent noise variances $\sigma_{AA}^2(t)$, $\sigma_{DD}^2(t)$, and $\sigma_{DA}^2(t)$ from data. The Gaussian approximation is sufficiently precise for typical FRET measurements where the shot noise (or Poisson noise) originating from photon counting is the dominant source of measurement noise. When necessary, however, it is straightforward to incorporate into the B-FRET framework measurement noise that follows different probability distributions: one only needs to change the likelihood functions of model parameters accordingly (Online Methods and Supplementary Note 2).

The basic idea of the noise-variance estimation we used here draws on the fact that measurement noise is delta-correlated and hence fast, whereas other sources of changes in fluorescence signals, such as photobleaching and changes in donor-acceptor interactions, are slower. If the timescale separation is clear, the noise-variance estimation is easy since one just needs to subtract the slowly-changing components (estimated by, e.g., moving average of a time series) before computing variances of a time series. An example of such simple method can be found in Supplementary Note 5 where we applied the method to FRET data from cAMP responses of HeLa cells. However, in general, such "slow" factors may also contain some frequency components higher than the sampling frequency, which appears to be delta-correlated and requires a little more sophisticated method. Also, the fact that the noise variance changes over time due to, e.g., photobleaching complicates its estimation. Below, we describe a principled, general method to estimate the noise variances applicable to most fluorescence time-series data.

First, note that, if the noise variance is constant in time and a time-series data is sufficiently long, the estimation of noise variance is straightforward. This is achieved by using the autocorrelation function $C(\tau) = \langle I(t+\tau)I(t)\rangle$. In this function, the power of white noise is concentrated at $\tau = 0$, and therefore $C(0)$ estimates the sum of the noise variance and the power of the delta-correlated component form the other sources. On the other hand, $C(\tau)$ for $\tau > 0$ only estimates the variance from other sources and does not contain the power of measurement noise. Thus, defining $C'(0)$ as the extrapolated value at $\tau = 0$ from $C(\tau)$ for $\tau > 0$, the difference between $C(0)$ and $C'(0)$ gives the noise variance.

The method based on the auto-correlation function assumes constant noise variance. However, in fluorescence time-series data, the noise variance changes over time. Therefore, the method is applicable only to short local segments of a time series, where one can safely assume that the noise variance is approximately constant. As a result of segmenting a time series into short snippets, however, the noise-variance estimation suffers from higher statistical uncertainty, making it more challenging to estimate the trend of noise variance precisely. To address this, we draw on the fact (shown below) that, in the regime where shot noise is dominant, the variance of measurement noise ($\mathrm{Var}(I(t))$) is proportional to the expected value of a fluorescence intensity $\langle I(t)\rangle$, namely:

$$\alpha = \frac{\mathrm{Var}(I(t))}{\langle I(t)\rangle}. \quad (\mathrm{Eq.}\,3-3)$$

Here, $\alpha$ is an unknown proportionality constant that is fixed for a given fluorescence channel irrespective of the magnitude of a fluorescence intensity (shown below). Therefore, once one obtains $\alpha$ for a given fluorescence channel, the problem of estimating noise variance at each time point (i.e., $\mathrm{Var}(I(t))$) is reduced to the problem of estimating the expected value of a fluorescence intensity at each time point (i.e., $\langle I(t)\rangle$), which is much easier. We estimated $\langle I(t)\rangle$ by fitting single or bi-exponential functions to a fluorescence time-series data.

To estimate $\alpha$ for a fluorescence channel, we plotted the estimations of $\mathrm{Var}(I(t))$ against the estimations $\langle I(t)\rangle$ and determined the slope (Supplementary Fig. 5). Although both estimations suffer from statistical uncertainty, we reduced the uncertainty by aggregating many data. To be concrete, by denoting estimations of average and variance for a snippet time series $\{I_i(t)\}$ (labeled by $i$) by $A(\{I_i(t)\})$ and $V(\{I_i(t)\})$ respectively, we determine $\alpha$ by computing

$$\alpha = \underset{\alpha'}{\mathrm{argmin}} \sum_i \left( \alpha' A(\{I_i(t)\}) - V(\{I_i(t)\})\right)^2.$$

Now we show the proportionality between the expectation of a fluorescence intensity and its variance (Eq. 3-3). We start from the assumption that the number of photons $n_p$ from a sample and collected by a microscopy fluorescence channel follows Poisson distribution

$$p(n_p|\Lambda) = \frac{\Lambda^{n_p} e^{-\Lambda}}{n_p!},$$

where the average and variance of $n_p$ are the same, $\Lambda = \langle n_p\rangle = \mathrm{Var}(n_p)$. At the detector, the photons are converted into photoelectrons with a wave-length-dependent efficiency $QE(\lambda)$ on average, and thus the average number of photoelectrons $n_{e^-}$ can be written as

$$\langle n_{e^-}\rangle = QE(\lambda) \times \langle n_p\rangle.$$

The variance of $n_{e^-}$ can be written as

$$\mathrm{Var}(n_{e^-}) = F_n^2 \times QE(\lambda)^2 \times \mathrm{Var}(n_p),$$

where $F_n$ is called noise factor associated with the multiplicative noise in signal amplifying process, and typically $F_n = \sqrt{2}$ for an EM-CCD detector and $F_n = 1$ for CCD and CMOS detectors. The photoelectrons are converted into pixel counts (or intensity), and this can be written as

$$I = \frac{n_{e^-}}{CF},$$

where $CF$ is a conversion factor (electron/count) dependent on a detector. On average, the intensity and the number of photons are connected by

$$\langle I \rangle = \frac{QE(\lambda)}{CF} \langle n_p \rangle.$$

The variance of the intensity can be written as

$$\mathrm{Var}(I) = \mathrm{Var}\left(\frac{n_{e^-}}{CF}\right) = \frac{1}{CF^2} \mathrm{Var}(n_{e^-})$$

$$= \frac{QE(\lambda)^2 F_n^2}{CF^2} \mathrm{Var}(n_p)$$

$$= \frac{QE(\lambda)^2 F_n^2}{CF^2} \langle n_p \rangle$$

$$= \frac{QE(\lambda)}{CF} F_n^2 \langle I \rangle.$$

Therefore, we get

$$\frac{\mathrm{Var}(I)}{\langle I \rangle} = \frac{QE(\lambda)}{CF} F_n^2 \equiv \alpha.$$

Thus, $\alpha$ defined in Eq. 3-3 is constant for a fluorescent channel and can be estimated by using data with different absolute values of $\mathrm{Var}(I)$ and $\langle I \rangle$.

**References**


1. Zal, T. & Gascoigne, N. R. J. Photobleaching-Corrected FRET Efficiency Imaging of Live Cells. *Biophys. J.* **86**, 3923–3939 (2004).
2. Mattingly, H. H., Kamino, K., Machta, B. B. & Emonet, T. Escherichia coli chemotaxis is information limited. *Nat. Phys.* **17**, 1426–1431 (2021).
3. Babel, H. *et al.* Ratiometric population sensing by a pump-probe signaling system in Bacillus subtilis. *Nat. Commun.* **11**, 1176 (2020).
4. Miyawaki, A. Development of Probes for Cellular Functions Using Fluorescent Proteins and Fluorescence Resonance Energy Transfer. *Annu. Rev. Biochem.* **80**, 357–373 (2011).

### Supplementary Note 4

#### Synthetic data and model functions used to analyze data

Keita Kamino, Nirag Kadakia, Kazuhiro Aoki, Thomas S. Shimizu, and Thierry Emonet

**Synthetic data**

All the synthetic data in this paper were generated according to the following equations (see Eqs. 3 in Online Methods):

$$I_{AA}(t) = C_{AA} f_A(t)[A_{total}] + \xi_{AA}(t),$$

$$I_{DD}(t) = C_{DD}\big(f_D(t)[D_{total}] \; - f_A(t) f_D(t)\chi(t)\big) + \xi_{DD}(t),$$

$$I_{DA}(t) = aC_{AA} f_A(t)[A_{total}] + dC_{DD} f_D(t)[D_{total}] + C_{DD}(G-d) f_A(t) f_D(t)\chi(t) + \xi_{DA}(t).$$

See **Photophysical model** in Online Methods for how these equations are derived and for the definitions of parameters and variables. $\xi_{AA}$, $\xi_{DD}$ and $\xi_{DA}$ are the stochastic variables that represent the measurement noise whose variances are dependent on time. We emulated a situation where shot-noise is the dominant source of measurement noise. Thus, the variance of shot noise is proportional to the expected value of a fluorescence intensity (Supplementary Note 3), and therefore we write

$$\xi_{AA}(t) \sim \mathrm{N}(0, \alpha_A C_{AA} f_A(t)[A_{total}]),$$

$$\xi_{DD}(t) \sim \mathrm{N}\Big(0, \alpha_D \Big(C_{DD} f_D(t)\big([D_{total}] \; - f_A(t)\chi(t)\big)\Big)\Big),$$

$$\xi_{DD}(t_k) \sim \mathrm{N}\Big(0, \alpha_D\big(aC_{AA} f_A(t)[A_{total}] + dC_{DD} f_D(t)[D_{total}] + C_{DD}(G-d) f_A(t) f_D(t)\chi(t)\big)\Big),$$

where $\alpha_A$ and $\alpha_D$ are the proportionality constants that converts the expected value of an intensity into the variance (Supplementary Note 3).

Common parameter values used in all cases below are: $[A_{total}] = 18, [D_{total}] = 8, C_{DD} = 1360, C_{AA} = 340, a = 0.35, d = \; 0.09, G = 0.5, E_{max} = 0.1, \alpha_A = 1/0.46, \; \alpha_D = 1/0.46.$

<u>Oscillatory FRET data (Fig. 1)</u>

We assumed photobleaching dynamics that follows

$$f_A(t) = \delta_A e^{-\frac{t}{\tau_{A1}}} + (1-\delta_A) e^{-\frac{t}{\tau_{A2}}},$$

$$f_D(t) = \delta_D e^{-\frac{t}{\tau_{D1}}} + (1-\delta_D) e^{-\frac{t}{\tau_{D2}}},$$

where $\tau_{D1} = 1200, \tau_{D2} = 120, \tau_{A1} = 3000, \tau_{A2} = 300, \delta_A = 0.6, \delta_D = 0.6$. We also assumed all three fluorescence signals $I_{AA}$, $I_{DD}$, and $I_{DA}$ were sampled at $\{t_k\} = \{0, 0.5, 1, \ldots, 600\}$ with fixed interval $\Delta t = 0.5$. The hidden variable $\chi(t)$ follows

$$\chi(t) = a_0 + a_1 \sin(\omega t_k) + a_2 \sin(b\omega t_k),$$

where $a_0 = 0.5, a_1 = 0.12, a_2 = 0.1, b = 2.5, \omega = 0.1$.

Random FRET data (Fig. 2)

We assumed the same photobleaching dynamics and sampling time as the oscillatory FRET data. The dynamics of $\chi(t)$ was assumed to follow the Ornstein-Uhlenbeck process,

$$\frac{d\chi}{dt} = -\frac{1}{\tau_c}(\chi(t) - \chi_0) + \sqrt{2D_n}\xi(t),$$

where $\xi(t)$ is a Gaussian white noise with average zero and a delta correlation in time:

$$\langle \xi(t) \rangle = 0, \quad \langle \xi(t)\xi(t') \rangle = \delta(t - t'),$$

where $\delta(t)$ is the Dirac delta function. The parameters used were $\chi_0 = 0.5$, $\tau_c = 5$, $D_n = 0.025/\tau_c$. We simulated this process by using the following update rule for discretized time $\Delta t = 0.5$, which exactly reproduces the continuous dynamics[1]:

$$\chi(t + \Delta t) \sim \mathrm{N}\left(\chi(t)e^{-\Delta t/\tau_{OU}} + \chi_0, D_n\tau_{OU}\left(1 - e^{-\frac{2\Delta t}{\tau_{OU}}}\right)\right),$$

where $\mathrm{N}(\mu, \sigma^2)$ is a Gaussian distribution. After computing all $\{\chi(t)\}$, we replaced negative values of $\chi(t)$ with zero to satisfy $\chi(t) \geq 0$.

Step FRET data (Fig. 2)

We assumed the same photobleaching dynamics and sampling time as the oscillatory FRET data. The dynamics of $\chi(t)$ follows

$$\chi(t) = \begin{cases} 0.3 \ (300 < t \leq 450, 900 < t \leq 1150) \\ 0.5 \ (0 \leq t \leq 150, 450 < t \leq 750, 1150 < t) \\ 0.7 \ (150 < t \leq 300, 750 < t \leq 900) \end{cases}$$

Random FRET data in various measurement conditions (Fig. 3)

We assumed single-exponential photobleaching dynamics:

$$f_A(t) = e^{-\frac{t}{\tau_{A1}}},$$

$$f_D(t) = e^{-\frac{t}{\tau_{D1}}},$$

where $\tau_{D1} = 4800$ $\tau_{A1} = 12000$. We assumed all three fluorescence signals $I_{AA}$, $I_{DD}$, and $I_{DA}$ were sampled at $\{t_k\} = \{0, 0.5, 1, \ldots, 1200\}$ with fixed interval $\Delta t = 0.5$. The dynamics of $\chi(t)$ was assumed to follow the Ornstein-Uhlenbeck process

$$\frac{d\chi}{dt} = -\frac{1}{\tau_c}(\chi(t) - \chi_0) + \sqrt{2D_n}\xi(t),$$

essentially in the same way as above, but we explored different time constants ranging from $\tau_c = 0.025$ (under-sampling) to $\tau_c = 100$ (over-sampling) while keeping the long-term variance fixed to $D_n\tau_c = 0.025$. We also simulated different measurement-noise levels by changing the proportionality constants that converts the expected value of an intensity into the variance $\alpha_A$ and $\alpha_D$. We explored the values ranging from $\alpha_A = \alpha_D = 0.02$ (high SNR) to $\alpha_A = \alpha_D = 40$ (low SNR). When different $\tau_c$ was explored, representative values of $\alpha_A = \alpha_D = 0.2$ and $\alpha_A = \alpha_D = 2$ were used for high and low SNR conditions respectively. When different levels of measurement noise were explored, representative values of $\frac{\tau_c}{\Delta t} \approx 1$ and $\frac{\tau_c}{\Delta t} \approx 10$ were used for under- and over sampling regimes respectively.

**Settings for B-FRET analyses**

Here we summarize the exact models and priors used to analyze each of the data set presented in this paper. See **Photophysical model** in Online Methods or **Overview** in Supplementary Note 2 for the definition of the model used in all the analyses.

Analysis of the synthetic oscillatory FRET data (Fig. 1)

The photobleaching functions we used were bi-exponential functions

$$f_A(t; \delta_A, \tau_{A1}, \tau_{A2}) = \delta_A e^{-\frac{t}{\tau_{A1}}} + (1 - \delta_A) e^{-\frac{t}{\tau_{A2}}},$$

$$f_D(t; \delta_D, \tau_{D1}, \tau_{D2}) = \delta_D e^{-\frac{t}{\tau_{D1}}} + (1 - \delta_D) e^{-\frac{t}{\tau_{D2}}}.$$

The prior distributions for all model parameters were constructed as described in **Prior distributions of the model parameters** in Supplementary Note 2. Specifically, uniform distributions bounded by $[0, 1]$ were used for the priors for $\delta_A$ and $\delta_D$, and log-normal distributions, $\mathrm{lognormal}(x|\mu, \sigma^2)$, were used for the other parameters. The modes of the log-normal prior distributions ($\exp(\mu - \sigma^2)$) were determined as described in **Prior distributions of the model parameters**. The parameter $\sigma$ was set to $\log 2$ for $\tau_{A1}$, $\tau_{A2}$, $\tau_{D1}$, $\tau_{D2}$, $[D_{total}]$, and $[A_{total}]$. For a Gaussian process noise, whose variance is parameterized by $\sigma_\chi^2$, we used the prior distribution

$$\sigma_\chi \sim \mathrm{lognormal}\left(\sigma_\chi | \log 10 + (\log 30)^2, (\log 30)^2\right).$$

For a Non-Gaussian process noise, we used the Student's t-distribution $St(q|\sigma_\chi, \nu)$ (Online Methods), and used the priors of

$$\sigma_\chi \sim \mathrm{lognormal}\left(\sigma_\chi | \log 10 + (\log 30)^2, (\log 30)^2\right),$$

and

$$\nu \sim \mathrm{lognormal}(\nu | \log 1 + (\log 10)^2, (\log 10)^2).$$

Analysis of the synthetic random FRET data (Fig. 2)

The same model and priors as the synthetic oscillatory FRET data were used.

Analysis of the synthetic step FRET data (Fig. 2)

The same model and priors as the synthetic oscillatory FRET data were used.

Analysis of the synthetic random FRET data in various measurement conditions (Fig. 3)

The photobleaching functions we used were bi-exponential functions

$$f_A(t;\tau_{A1}) = e^{-\frac{t}{\tau_{A1}}},$$

$$f_D(t;\tau_{D1},) = e^{-\frac{t}{\tau_{D1}}}.$$

Again, the prior distributions for all model parameters were constructed as described in **Prior distributions of the model parameters** in Supplementary Note 2. Log-normal distributions, $\mathrm{lognormal}(x|\mu,\sigma^2)$, were used for all parameters. The parameter $\sigma$ was set to $\log 2$ for $\tau_{A1}$, $\tau_{D1}$, $[D_{total}]$, and $[A_{total}]$. For a Gaussian process noise, whose variance is parameterized by $\sigma_\chi^2$, we used the prior distribution

$$\sigma_\chi \sim \mathrm{lognormal}\big(\sigma_\chi | \log 10 + (\log 30)^2, (\log 30)^2\big).$$

Analysis of the FRET data from single E. coli cells (Fig. 4)

Because the rate of photobleaching was relatively small for the acceptor while it is relatively large for the donor, we used single-exponential and bi-exponential functions for the acceptor and donor respectively:

$$f_A(t;\tau_{A1}) = e^{-\frac{t}{\tau_{A1}}},$$

$$f_D(t;\delta_D,\tau_{D1},\tau_{D2}) = \delta_D e^{-\frac{t}{\tau_{D1}}} + (1-\delta_D)e^{-\frac{t}{\tau_{D2}}}.$$

We used the Student's t-distribution $St(q|\sigma_\chi,\nu)$ (Online Methods) for the process noise.

Again, the prior distributions were determined **Prior distributions of the model parameters** in Supplementary Note 2. Briefly, uniform distribution bounded by $[0,1]$ was used for the prior of $\delta_D$. Log-normal distributions, $\mathrm{lognormal}(x|\mu,\sigma^2)$, were used for the other parameters. The parameter $\sigma$ was set to $\log 2$ for $\tau_{A1}$, $\tau_{D1}$, $\tau_{D2}$, $[D_{total}]$, and $[A_{total}]$. The priors for the process noise parameters were:

$$\sigma_\chi \sim \mathrm{lognormal}\big(\sigma_\chi | \log 10^3 + (\log 30)^2, (\log 30)^2\big),$$

and

$$\nu \sim \mathrm{lognormal}(\nu | \log 1 + (\log 10)^2, (\log 10)^2).$$

Analysis of the FRET data from single eukaryotic cells (Fig. S3)

We used the same photobleaching functions as those used for *E. coli* data, and a Gaussian distribution for the process noise. About the prior distributions, first, uniform distribution bounded by $[0,1]$ was used for the prior of $\delta_D$. Log-normal distributions, $\mathrm{lognormal}(x|\mu,\sigma^2)$, were used for the other parameters. The parameter $\sigma$ was set to $\log 10$ for $\tau_{A1}$, $\tau_{D1}$, $\tau_{D2}$, $[D_{total}]$, and $[A_{total}]$. For the standard deviation of a Gaussian process noise, we used the prior distribution

$$\sigma_\chi \sim \mathrm{lognormal}\left(\sigma_\chi \,|\, \log 1 + (\log 30)^2, (\log 30)^2\right).$$

**References**


1. Gillespie, D. T. Exact numerical simulation of the Ornstein-Uhlenbeck process and its integral. *Phys. Rev. E* **54**, 2084–2091 (1996).

### Supplementary Note 5

#### Unimolecular FRET experiment

Keita Kamino, Nirag Kadakia, Kazuhiro Aoki, Thomas S. Shimizu, and Thierry Emonet

Although we focused on a bimolecular FRET system expressed in *E. coli* cells in the main text, the B-FRET method is equally applicable to unimolecular FRET systems, where, unlike bimolecular FRET systems, the donor and acceptor are fused to the same molecule. We demonstrated this by applying B-FRET to a specific unimolecular FRET system expressed in HeLa cells (Supplementary Fig. 3). Below we describe the details of the experiment and the analysis of the obtained data.

**Strains and plasmids**

The cAMP FRET biosensor (mTFP-Epac-mVenus) was developed based on the previous work[1]. This contains the human RAPGEF3 (EPAC) gene (corresponding to amino acids 149-881). The cDNA of the cAMP biosensor was inserted into the pCX4neo vector[2], providing pCX4neo-mTFP-Epac-mVenus. This vector was used for transient expression. The cDNAs for mVenus and mTFP were subcloned into pCAGGS vector[3] generating pCAGGS-mVenus and pCAGGS-mTFP, respectively.

HeLa cells, a kind gift from Dr. Matsuda (Kyoto University, Japan), were cultured in Dulbecco's Modified Eagle's Medium (DMEM) high glucose (Wako; nacalai tesque) supplemented with 10% fetal bovine serum (Sigma-Aldrich) at 37°C in 5% CO2.

**Measurements of optical parameters**

HeLa cells were plated on CELLview cell culture dishes (glass bottom, 35 mm diameter, 4 components: The Greiner Bio-One) one day before transfection. The cells were transfected with the plasmids pCAGGS-mVenus or pCAGGS-mTFP by 293fectin transfection reagent (Thermo Fisher Scientific). One day after the transfection, the medium was replaced with the imaging medium (FluoroBrite (nacalai tesque)/1x GlutaMAX (GIBCO)/0.1% BSA), followed by fluorescence imaging.

**Cell preparation and FRET measurements**

HeLa cells were plated on CELLview cell culture dishes one day before transfection. The cells were transfected with plasmids encoding cAMP FRET biosensor (pCX4neo-mTFP-Epac-mVenus) by 293fectin transfection reagent. One day after the transfection, the medium was replaced with the imaging medium (FluoroBrite (nacalai tesque)/1x GlutaMAX (GIBCO)/0.1% BSA) several hours before the imaging was started. 50 uM Forskolin and 100 uM IBMX, both of which were purchased from Wako (Osaka, Japan), were applied to the cells 10 min after the start of time-lapse imaging.

**FRET imaging system**

Images were acquired on an IX81 inverted microscope (Olympus) equipped with a Retiga 4000R cooled Mono CCD camera (QImaging), a Spectra-X light engine illumination system (Lumencor), an IX2-ZDC laser-based autofocusing system (Olympus), a UPLXAPO 60X NA1.42 oil iris objective lens (Olympus), a MAC5000 controller for filter wheels and XY stage (Ludl Electronic Products), an incubation chamber (Tokai Hit), and a GM-4000 CO2 supplier (Tokai Hit). The following filters and dichroic mirrors were used: for FRET, an FF01-438/24 excitation filter (Semrock), an XF2034 455DRLP dichroic mirror (Omega Optical), an FF01-542/27 emission filter (Semrock), 10% intensity of Blue light in the Spectra-X light engine illumination system, 100 msec exposure time; for mTFP, an FF01-438/24 excitation filter (Semrock), an XF2034 455DRLP dichroic mirror (Omega Optical), and an FF01-483/32 emission filter (Semrock), 10% intensity of Blue light in the illumination system, 100 msec exposure time; for mVenus, an FF01-475/28 excitation filter (Semrock), an XF2034 455DRLP dichroic mirror (Omega Optical), an FF01-542/27 emission filter (Semrock), 100% intensity of Cyan light in the illumination system, 100 msec exposure time. Camera binning is 2x2, and images were obtained every 3 sec for 50 min. The microscopes were controlled by MetaMorph software (Molecular Devices).

**Measurement noise estimation**

For each cell, three background-subtracted fluorescence time series $\mathcal{D} = \{I_{AA,1:N_A}, I_{DD,1:N_D}, I_{DA,1:N_D}\}$ were obtained by manually selecting ROIs (regions of interest) that enclose the cell and subtracting background signals. The magnitude of measurement noise time series at each time point was estimated assuming that there is a large time-scale separation between measurement noise and temporal changes in fluorescence intensity due to other reasons (e.g., photobleaching and biological signals). Specifically, we first segmented each time series into $n_{seg}$ (= 8) non-overlapping and equal-length segments. Then, to estimate the magnitude of measurement noise at each segment, we computed, at each segment, the standard deviation ($s_i$) and its 95% confidence interval ($\sigma_i$; using the bootstrap method) of the difference between a raw and smoothed (20-th order median filtering) time series. A linear function $f(t, \boldsymbol{\theta}_m) = \theta_{m,1} + \theta_{m,2}t$ was fitted by maximizing the log-likelihood function:

$$\boldsymbol{\theta}_{m,opt} = \underset{\boldsymbol{\theta}_m}{\operatorname{argmax}} \sum_{i=1}^{n_{seg}} -\frac{(f(t_i, \boldsymbol{\theta}_m) - s_i)^2}{2\sigma_i^2},$$

where $\{t_i\}_{i=1}^{n_{seg}}$ are the midpoints of the segments. The magnitude (or the standard deviation) of measurement noise at time $t$ was then estimated to be $f(t, \boldsymbol{\theta}_{m,opt})$.

**References**


1. Ponsioen, B. *et al.* Detecting cAMP-induced Epac activation by fluorescence resonance energy transfer: Epac as a novel cAMP indicator. *EMBO Rep.* **5**, 1176–1180 (2004).
2. Akagi, T., Shishido, T., Murata, K. & Hanafusa, H. v-Crk activates the phosphoinositide 3-kinase/AKT pathway in transformation. *Proc. Natl. Acad. Sci.* **97**, 7290–7295 (2000).

3. Hitoshi, N., Ken-ichi, Y. & Jun-ichi, M. Efficient selection for high-expression transfectants with a novel eukaryotic vector. *Gene* **108**, 193–199 (1991).

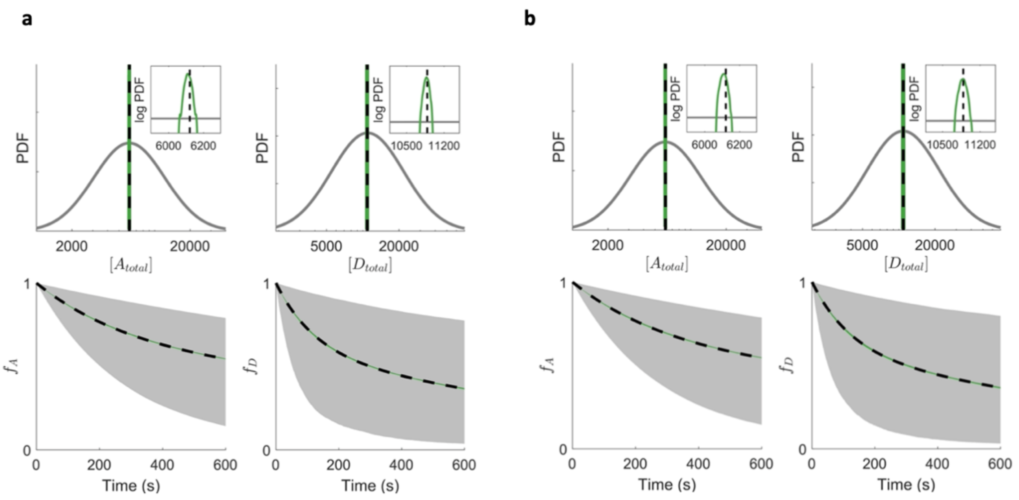


**Supplementary Figure 1** Prior and posterior distributions of the model parameters for the random and step FRET data shown in Fig. 2a.

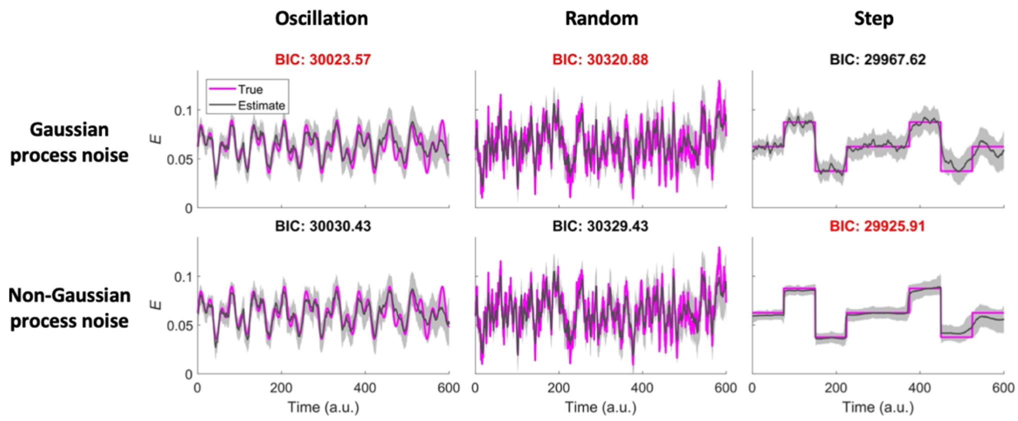


**Supplementary Figure 2** B-FRET, combined with model selection, automatically selects a model best evidenced by data. Three synthetic FRET data sets with oscillatory (left), random (middle) and step-like (right) FRET dynamics were analyzed by B-FRET, assuming Gaussian (top) and Non-Gaussian (bottom) process noise (Online Methods). True signal (magenta), estimated signal (grey), and 95% credible intervals (grey shade) are shown. For Non-Gaussian process noise, we used the Student's t-distribution, which has one more parameter and contains Gaussian distribution as a special case (Online Methods). For the oscillation and random data, the Bayesian information criterion (BIC) selects Gaussian process noise, implying the extra parameter of the Student's t-distribution does not contribute to inferring the FRET signals but only increases the complexity of the model. On the other hand, for the step data, the BIC selects the Non-Gaussian model because it captures the abrupt changes in the FRET signal while the Gaussian model fails to do so. Lower BIC values are highlighted by red.

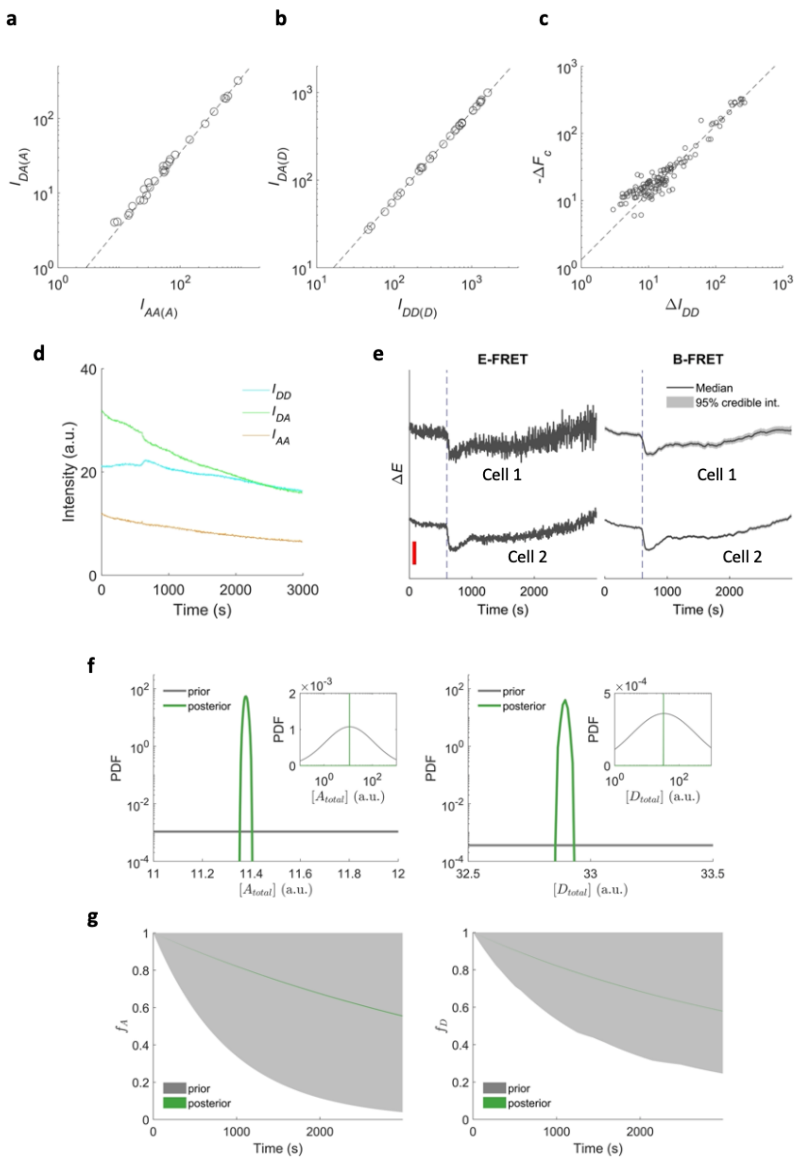


**Supplementary Figure 3** B-FRET can be applied to unimolecular FRET systems. (**a**) Single-cell fluorescence intensities from a strain that only express the acceptor (mVenus), obtained through the acceptor channel $I_{AA(A)}$ and the FRET channel $I_{DA(A)}$. The slope gives an estimate of the cross-excitation coefficient $a = 0.346$ $(0.331 - 0.357$; 95% confidence interval). (**b**) Single-cell fluorescence intensities from a strain that only express the donor (mTFP), obtained through the donor channel $I_{DD(D)}$ and the FRET channel $I_{DA(D)}$. The slope gives an estimate of the bleedthrough coefficient $d = 0.606\ (0.601\ - 0.609)$. (**c**) Changes in the donor fluorescence signal $\Delta I_{DD}$ and the negative change in the sensitized emission $-\Delta F_C$ before and after a stimulus (50 uM Forskolin and 100 uM IBMX ) application, obtained from a strain that expresses the unimolecular FRET probe harboring the donor and acceptor. The slope gives the parameter $G = 1.316\ (1.253\ - 1.396)$. (**d**) Background-subtracted fluorescence time series obtained from a single-cell. (**e**) Two representative time series of FRET index $E$ estimated by E-FRET (left) and B-FRET (right). The red bar is $\Delta E = 0.05$. The time at which 50 µM Forskolin and 100 µM IBMX were applied is indicated by the blue dashed lines. For B-FRET, the median and 95% credible interval are shown. The data shown in **d**, **f**, and **g** corresponds to the bottom data in **b**. This demonstrates that B-FRET also improves the SNR for a unimolecular FRET system. (**f**) The priors and posteriors of the total acceptor ($[A_{total}]$) and donor ($[D_{total}]$) concentrations. The same plots are shown in the insets with wider x ranges. (**g**) The ranges of priors and posteriors (0.1 - 99.9 percentile) of the intact fraction of the acceptor $f_A$ (left) and the donor $f_D$ (right) according to the prior (grey) and posterior (green) distributions are shown.

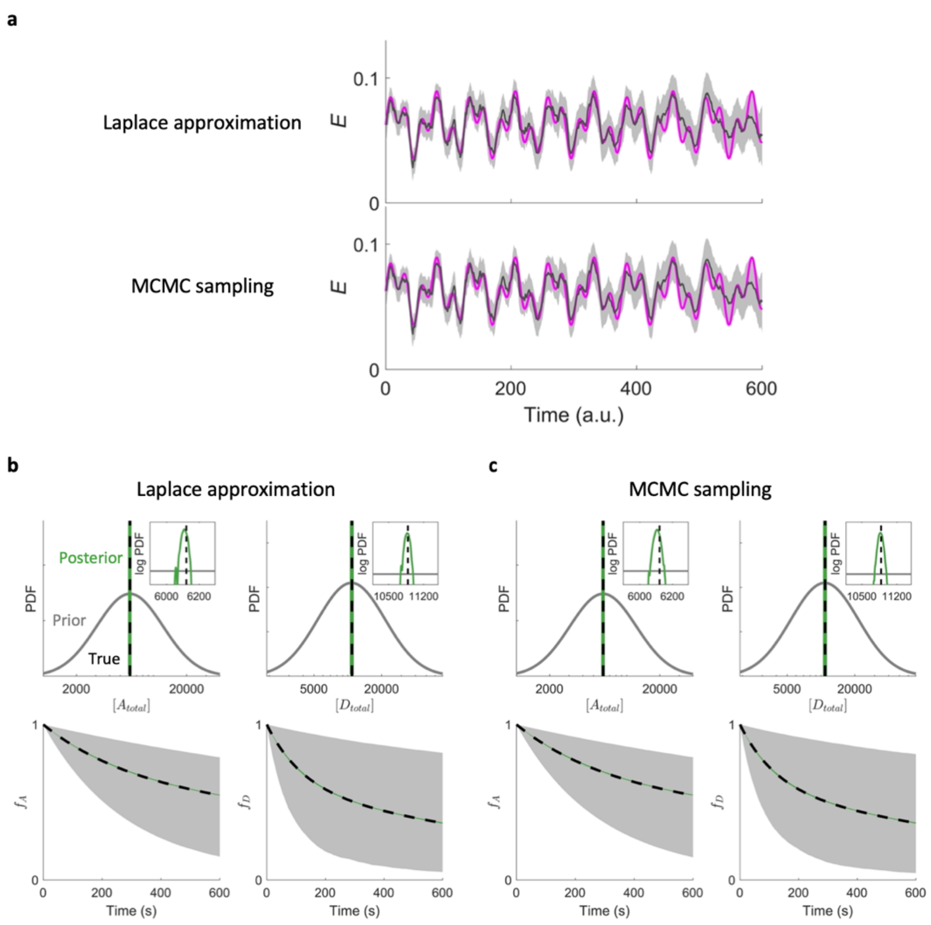


**Supplementary Figure 4** Gaussian approximation of the posterior distributions of the model parameters reduces the computational cost without affecting the results significantly. (**a**) The exact evaluation of the posterior distribution $p(\boldsymbol{\theta}|\mathcal{D}, \mathcal{M})$ by a Markov chain Monte Carlo (MCMC) method is computationally costly, so we approximated the posterior distribution by a log-normal distribution (Laplace approximation) by computing the Hessian matrix at the mode of the distribution (see Supplementary Note 2). The extracted FRET index $E$ by B-FRET with (top) and without (bottom) the approximation are shown. The difference between the two method is practically negligible, validating the usage of the approximation. (**b**) True values (black) and prior(grey) and posterior (green) distributions of the parameters obtained by the Laplace approximation are shown (same as Fig. 1c). (**c**) Same as **b** except the posterior distributions (green) are obtained by a MCMC sampling method.

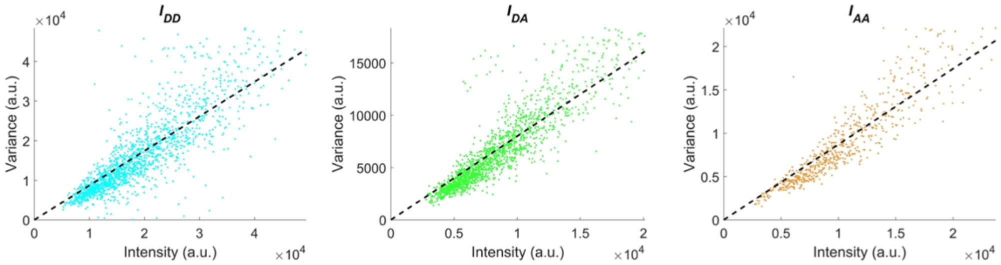


**Supplementary Figure 5** Determining the relationship between the expected value of a fluorescence intensity and the variance of measurement noise. When the measurement noise is dominated by shot noise, the variance of shot noise is proportional to fluorescence intensity (Supplementary Note 3). The proportionality constant can be estimated by plotting the variance of shot noise, which is estimated from the autocorrelation function of a segment of fluorescence time series, against the average fluorescence intensity, and by computing the slope. For the setup used in this paper, the slopes are: 0.872 (0.856 - 0.886) for $I_{DD}$, 0.801 (0.789 - 0.811) for $I_{DA}$, and 0.872 (0.854 - 0.889) for $I_{AA}$, where 95% bootstrap confidence intervals are shown in parentheses.
